# Beyond Visual Inspection: Principled Benchmarking of Single-Cell Trajectory Representations with scTRAM

**DOI:** 10.1101/2025.06.23.661141

**Authors:** Kemal Inecik, Antony Rose, Muzlifah Haniffa, Malte Luecken, Fabian J. Theis

## Abstract

Single-cell representation learning aims to construct low-dimensional embeddings that preserve biologically salient manifold structures. Yet current evaluation protocols remain inadequate for assessing trajectory fidelity—essential for studying cell differentiation, immune responses, disease progression and perturbation—relying on heuristic visualizations or scalar surrogates that overlook multiscale trajectory geometry. To address this gap, we present *scTRAM*—single-cell trajectory representation metrics—a principled benchmarking framework that quantitatively evaluates how well single-cell embeddings retain groundtruth trajectories across complementary failure modes, from local neighborhood scrambling to global branch mis-ordering. Through our experiments, we show that *scTRAM* reveals model-specific trade-offs in preserving distinct aspects of trajectory structure, with edge-specific decomposition exposing localized representational biases at high resolution. Systematic validation across diverse biological contexts confirms that the metric suite is sensitive to genuine structural properties rather than statistical artifacts. By shifting from qualitative, ad-hoc judgments to quantitative, principled approach, *scTRAM* enables rigorous comparison of embedding methods and informs downstream analyses where trajectory integrity is essential for biological interpretation.

## 1. Introduction

Single-cell omics technologies now profile millions of cells across diverse tissues, perturbations, and timepoints, revealing high-dimensional manifolds of gene-expression states—yet the underlying structure is obscured by technical noise and sampling sparsity (Inecik et al., 2024; Cui et al., 2025). Modern representation learning pipelines—from flow matching to deep variational and single cell transformer generative models—thus aim to embed these data into lowdimensional latent representations that enable scalable inference, visualization, and hypothesis generation while retaining the geometry of biologically salient processes such as cell differentiation, tissue development, and disease progression (Inecik & Theis, 2023; Schaar et al., 2024; Klein et al., 2025). For developmental, immunological, and pathological systems alike, an important aspect of the manifold’s most biologically critical features is cell-state trajectories: continuous paths and branch points that trace lineage commitment, dose–response dynamics, or disease progression over pseudotime (Saelens et al., 2019). However, current evaluation protocols remain ad-hoc, emphasizing cluster purity or reconstruction error rather than trajectory fidelity; they fail to assess whether embeddings preserve geodesic continuity, respect branch topology, and maintain pseudotemporal ordering consistent with lineage tracing or perturbation timestamps (Luecken et al., 2022; Hu et al., 2024; Song et al., 2023). While qualitative visual inspection of trajectories— such as plotting marker genes along pseudotime or overlaying branch points on UMAPs—is routinely employed in single-cell atlassing studies, these approaches lack reproducibility, scalability, and quantitative rigor, as they depend heavily on subjective annotations, heuristic parameter tuning, and small-scale validation that cannot generalize across datasets or models (Chari & Pachter, 2023; Heumos et al., 2023). The absence of standardized, principled, and quantitative benchmarks impedes rigorous model selection and hampers biological interpretation, motivating the need for a framework grounded in differential-geometric distortion measures and temporal alignment theory to systematically score embeddings based on their conservation of the intrinsic manifold structure underlying cellular transitions.

Quantitatively evaluating whether an embedding respects a ground-truth cellular trajectory entails comparing two metric–topological objects: the intrinsic trajectory graph (*G*, *d* _*G*_), derived from lineage labels or time-course experiments, and its image under an embedding *f* inside a latent space (*Y*, *d*_*Y*_). The embedding *f* must approximate a branch-preserving, arc-length-monotone homeomorphism— retaining not only topological connectivity but also the relative progression rates along paths—up to bounded distortion. Fidelity assessment must address two interrelated scales: locally, where violations of Lipschitz continuity scramble neighborhood context and obscure short-range lineage bifurcations, and globally, where seemingly minor edgewise distortions reorder branch points, alter Betti numbers, or flatten long-range pseudotemporal gradients (Haghverdi et al., 2015; Munkres, 2000). Formalizing this comparison as an optimization problem—whether via optimal transport on branched manifolds or minimum-cost graph-edit operations—faces inherent NP-hardness barriers (Chen et al., 2018; Qu et al., 2025), while existing surrogate measures—e.g., geodesic stress, trustworthiness, or local continuity—illuminate only one failure mode at a time (Najim, 2015; Vathy-Fogarassy & Abonyi, 2009). Topological persistence diagrams detect structural rearrangements yet remain indifferent to arc-length monotonicity or pseudo-time alignment, while manifold-learning surrogates such as Isomap or diffusion-based kernels prioritize distance preservation over topological consistency (Qu & Cai, 2017). These intertwined limitations make it impossible to declare victory with a lone scalar: rigorous evaluation demands a battery of complementary, computationally tractable metrics that decompose trajectory fidelity into topology preservation, geodesic continuity, and pseudotime monotonicity, thereby exposing trade-offs that any representation unavoidably incurs.

To address this critical gap, we present *scTRAM* (single-<u>c</u>ell <u>TRA</u>jectory representation <u>M</u>etrics): a systematically designed benchmarking framework that offers a cohesive suite of complementary metrics—spanning topological consistency, manifold continuity, and pseudotime alignment—to evaluate how effectively a representation preserves user-specified ground-truth trajectories. By decomposing trajectory integrity into distinct axes of performance rather than relying on a single scalar, *scTRAM* pinpoints where an embedding may fail to maintain local continuity, respect branched structures, or preserve correct temporal progression, while remaining user-friendly through an end-to-end pipeline. This decomposed, multi-axis approach enables fine-grained evaluation of trajectory fidelity across a range of biologically and computationally relevant scenarios: for instance, comparing trajectories across conditions (e.g., wild-type versus CRISPR knockout differentiation paths), benchmarking trajectory representations from competing generative or dimensionality reduction approaches, evaluating data integration processes with respect to trajectory conservation (e.g., assessing whether a single-cell atlas retains information present in its component datasets). In this study, we *(i)* perform comprehensive benchmarking of multiple trajectory representations across diverse models using a large-scale single-cell dataset, *(ii)* demonstrate how failure mode analysis of trajectory representations informs downstream data analysis decisions, and *(iii)* provide rigorous benchmarking experiments validating the robustness and discriminative power of the *scTRAM* metric suite. Consequently, *scTRAM* helps shift trajectory-representation evaluation from largely qualitative, ad-hoc judgments toward a more principled approach grounded in multiscale, structure-aware benchmarks. ^1^

## 2. Methods

*scTRAM* evaluates trajectory fidelity by comparing a lowdimensional representation against a ground-truth reference trajectory, formalized as a directed graph *G*_ref_ = (*V*, ℰ) with vertices *V* (cell types) and edges (state transitions). Given a user-provided representation *Y* ∈ ℝ^*N* ×*e*^ (N cells, e dimensions) and _ref_, *scTRAM* preprocesses both into intermediate representations that enable trajectory-aware metric computation (Figure 1). First, cells are subset to *Y*_sub_ ∈ ℝ^*n*×*e*^ (*n* ≤*N*), retaining only those annotated with cell types in *G*_ref_. For *Y*_sub_, a k-nearest-neighbor (kNN) graph is constructed and its edge weights are converted into a row-stochastic diffusion operator that represents transition probabilities between cells. By diagonalizing this operator, diffusion components are extracted and the pseudotime vector *p*_inf_ ∈ ℝ^*n*^ is defined as the diffusion distance from a chosen root cell, in a manner analogous to scanpy’s DPT method (Wolf et al., 2018). Concurrently, connectivity at the cell-type level is summarised by pooling kNN edges between every pair of annotated types and normalizing the resulting counts by the size of the smaller cluster, yielding a weighted inter-type matrix *W*_inf_ ∈ ℝ^*t*×*t*^ (*t* cell types) in a procedure analogous to scanpy’s PAGA method (Wolf et al., 2019). A custom significance threshold is then applied to *W*_inf_ to obtain a complementary binary adjacency matrix *A*_inf_ ∈ {0, 1} ^*t*×*t*^; both are kept and used by different benchmarking metrics as needed.

**Figure 1.**
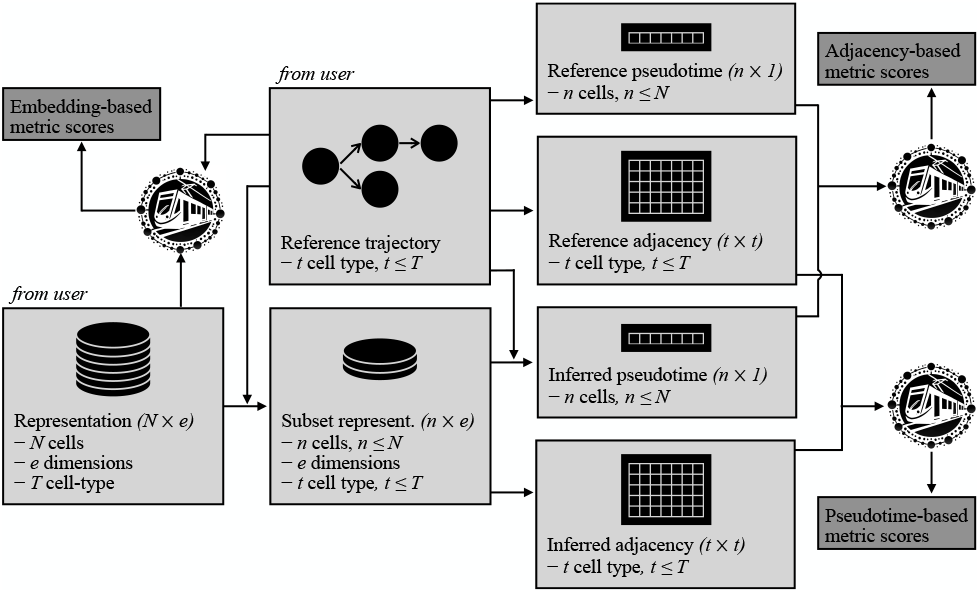
Overview of *scTRAM* benchmarking workflow. From an input embedding *Y* and reference trajectory graph *G*_ref_, *scTRAM* (i)restricts to cells present in *G*_ref_ producing *Y*_sub_, *(ii)* computes reference and inferred pseudotimes (*p*_ref_, *p*_inf_) and inter-type connectivities (*A*_ref_, *Z*_inf_), and *(iii)* assembles embedding-, adjacency-, and pseudotime-based metrics whose values form the score vector *s ∈* ℝ ^*M*^.

Meanwhile, *G*_ref_ is mapped to a symmetric adjacency matrix *A*_ref_ *∈* {0, 1}^*t*×*t*^ together with a type-level pseudotime vector 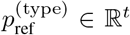. Rather than relying on shortest-path distances, 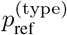 is derived by a damped diffusion with restart procedure. Let *D* = diag(*A*_ref_1_*t*_) denote the degree matrix and define the row-stochastic random-walk operator *P* = *D*^*−*1^*A*_ref_. Starting from the one-hot vector *e*_*r*_ that marks the user-specified root cell (inferred by *G*_ref_), we iterate *F* ^(*k*+1)^ = *αP* ^⊤^*F* ^(*k*)^ + (1−*α*)*e*_*r*_ with damping factor *α* ∈ (0, 1) until ∥*F* ^(*k*+1)^ −*F* ^(*k*)^ _1_ ∥*< ε*, where *ε >* 0 is a prescribed convergence tolerance, finally setting 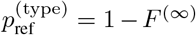. The steady-state *F* ^(∞)^ gives the visitation probability of each type under an *α*-resummed random walk that constantly restarts at the root, so values smoothly increase with topological distance from the origin. Finally, 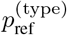 is broadcast to the cell level by assigning every cell the pseudotime of its annotated type, yielding *p*_ref_ ∈ ℝ^*n*^ that is compatible with all downstream cell-resolved metrics. To minimize redundant computation, *scTRAM* performs graph construction, diffusion smoothing, and binarization with sparse-matrix routines while caching salient intermediates— such as KNN indices, diffusion maps, and centroids—along with the objects (e.g. *W*_inf_, *A*_ref_, *A*_inf_), so that subsequent metric evaluations incur negligible overhead.

During evaluation, *scTRAM* assembles a set of *M* deterministic functionals 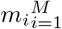, each instantiated as a mapping *m*_*i*_(*O* _ref_, *O* _inf_) →*s*_*i*_ ∈ ℝ where the ordered pair of operands (*O*_ref_, *O*_inf_) is drawn from exactly one of three dimensionality-compatible domains: *(i)* the pseudotime domain (*p*_ref_, *p*_inf_) ∈ ℝ^*n*^× ℝ^*n*^, yielding *Group 1* scores that probe temporal monotonicity via, e.g., Spearman’s *ρ*, dynamic time warping distance, and the concordance index; *(ii)* the connectivity domain (*A*_ref_, *Z*_inf_) ∈ {0, 1} ^*t*×*t*^ ×ℝ^*t*×*t*^, where *Z*_inf_ ∈ *A*_inf_, *W*_inf_ allows metrics to operate on either the binarized or weighted inferred graph and furnishes *Group 2* scores that evaluate topological overlap through metrics such as persistence diagram distance, GNN-based cosine similarity with GIN architecture, and Wasserstein distance between the normalized Laplacian eigenvalue spectra; and *(iii)* the embedding–trajectory domain (*A*_ref_, *Y*_sub_) ∈ {0, 1} ^*t*×*t*^ ×ℝ^*n*×*e*^, producing *Group 3* scores that compare manifold geometry using, for instance, Sammon’s stress, graph-based trustworthiness, and directionality preservation (refer to Supplementary D.3 for comprehensive descriptions of the *scTRAM* metrics). The resulting scalars *s*_*i*_ are concatenated into the score vector *s* = (*s*_1_, …, *s*_*M*_), which is returned for downstream aggregation and reporting.

## 3. Results

### 3.1. *scTRAM* enables comprehensive quantitative evaluation of trajectory representation fidelity in single-cell embeddings

To demonstrate *scTRAM*’s utility in quantitatively assessing trajectory representation fidelity across data integration methods, we conducted a comprehensive benchmarking experiment using *Suo* dataset (Suo et al., 2022)—a highly heterogeneous large-scale single-cell dataset spanning multiple organs and developmental time points—with particular focus on the trajectories of hematopoietic lineage development. While data integration benchmarking tools like scIB (Luecken et al., 2022) effectively quantify batch correction and cluster alignment, they fail to capture whether embeddings preserve the differential-geometric properties of continuous cellular transitions that underpin lineage commitment and differentiation dynamics. Such preservation is crucial for developmental biology, where accurate representation of transition paths directly determines the validity of downstream analyses such as lineage bifurcation timing, transcription factor activation sequencing, and gene-regulatory network inference—analyses that collapse when trajectory topology is distorted.

To ensure comprehensive evaluation across diverse developmental contexts and to counteract the risk that a single dominant path biases overall metrics, we partitioned the hematopoietic lineage into 27 biologically meaningful trajectory segments spanning four functional groups: myeloid, lymphoid, MEM (megakaryocyte-erythroid-mast lineage), and stem. These segments encompassed both global structures (e.g., the entire hematopoietic lineage or the complete myeloid trajectory) as well as fine-grained branches (e.g. T-cell maturation), ensuring balanced coverage of distinct developmental pathways. As illustrated in Figure 2 (left), our rank-based performance analysis—where each metric for each trajectory was ranked across trained models to normalize scale differences—revealed that scANVI demonstrated superior overall trajectory representation performance (median rank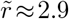), followed closely by TarDis 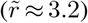. This performance hierarchy contrasts with conventional scIB rankings (Table 1), where TarDis performed marginally better on data integration metrics. This discrepancy underscores how *scTRAM* complements scIB’s bio-conservation scores by specifically evaluating trajectory preservation, offering a more complete framework for data integration quality assessment in developmental contexts. To align with scIB conventions, we also reported inverse average ranks (higher is better) in Table 1, revealing that optimal representation selection depends critically on whether trajectory fidelity or batch correction is the primary analytical concern—a nuance previously unavailable to practitioners relying solely on conventional integration benchmarks. Although we have not yet established a rigorous weighting strategy to combine these aggregates, we contend that careful attention to trajectory fidelity is crucial for a balanced assessment of data integration performance.

**Figure 2.**
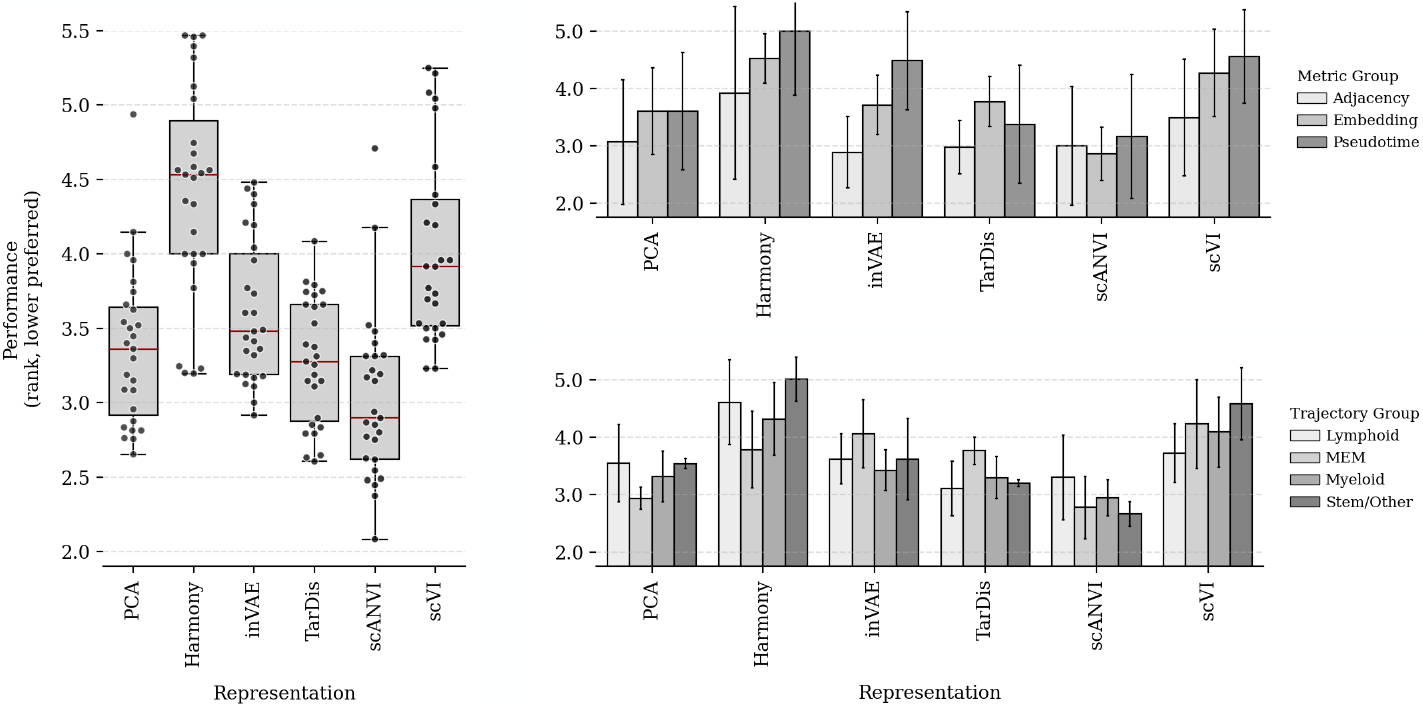
Comparative trajectory fidelity of six data integration models on the prenatal *Suo* dataset. Left, box and scatter plots show the rank distributions (lower is better) of 27 biologically curated trajectory segments that tile the hematopoietic manifold; each dot is a segment, red bars denote median rank per model, and the outer box conveys dispersion. Top-right, mean (±s.e.m.) ranks within the three *scTRAM* metric groups, revealing model-specific strengths in preserving different aspects of trajectory structure. Bottom-right, mean (± s.e.m.) rank after grouping trajectory segments by major lineage class. The decomposition highlights failure modes that are invisible to a single aggregate score but critical for downstream inference.

**Table 1.** Joint benchmarking with scIB data integration scores and the *scTRAM*’s trajectory representation aggregate (inverse average rank) across six model representations. Columns report bio conservation *(Bio cons*.), batch-effect removal *(Batch corr*.), and their composite *(Total scIB)* from scIB, followed by the mean trajectory-representation score *(Traj. repr*.). Refer to Supplementary D.2 for descriptions of the scIB suite and C.2 for model details.

|  | Bio cons. | Batch corr. | Total scIB | Traj. repr. |
| --- | --- | --- | --- | --- |
| PCA | 0.606 | 0.402 | 0.524 | 0.297 |
| Harmony | 0.569 | 0.492 | 0.538 | 0.226 |
| inVAE | 0.629 | 0.382 | 0.530 | 0.277 |
| TarDis | 0.643 | **0.627** | **0.637** | 0.303 |
| scANVI | **0.701** | 0.523 | 0.629 | **0.332** |
| scVI | 0.598 | 0.603 | 0.600 | 0.248 |

### 3.2. Trajectory-aware benchmarking reveals model-specific trade-offs in hematopoietic lineage representations

This finding of model-specific strengths motivated us to further dissect performance patterns across our metric taxonomy and biological lineage groups, revealing fundamental trade-offs inherent in representation learning approaches. The hierarchical decomposition of the 27 trajectory segments described earlier enabled evaluation across multiple biological scales, providing a multi-resolution view of model performance across the hematopoietic landscape. Disaggregating performance by metric group revealed model-specific strengths in preserving different aspects of trajectory structure. inVAE exhibited superior performance in adjacency-based metrics (mean rank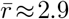), followed by TarDis 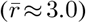, suggesting particular strength in preserving topological relationships between cell types. Conversely, scANVI excelled in embedding and pseudotime-based metrics (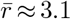and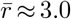, respectively), again with TarDis showing strong secondary performance (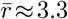and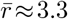, respectively), indicating better preservation of manifold geometry. This pattern reveals a critical trade-off in representation learning: models optimized for preserving topological connectivity may sacrifice fidelity in pseudotemporal ordering and vice versa—an insight crucial for selecting appropriate models based on downstream analytical priorities. Similarly, trajectory-specific analysis demonstrated that no single model universally outperformed across all lineage segments; while scANVI generally dominated in MEM 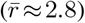, myeloid 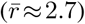, and stem/other groups 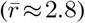, TarDis achieved superior representation of lymphoid trajectories (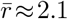TarDis; 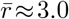 scANVI). This heterogeneity in performance across trajectory types indicates that model architectures exhibit subtle biases in handling branching complexity and local continuity, reflecting how different objective functions and regularization strategies balance different aspects of trajectory fidelity.

Our hierarchical segmentation of trajectory slices enabled multi-scale interrogation of performance across the hematopoietic landscape, yet such a manual approach introduces constraints on scalability and generalizability, and ease of use when examining complex, branching developmental systems. To address these limitations while maintaining analytical precision, we implemented an automated trajectory decomposition algorithm specifically optimized for single-cell trajectories that partitions arbitrary manifolds into constituent segments without requiring manual curation. ^2^ Application of this algorithm to the hematopoietic lineage graph yielded hundreds of trajectory segments, spanning diverse developmental transitions from stem cell activation to terminal differentiation. To evaluate model performance within this refined framework, we computed the mean rank performance for each edge in the full hematopoietic trajectory network, averaging across all applicable metrics and trajectory segments containing that edge, thereby deriving edge-specific fidelity scores for each model representation. This granular decomposition revealed localized performance disparities between representations that remained obscured in aggregate scoring schemes. As illustrated in Figure 3a and 3b, ^3^ scANVI and TarDis exhibited distinct edge-level performance signatures across the hematopoietic manifold, despite relatively comparable overall metrics discussed earlier. Specifically, scANVI more faithfully preserved early bifurcation events, stem cell transitions, and the myeloid lineage (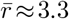TarDis; 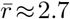 scANVI), whereas TarDis demonstrated superior performance in maintaining the lymphoid differentiation branch—particularly within B-cell differentiation trajectories (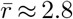TarDis; 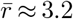scANVI)—which is consistent with earlier findings.

**Figure 3.**
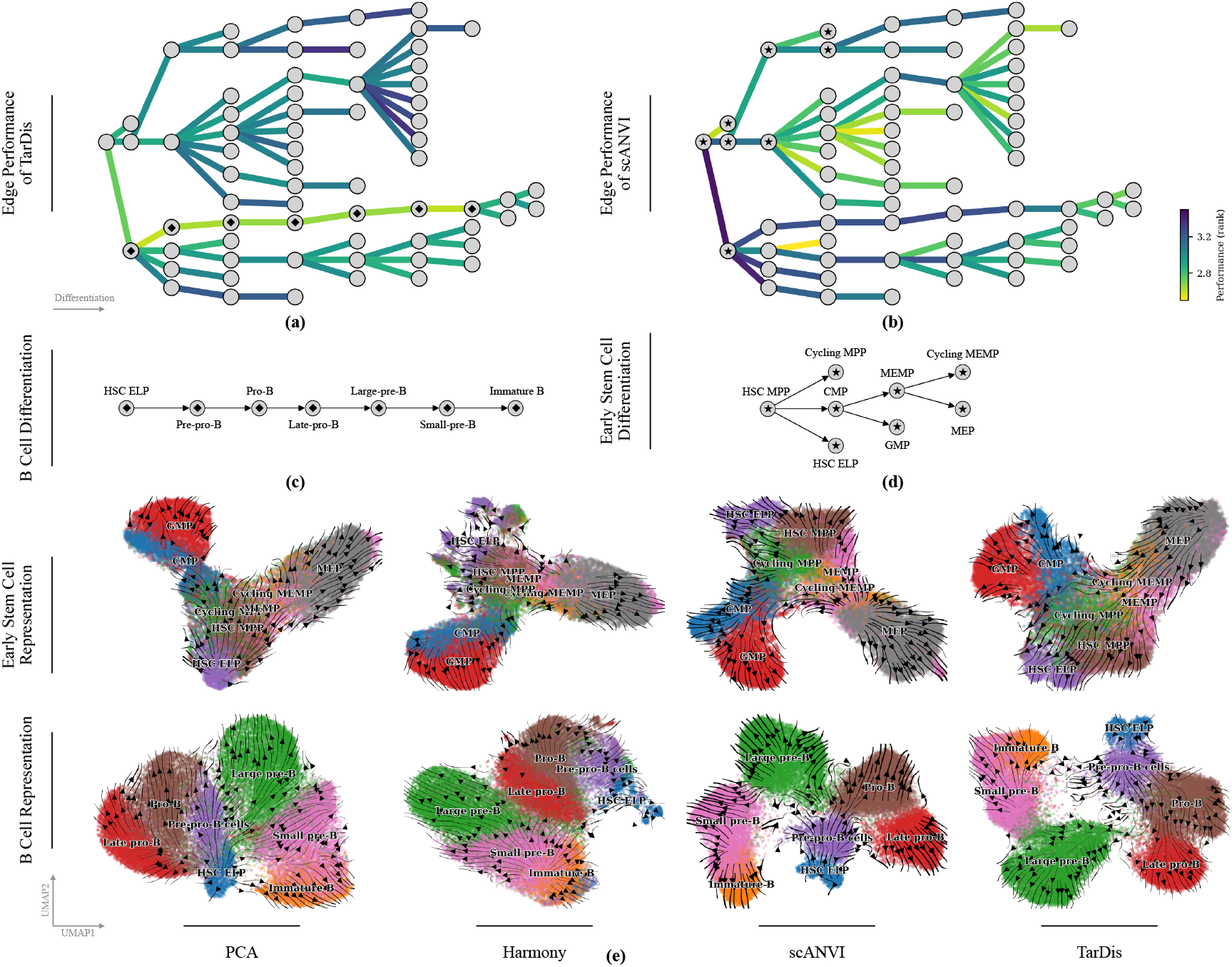
Trajectory edge-specific performance analysis revealing localized strengths and weaknesses across integration methods. Panels (a) and (b) depict the mean edge-level performance of TarDis and scANVI, respectively, across the full hematopoietic manifold, where each directed edge in the reference lineage graph is colored by its average performance rank. Panels (c) and (d) provide schematics of reference early stem-cell and B-cell differentiation trajectories, respectively, for visual correspondence with the embedding projections in panel (e). Panel (e) juxtaposes UMAP projections of the embeddings produced by PCA, Harmony, scANVI, and TarDis; points are coloured by annotated cell types, and black arrows denote the local expected displacement vectors computed with CellRank’s pseudotime-kernel (Weiler et al., 2024). TarDis preserves the B-cell manifold in the lower-right sector with minimal geometric distortion, whereas the edge-wise analysis of early progenitors reveals a mosaic fidelity pattern split across TarDis, scANVI, PCA, and Harmony.

The edge-specific performance decomposition revealed representational biases undetectable by neither conventional integration benchmarks nor aggregate trajectory metrics, establishing a principled basis for context-dependent model selection that prioritizes trajectory fidelity in regions of highest biological relevance. For instance, researchers focusing on B-cell development would benefit substantially from selecting TarDis despite its marginally lower overall trajectory representation score, as it outperforms scANVI in modeling B-cell lineage progression (Figure 3e, bottom). Conversely, investigators primarily concerned with myeloid differentiation would achieve superior results using scANVI despite its marginally lower integration scores. More strikingly, our decomposition demonstrated that even within specific developmental programs—such as stem cell differentiation as illustrated in Figure 3e, top—certain transition edges were better preserved by TarDis while others by scANVI, suggesting that optimal representation selection depends not merely on broad lineage categories but on specific developmental transitions under investigation. This heterogeneity in trajectory-level and edge-level performance fundamentally challenges the prevailing paradigm of representation selection based on aggregate benchmarks, demonstrating instead that model selection criteria should be trajectory-specific— or even edge-specific—when downstream analytical priorities are well-defined.

Beyond comparative model evaluation, we propose that *scTRAM* can serve as an optimization criterion during hyperparameter tuning, where edge-specific or trajectory-specific scores guide model selection during training to yield representations optimized precisely for the developmental transitions most relevant to biological inquiry, rather than for general-purpose integration metrics that may inadvertently sacrifice critical trajectory features. This approach transforms representation learning for developmental systems from generalized fitting to targeted preservation of biologically critical structures, enabling more biologically grounded embeddings for trajectory-focused downstream analyses such as pseudotemporal gene expression modeling, branching point characterization, and fate-decision mapping. We contend that this trajectory-aware optimization paradigm is not confined to developmental systems, but generalizes to disease-progression modeling—such as selecting representations that best preserve transcriptional transitions from premalignant to metastatic states—or perturbation-response settings, where treated cells span a continuous manifold across dosage gradients or temporal windows. Likewise, in studies of immune repertoire dynamics, where antigen-driven clonal expansion unfolds along bifurcating trajectories, accurate localization of branch points is essential. We regard *scTRAM* as a promising, general-purpose tool for such settings, offering a trajectory-aware lens that complements existing evaluation frameworks while remaining adaptable to diverse biological contexts.

### 3.3. Systematic validation of *scTRAM* framework confirms sensitivity to trajectory distortion in controlled experiments

To validate that our trajectory-representation metrics behave as theoretically expected across disparate biological settings, we first refined the *scTRAM* suite on fully synthetic singlecell datasets generated with a custom simulator—bundled with the package for reproducibility and extension—and qualitatively validated metric behavior on the experimental *Suo* dataset through manual inspection of trajectory segments and score patterns; results omitted as both analyses were conducted solely for internal validation. We therefore proceeded to rigorously evaluate whether the *scTRAM* metric suite satisfies fundamental expectations across three distinct experimental paradigms, beginning with the hypothesis that during the training of variational autoencoder architectures such as scVI and TarDis, the latent representation should progressively improve with respect to bio conservation and trajectory preservation metrics, while batch correction performance should remain relatively stable due to the maximal latent space mixing achieved at initialization. To test this hypothesis, we evaluated trajectory representation fidelity and data integration performance across intermediate training states by tracking both *scTRAM* and scIB scores on three representative settings—the neurogenesis dataset *La-Manno* (La Manno et al., 2018), the germline development dataset *Garcia-Alonso* (Garcia-Alonso et al., 2022), and the dose-response perturbation dataset *Srivatsan* (Srivatsan et al., 2020)—using scVI or TarDis architectures. Metric values were harmonized by *z*-scoring each metric across checkpoints and inverting the sign for those whose optimum lies at lower numerical values, thereby producing a directionally consistent aligned score that can be averaged across heterogeneous metrics without bias. As anticipated, biological and trajectory conservation metrics exhibited overall improvement throughout training, with particularly pronounced gains during early epochs followed by asymptotic convergence (Figure 4). Simultaneously, batch correction metrics remained consistently high from initialization through convergence, confirming that the stochastic initialization procedure effectively induces latent space mixing that is subsequently preserved during optimization. These findings not only validate the sensitivity of *scTRAM* to genuine improvements in representation quality but also suggest that, at least within the scVI and TarDis training regimes, bio conservation and batch correction objectives are not inherently antagonistic—challenging the prevailing view that these goals necessarily represent conflicting constraints in representation learning.

**Figure 4.**
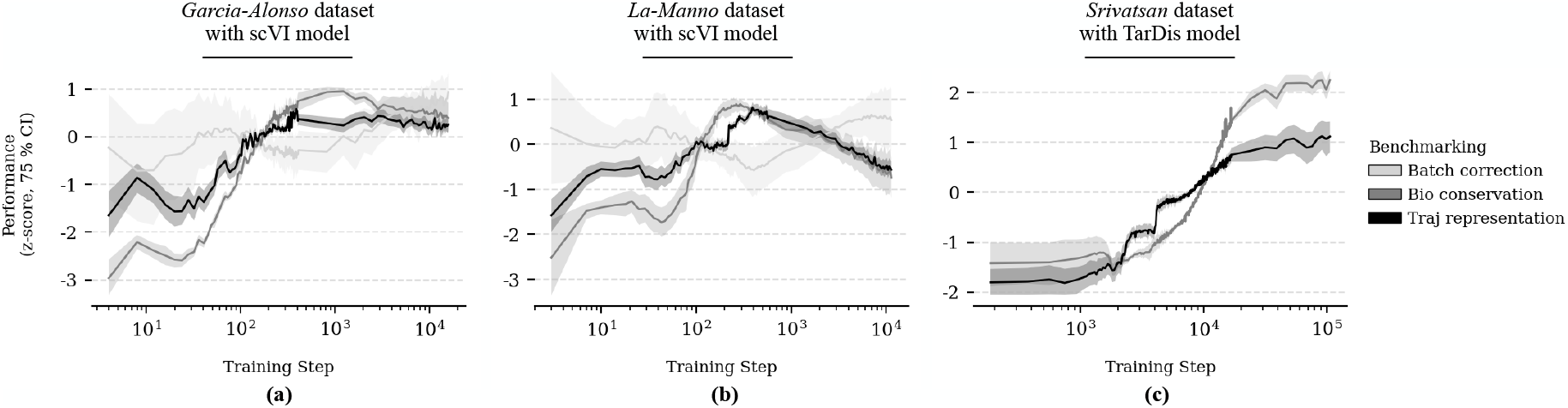
Training dynamics of trajectory representation metrics across distinct biological systems. *z*-score normalized performance of *scTRAM* metrics across training epochs for (a) the *Garcia-Alonso* germline development dataset using scVI, (b) the *La-Manno* neurogenesis dataset using scVI, and (c) the *Srivatsan* dose-response perturbation dataset using TarDis. Metrics are grouped into three categories: batch correction (light gray), bio conservation (medium gray), and trajectory representation (black). Shaded regions represent 75% confidence intervals across multiple metrics within each category.

Our second validation focused on the differential impact of cell-type ablation on trajectory representation fidelity. We postulated that selective removal of cells belonging to a trajectory-constituting cell type should progressively deteriorate trajectory representation quality, whereas removing cells from trajectory-extraneous populations should minimally affect trajectory metrics. To systematically investigate this relationship, we conducted ablation experiments on the *La-Manno* dataset, wherein we incrementally removed increasing percentages of cells from either trajectory-integral or trajectory-peripheral cell types (Figure 5a). Consistent with our hypothesis, *scTRAM* metrics exhibited monotonic degradation when trajectory-constituting cells were removed, with representation quality declining in direct proportion to ablation severity (Figure 5c). Conversely, when equivalent proportions of cells were ablated from trajectory-irrelevant populations, trajectory-representation scores remained remarkably stable, demonstrating robust invariance to perturbations that spare the reference lineage graph and leave the principal manifold intact. We further extended this analysis to evaluate the impact of feature (gene) selection through a systematic feature ablation experiment, showing that trajectory representation fidelity similarly scales with the proportion of informative genes retained (Figure 5b), a finding also with significant implications for feature selection strategies for trajectory representation fidelity. The differential response of *scTRAM* metrics to these precisely controlled perturbations provides important evidence that our framework captures genuine structural properties of trajectory representations rather than statistical artifacts or embedding-specific biases.

**Figure 5.**
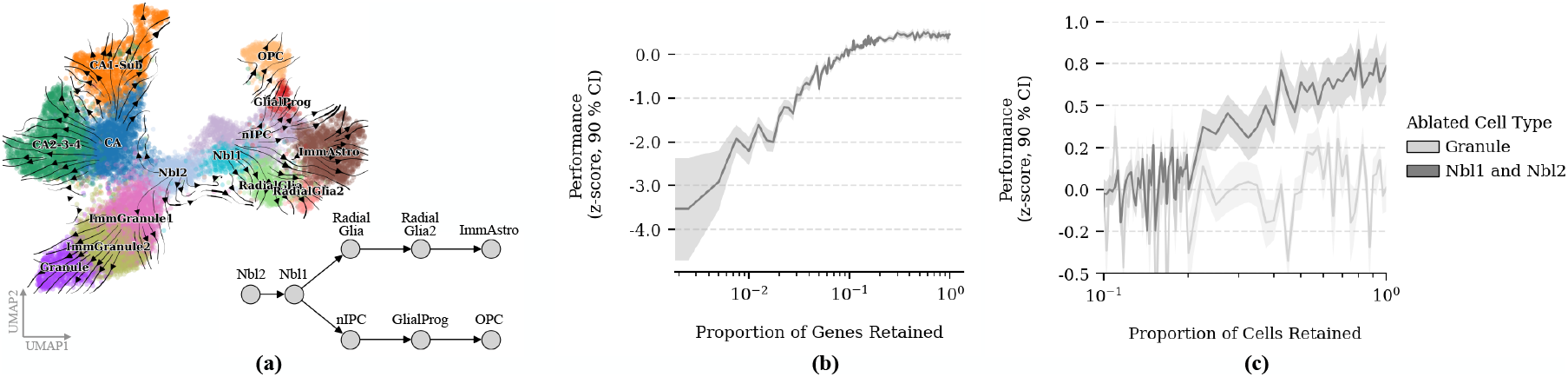
Systematic ablation experiments for validation of the *scTRAM* sensitivity to trajectory-specific perturbations. (a) UMAP visualization of the La-Manno neurogenesis dataset showing cell-type clusters and the reference trajectory graph with edges connecting developmentally related cell types. (b) Impact of feature ablation on trajectory representation fidelity, showing monotonic degradation of *scTRAM* performance (*z*-scored with 90% CI) as the proportion of informative genes retained decreases. (c) Differential sensitivity of *scTRAM* metrics to cell-type ablation, demonstrating robust preservation of scores when trajectory-irrelevant granule cells are removed (light gray) versus significant degradation when trajectory-integral Nbl1 and Nbl2 neuroblast populations are ablated (dark gray).

Finally, we investigated the relationship between disentanglement strength and trajectory representation in the TarDis architecture. TarDis, a variational autoencoder designed for disentanglement tasks, employs specialized loss functions that, when properly configured, guide the model toward local minima where desired trajectory representations are optimally preserved. We specifically applied TarDis disentanglement to drug and dose covariates, conditioning dose as a continuous axis, consistent with the original implementation. Our hypothesis posited that increasing the relative weight of disentanglement losses would progressively enhance covariate-specific representations and continuous dose-axis formation, thereby yielding more faithful trajectory preservation within the specialized latent subspaces. To evaluate this relationship, we conducted a comprehensive parameter sweep, training multiple TarDis models with incrementally increasing disentanglement weights while keeping all other hyperparameters constant. For each trained model, we extracted the dose-specific latent dimensions and applied *scTRAM* to quantify the fidelity of dose-response trajectories within this specialized subspace. The results confirmed our expectations: trajectory fidelity within the dose-specific latent subspace improved monotonically with increasing disentanglement weight, demonstrating that stronger covariate-specific regularization yields more coherent representations of continuous biological processes (Figure 6a, b; e, top row). Notably, we extended this analysis to the complete latent space of TarDis—comprising the union of dose-specific, drug-specific, and unreserved latent dimensions—revealing that *scTRAM* effectively quantifies trajectory preservation even in complex, compositional latent spaces where multiple biological processes are simultaneously encoded (Figure 6c, d; e, bottom row). Collectively, these systematic validation experiments confirm that *scTRAM* metrics respond to perturbations in ways that align with theoretical expectations across diverse contexts, reinforcing their utility as principled quantitative measures of trajectory representation quality.

**Figure 6.**
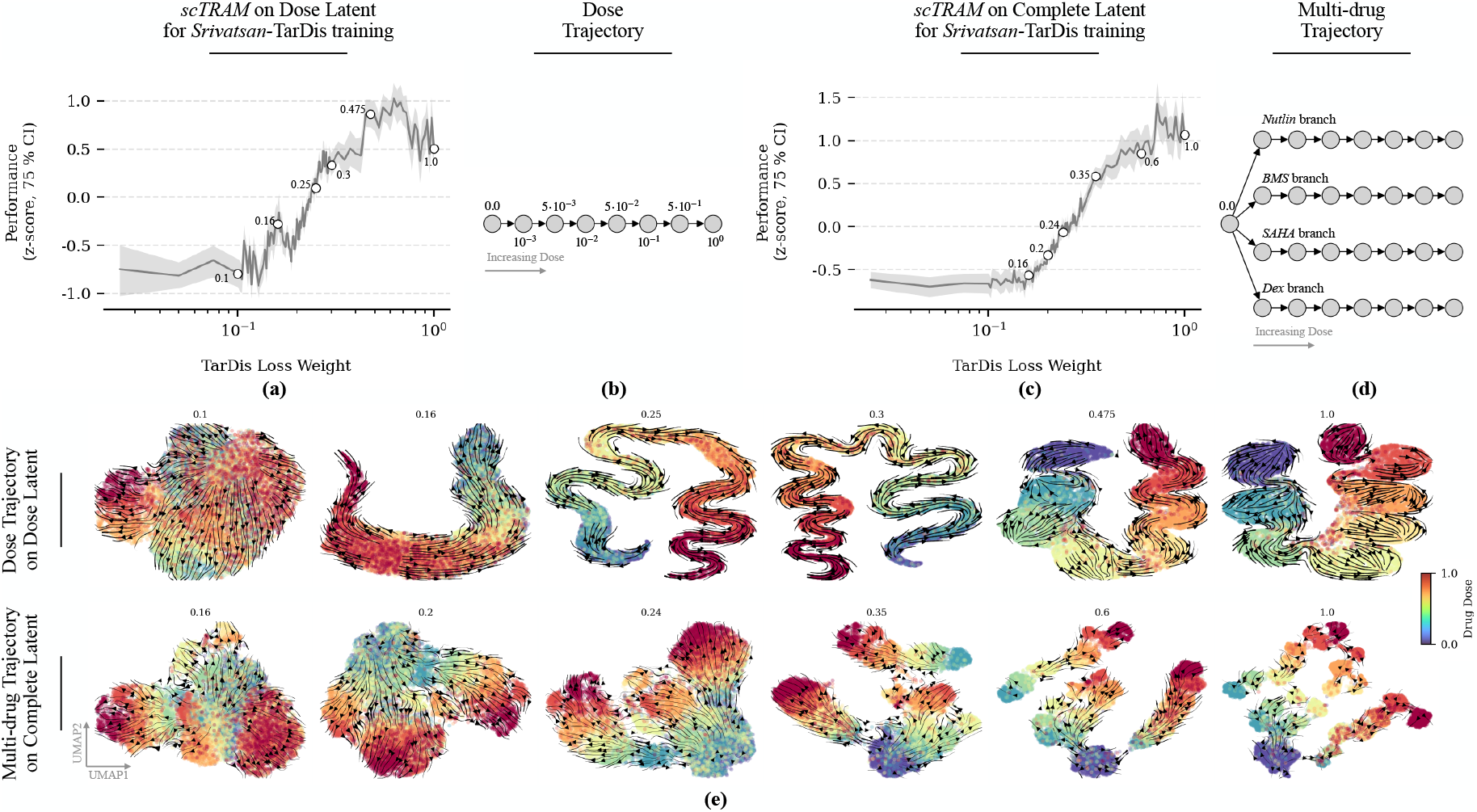
Improved trajectory representation fidelity with increasing disentanglement strength in the TarDis architecture, as quantified by the *scTRAM* suite. (a) In dose-specific latent dimensions, trajectory fidelity scores (*z*-normalized with 75% confidence intervals) exhibit improvement with increasing disentanglement weight. (b) Schematic of the dose-response trajectory showing the continuous progression across increasing drug concentrations, the trajectory used in experiment in Panel-a. (c) Analysis of the complete latent space confirms that *scTRAM* effectively quantifies trajectory preservation in complex, compositional embeddings containing multiple biological signals, with similar performance transitions. (d) Schematic of the multi-drug trajectory with four distinct branches progressing through increasing dose levels, the trajectory used in experiment in Panel-c. (e) Representative visualizations of trajectory embeddings with increasing TarDis disentanglement weights, showing the progressive formation of coherent trajectories as regularization strength increases. Cells are colored by dose level, and black arrows denote the local expected displacement vectors.

**Figure 7.**
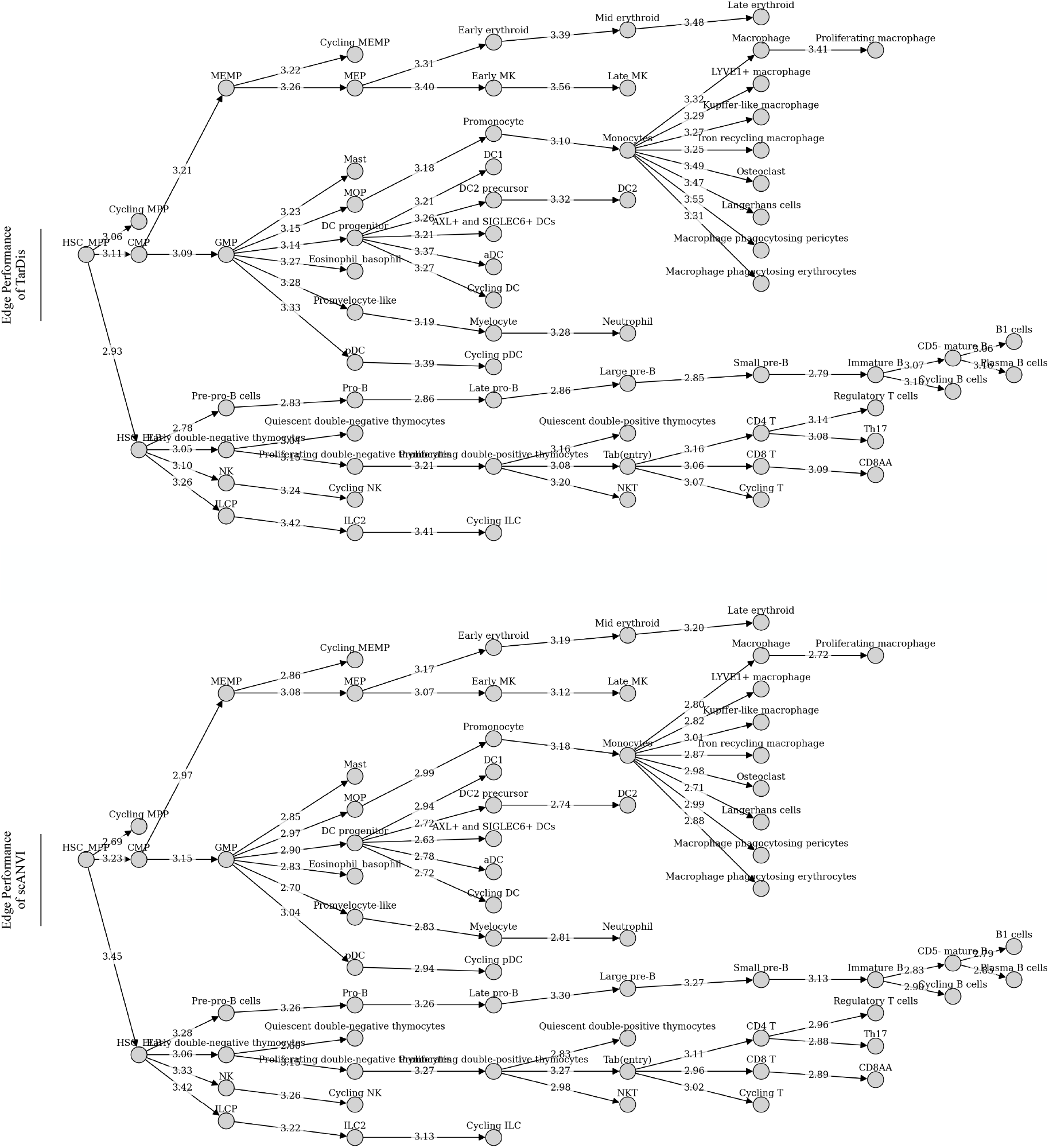
Detailed visualization of edge-level performance analysis for TarDis (top) and scANVI (bottom) representations on the complete hematopoietic manifold from the *Suo* dataset. Each node represents a distinct cell type within the hematopoietic lineage hierarchy, positioned according to developmental relationships and branching topology. Directed edges correspond to validated developmental transitions between cell types, with edge weights indicating the mean performance rank of that specific transition across all applicable *scTRAM* metrics and trajectory segments containing the edge (lower values indicate superior preservation). The graph encompasses the full spectrum of hematopoietic development, from multipotent HSC MPP to terminally differentiated lineages including myeloid (macrophages, neutrophils, eosinophils), lymphoid (T cells, B cells, NK cells), and MEM (megakaryocyte-erythroid-mast) branches. Edge performance values reveal model-specific biases in preserving distinct developmental transitions; TarDis demonstrates superior fidelity in lymphoid differentiation pathways (particularly B-cell maturation sequences with edge weights ≈ 2.8–3.0), whereas scANVI excels in early stem cell bifurcations and myeloid lineage transitions (edge weights ≈ 2.7–3.2). Notably, both models exhibit comparable performance in preserving MEM lineage transitions, suggesting that megakaryocyte-erythroid-mast differentiation pathways may be inherently more robust to representation distortions. The heterogeneous landscape of edge-specific performance underscores the necessity of trajectory-aware benchmarking for context-specific model selection, particularly when downstream analyses focus on discrete developmental subpaths or individual transition edges where aggregate metrics obscure critical localized differences in trajectory preservation fidelity.

**Figure 8.**
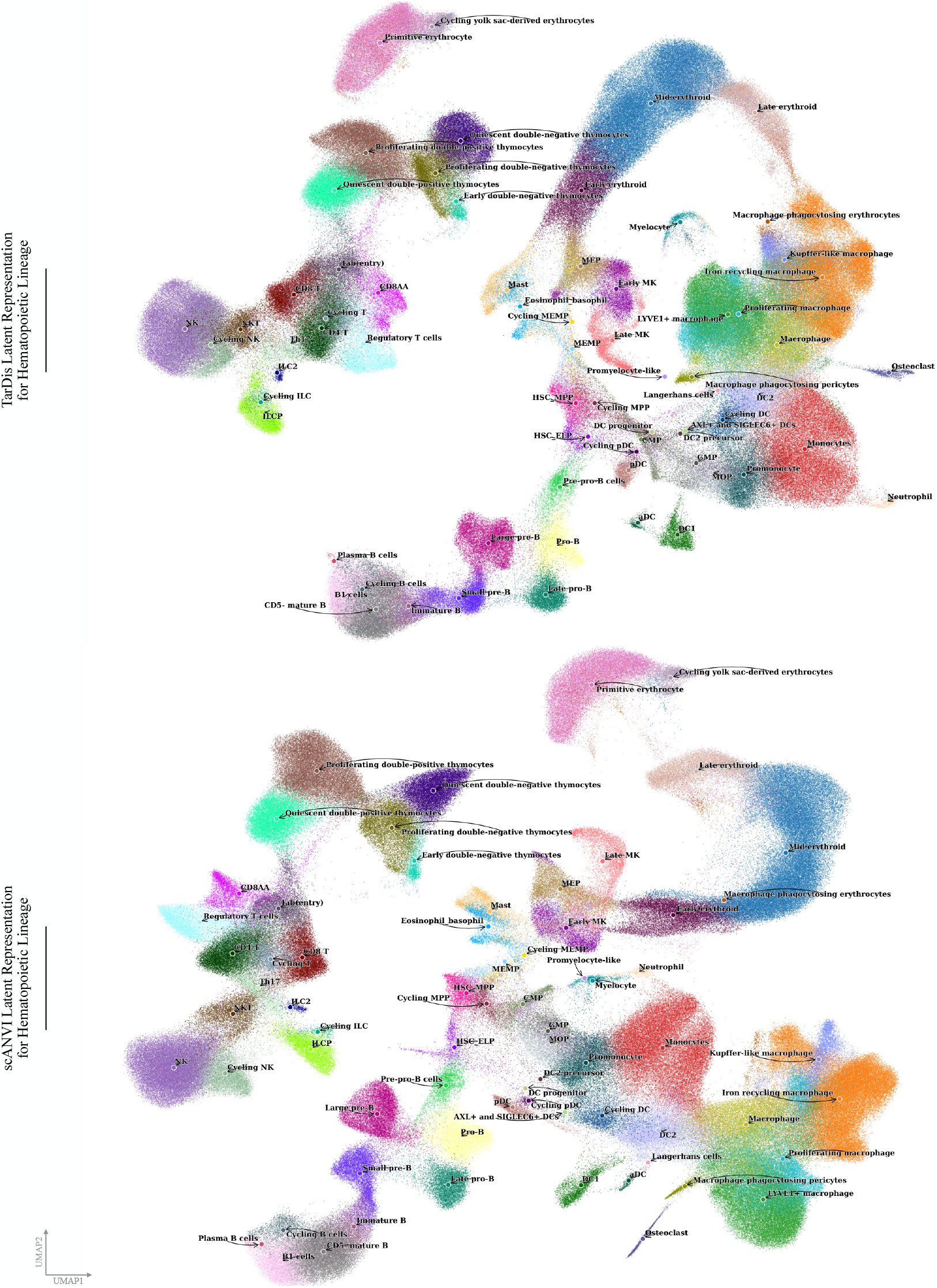
UMAP projections of the complete hematopoietic manifold from the *Suo* dataset, comparing trajectory preservation across TarDis (top) and scANVI (bottom) representations. Points represent individual cells colored by annotated cell type, revealing the complex landscape of developmental states spanning hematopoietic stem cells through terminally differentiated lineages. While these twodimensional projections facilitate visual inspection of cluster separation and gross topological relationships, UMAP’s stochastic neighbor embedding algorithm prioritizes local neighborhood preservation over global trajectory continuity, leading to artificial compression, stretching, and reordering of developmental paths that can fundamentally misrepresent the temporal sequence and branching topology of cellular differentiation processes. Researchers should exercise caution when interpreting developmental relationships from UMAP visualizations alone, particularly in complex systems where multiple lineages exhibit overlapping gene expression signatures or when trajectory branches span similar regions of the high-dimensional expression space, as the resulting two-dimensional layout may suggest spurious connections while obscuring genuine developmental pathways.

## 4. Conclusion

In this study, we introduced *scTRAM*, a principled benchmarking framework that systematically quantifies trajectory representation fidelity in single-cell embeddings across multiple complementary dimensions. Our framework addresses a critical gap in current evaluation protocols by providing quantitative measures of how effectively an embedding maintains the structure of ground-truth trajectories, thereby shifting from qualitative visual assessment toward principled, reproducible benchmarking. While *scTRAM* substantially advances quantitative trajectory evaluation, several important directions remain for future development. First, the current metric suite, though comprehensive, contains partially redundant measures whose relative informativeness may vary across biological contexts (refer to Figure S9); systematic meta-analysis across diverse single-cell atlases is needed to establish optimal metric selection criteria that maximize discriminative power while minimizing noise and redundancy. Second, the metrics should be categorized better to give more in-depth insights about the trajectory under evaluation, potentially integrating with language models to generate more accessible summaries for bench researchers. Third, as single-cell technologies increasingly capture multimodal measurements—spanning transcriptomics, epigenomics, proteomics, and spatial information—extending *scTRAM* to evaluate joint embeddings across these modalities represents a crucial next step for comprehensive trajectory benchmarking. Fourth, integrating *scTRAM* metrics directly into representation learning objectives offers a promising avenue for developing embeddings explicitly optimized for trajectory preservation, rather than relying on post-hoc evaluation of models trained with generic reconstruction or clustering objectives. As single-cell atlas construction increasingly emphasizes dynamic processes rather than static snapshots—from embryogenesis to immune repertoire dynamics to disease progression—we anticipate that trajectory-aware benchmarking and optimization will become essential components of representation learning pipelines, enabling more faithful recapitulation of the continuity underlying cellular state transitions and ultimately yielding deeper insights into the molecular mechanisms governing development, homeostasis, and pathology. ^4 5^

## A. Supplementary Figures

## B. Limitations and Assumptions

Despite *scTRAM*’s demonstrated utility in quantifying trajectory preservation across diverse biological contexts, several methodological and conceptual limitations warrant consideration. First, the framework’s reliance on *k*-nearest-neighbor graph construction introduces potential instability when embeddings produce disconnected components or highly segregated clusters. In particular, when an embedding partitions trajectory-integral cell types into disjoint submanifolds, the diffusion-based pseudotime inference procedure may fail and reflect spurious discontinuities rather than biological progression. In such scenarios, *scTRAM* potentially misevaluate representations that maintain trajectory-relevant gene expression covariance but sacrifice spatial contiguity. Consequently, *scTRAM*’s metrics should be interpreted cautiously in scenarios exhibiting pronounced clustering artifacts or when evaluating embeddings derived from methods optimized for discrete classification rather than continuous manifold learning.

Another significant limitation stems from the discreteness of certain metrics, particularly evident in adjacency-based evaluations such as binary graph accuracy. For trajectories with limited branching complexity or few constitutive cell types, these metrics exhibit quantization effects that reduce their discriminative power and amplify the influence of stochastic fluctuations. Specifically, on small trajectory graphs with *t* ≪ 10 nodes, the adjacency matrix *A*_inf_ ∈ {0, 1} ^*t*×*t*^ contains relatively few entries, causing even minor topological perturbations to induce substantial changes in metric values. This discretization effect introduces evaluation variance that may obscure subtle yet biologically significant differences between competing representations. While our multi-metric approach partially mitigates this concern through ensemble evaluation, the sensitivity–specificity trade-off remains suboptimal for developmental systems characterized by linear or minimally branched trajectories.

The binarization threshold applied to the weighted inter-type connectivity matrix *W*_inf_ represents another methodological constraint with substantive implications for *scTRAM*’s evaluation outcomes. Currently, this threshold is calibrated based on empirical performance on the *Suo* dataset, which introduces potential dataset-specific bias and limits generalizability across diverse experimental contexts. The absence of a principled, data-adaptive thresholding procedure means that connectivity assessments may be inconsistently stringent across different biological systems or embedding dimensionalities. Indeed, optimal threshold selection depends on complex factors including dataset size, trajectory complexity, and embedding properties, creating an implicit hyperparameter that requires careful tuning. Future iterations of *scTRAM* would benefit from a systematic approach to threshold determination that balances sensitivity to genuine trajectory connections against robustness to noise-induced spurious edges.

The framework also exhibits differential sensitivity to disruptions in distinct regions of the trajectory graph. Our ablation experiments demonstrated that *scTRAM* metrics respond more strongly to perturbations in densely connected regions or near branch points compared to terminal differentiation edges, potentially biasing overall evaluations toward preservation of stem and progenitor transitions at the expense of terminal lineage segments. This spatial heterogeneity in metric sensitivity complicates interpretation of aggregate scores and may inadvertently guide model selection toward representations that excel in preserving early developmental transitions while tolerating substantial distortion in terminal differentiation dynamics. Although our edge-specific decomposition approach and metrics specifically focused on terminal cell-states enables more granular performance assessment, it does not fully resolve the underlying imbalance in metric responsiveness across trajectory regions.

Our framework’s current implementation tends to weight each metric equally within its respective group, which may not optimally reflect their relative importance or informativeness across diverse biological contexts. This uniform weighting scheme might inadvertently amplify the influence of metrics that capture redundant aspects of trajectory fidelity while diluting the signal from those that measure unique structural properties (refer to Figure 9). The challenge of establishing context-appropriate weighting schemes is exacerbated by the absence of a systematic meta-analysis correlating metric performance with downstream analytical utility; consequently, practitioners must rely on domain knowledge or heuristic approaches to prioritize specific metrics for their particular biological questions while we are actively developing methods to address this limitation. A more principled approach would entail learning optimal metric weights from a diverse corpus of annotated single-cell datasets, but the current scarcity of ground-truth trajectory annotations limits the feasibility of such data-driven optimization. A more principled approach would entail learning optimal metric weights from a diverse corpus of annotated single-cell datasets—a direction we are actively pursuing through the ongoing curation of datasets with high-quality ground-truth trajectory annotations.

**Figure 9.**
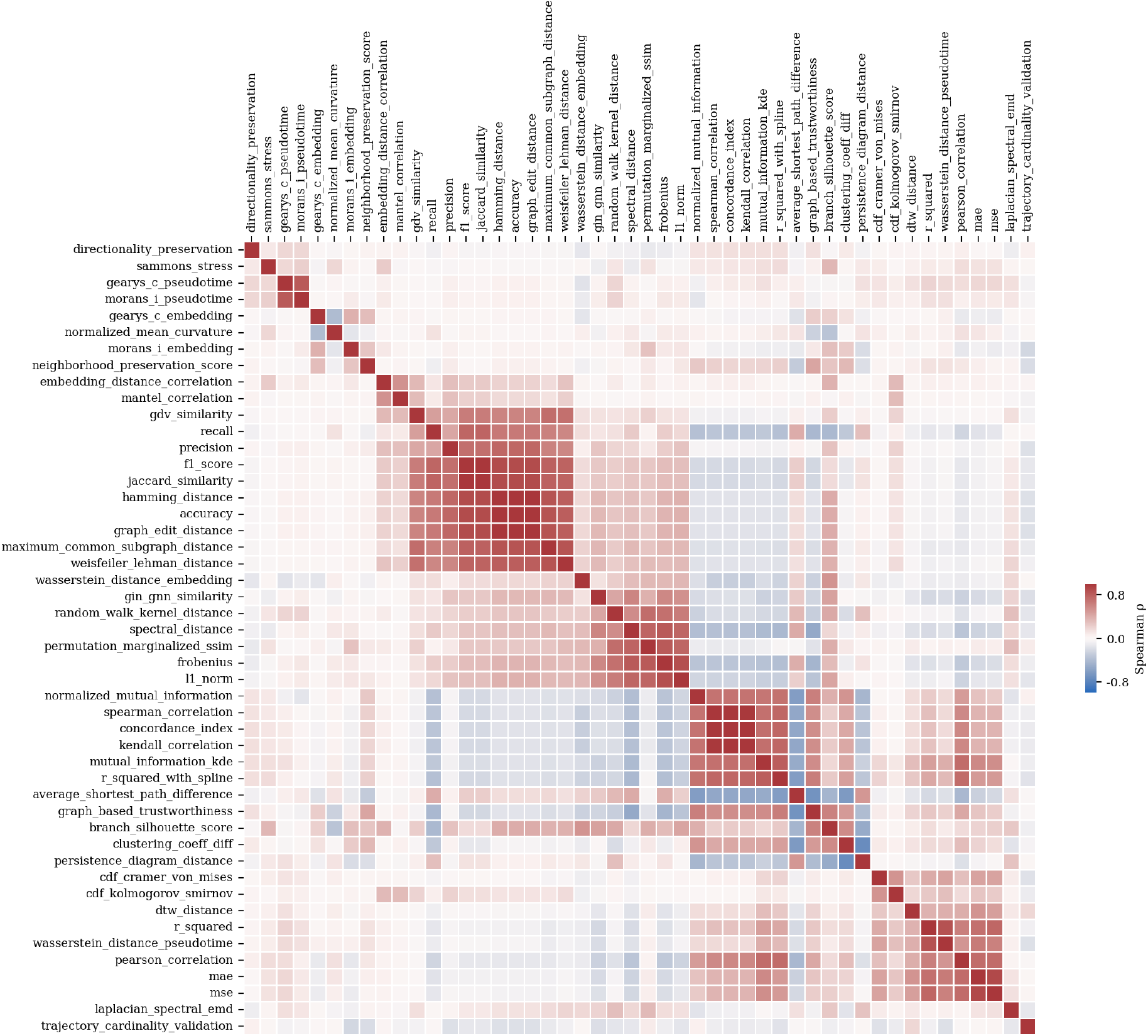
Pairwise correlation structure of *scTRAM* metrics reveals functional redundancy and orthogonality patterns across trajectory representation measures. This analysis was calculated exclusively on decomposed edge data from the Suo dataset and therefore represents a dataset-specific correlation structure that may not generalize across different biological contexts, trajectory topologies, or cellular systems. The heatmap displays pairwise Spearman correlation coefficients (*ρ*) between the 48 individual metrics comprising the *scTRAM* benchmarking suite, computed across all trajectory segments and model representations. Hierarchical clustering was applied using average linkage on correlation magnitude distances (1 −|*ρ*|) to group functionally similar measures and reveal the underlying correlation structure. The diverging color scale spans from *ρ* = − 0.8 (blue, indicating strong negative correlation) to *ρ* = +0.8 (red, indicating strong positive correlation), with white representing zero correlation (*ρ* ≈0). The correlation matrix reveals several distinct blocks of highly correlated metrics, particularly evident in the upper-left quadrant where embedding-based measures such as neighborhood preservation, manifold correlation, and various distance metrics exhibit strong positive correlations (*ρ >* 0.6), indicating substantial redundancy among trajectory representation measures that capture similar aspects of embedding fidelity. Conversely, several metric pairs demonstrate near-orthogonal behavior (*ρ* ≈ 0.1), such as between spectral distance measures and pseudotime correlation metrics, confirming that different metric groups capture complementary aspects of trajectory fidelity as theoretically expected from the *scTRAM* framework design. The observed correlation structure deviates substantially from the theoretical identity matrix that would characterize completely orthogonal metrics, highlighting both the inherent interconnectedness of trajectory fidelity measures and the need for systematic redundancy reduction to optimize the discriminative power and computational efficiency of the *scTRAM* framework. These correlation patterns suggest that a subset of highly informative, minimally redundant metrics could potentially achieve comparable benchmarking performance while reducing computational overhead. Future work should focus on developing principled metric selection pipelines that identify the most informative combinations within correlated metric groups, potentially through techniques such as principal component analysis of the correlation structure or information-theoretic approaches that maximize mutual information while minimizing redundancy across the metric suite.

Another limitation concerns *scTRAM*’s treatment of cellular heterogeneity within annotated cell types. The framework implicitly assumes that cells assigned to the same type exhibit comparable positional identity within the trajectory manifold, effectively collapsing intra-type variability when computing type-level adjacencies and pseudotimes. However, cellular states often exist along continua rather than in discrete clusters, with substantial heterogeneity in differentiation status even within expert-annotated types. By aggregating over this internal variance, *scTRAM* may obscure significant heterogeneity in trajectory representation quality within cell types, particularly for coarse-grained annotations that span substantial developmental windows. The pooling operations we employ when constructing *W*_inf_ effectively marginalize over this heterogeneity, potentially masking genuine embedding distortions that occur strictly within type boundaries. This limitation becomes especially pronounced in studies of transitional cell populations—such as those undergoing epithelial-mesenchymal transition or lineage reprogramming—where intra-type pseudotemporal variation constitutes a primary analytical target rather than a source of noise to be averaged away.

*scTRAM*’s current design primarily addresses trajectory fidelity in static snapshots rather than explicitly modeling longitudinal dynamics or temporal sampling effects. When applied to time-series experiments—where cells are sampled at discrete timepoints—the framework does not explicitly account for batch effects specific to temporal sampling, nor does it incorporate the additional constraints that true temporality imposes on trajectory structure. For instance, genuine time-course data inherently prohibits certain trajectory topologies, such as backward loops or temporally inconsistent branches, yet *scTRAM* lacks specialized metrics to penalize such violations. Similarly, the framework does not adequately capture velocity-field consistency or RNA velocity alignment, which provide orthogonal evidence for trajectory directionality. Integration of RNA velocity information— whether through explicit calculation of velocity-pseudotime concordance or through penalty terms for divergent flow fields—would substantially enhance *scTRAM*’s utility for dynamic biological processes.

Moreover, *scTRAM* presupposes the existence of a groundtruth reference trajectory, typically derived from expert annotation, lineage tracing, or temporal sampling. In many experimental contexts, however, such reference information remains incomplete, uncertain, or entirely absent, limiting the framework’s applicability to well-characterized systems. When reference trajectories are approximated from cluster-level relationships rather than derived from gold-standard lineage tracing, *scTRAM* evaluations inherit any errors or ambiguities in the reference construction. This circularity becomes particularly problematic when evaluating embeddings used to infer the very trajectories against which they are subsequently benchmarked. Additionally, the framework offers limited guidance for reconciling discrepancies between competing reference trajectories when multiple plausible models exist for a given biological system.

The framework’s emphasis on global trajectory preservation may inadvertently mask local distortions that, while small in aggregate, significantly impact biological interpretation of specific trajectory transitions. For instance, a representation might preserve overall pseudotemporal ordering and topological structure while severely distorting the geometry around critical bifurcation points or fate-decision boundaries. Although our edge-specific decomposition partially addresses this concern, the current metric suite lacks specialized sensitivity to these biologically crucial regions. Consequently, practitioners may select representations that excel on global measures while failing to maintain the fine-grained structure necessary for accurate inference of gene-regulatory networks, signaling dynamics, or transcription factor activities. Development of region-weighted or attention-based metrics that prioritize preservation of biologically significant submanifolds—such as critical transition states or regulatory checkpoints—would enhance *scTRAM*’s alignment with downstream analytical priorities.

From a computational perspective, *scTRAM*’s diffusion-based pseudotime inference procedure incurs substantial memory requirements for large-scale datasets, scaling as *O*(*n*^2^) with respect to cell count when employing dense matrix operations. Although sparse implementations significantly improve efficiency, the computational complexity still presents challenges for atlas-scale analyses with millions of cells, potentially necessitating subsampling strategies that may introduce sampling bias. Furthermore, the framework’s hyperparameter landscape—including *k* in kNN graph construction, *α* in damped diffusion, and the convergence tolerance *ε*—remains insufficiently characterized across diverse biological contexts, leaving practitioners with limited guidance for configuration decisions that may significantly impact evaluation outcomes.

Our focus on evaluating latent representations limits *scTRAM*’s applicability to directly assessing reconstruction quality in generative models. While latent fidelity constitutes a necessary condition for accurate modeling of trajectory structure, it provides only indirect evidence regarding the model’s capacity to generate realistic cells that respect developmental constraints. A comprehensive evaluation framework would ideally assess both the encoding and decoding components of generative models, examining whether generated cells maintain appropriate pseudotemporal positioning, preserve lineage-specific gene modules, and exhibit biologically plausible expression patterns. Extension of *scTRAM* to evaluate trajectory fidelity in generated or reconstructed expression space—rather than solely in latent representations—would provide a more complete assessment of model performance for trajectory-focused generative modeling tasks.

Finally, while *scTRAM* provides a comprehensive battery of complementary metrics spanning topological consistency, manifold continuity, and pseudotime alignment, the current suite lacks systematic quantification of uncertainty or statistical significance. Particularly for datasets with limited cell counts per trajectory segment, the framework does not adequately account for sampling variability or provide confidence intervals for metric values. This limitation becomes especially problematic when comparing embeddings whose performance differences fall within the expected range of stochastic variation. Furthermore, the absence of formal hypothesis testing procedures complicates determination of whether observed performance differentials reflect genuine representational advantages or merely arise from chance fluctuations in the evaluation process. Addressing this limitation would require development of principled bootstrap or permutation-based approaches for establishing statistical significance thresholds that account for trajectory complexity, cell count, and dataset heterogeneity.

Despite these limitations, *scTRAM* represents a substantial advancement over previous qualitative trajectory evaluation approaches by providing a systematic, quantitative framework that decompose trajectory fidelity into complementary axes of performance. By acknowledging these constraints and interpreting results with appropriate caution, practitioners can leverage *scTRAM* to make more informed decisions about representation selection while recognizing the inherent trade-offs between different aspects of trajectory preservation. Future refinements to the framework will focus on addressing these limitations through development of more robust graph construction methods, data-adaptive thresholding procedures, computational optimizations, and statistical significance testing, thereby enhancing *scTRAM*’s utility across diverse experimental contexts and biological systems.

## C. Experiment Insights

### C.1. Datasets

#### C.1.1. Suo dataset

Named in after the first author from the original publication, the *Suo* dataset offers a comprehensive single-cell transcriptomic landscape across nine prenatal human organs. It captures immune system development over embryonic time, highlighting organ-specific interactions and immune cell differentiation (Suo et al., 2022).

##### Number of Samples

Out of an initial total of 908,178 cells, 841,922 passed quality control filtering based on established single-cell processing standards (Heumos et al., 2023).

##### Number of Features

The dataset originally profiled 33,538 genes. After preprocessing, 8,192 highly variable genes (HVGs) were retained, following best practices in the field (Heumos et al., 2023).

##### Source

The raw dataset is publicly accessible via ArrayExpress under accession ID E-MTAB-11343. Processed data in AnnData format can be retrieved from Developmental Cell Atlas portal. Additional annotated metadata is hosted on the CellxGene platform, as described in (Biology et al., 2023).

#### C.1.2. Garcia-Alonso dataset

The *Garcia-Alonso* dataset offers a comprehensive single-cell and spatial transcriptomic map of human gonadal development during the first and second trimesters. By integrating single-cell RNA sequencing, spatial transcriptomics, chromatin accessibility assays, and fluorescent microscopy, this dataset delineates the cellular and molecular events underpinning sex determination and differentiation in human gonads. It identifies key somatic cell states, including a bipotent early supporting population that, in males, upregulates the testis-determining factor SRY, and in females, gives rise to granulosa cells. Additionally, the dataset characterizes unique macrophage populations in the developing testis, providing insights into the signaling interactions between germline and somatic cells during gonadogenesis (Garcia-Alonso et al., 2022).

##### Number of Samples

Out of an initial total of 222,779 cells, 219,731 passed quality control filtering based on established single-cell processing standards (Heumos et al., 2023). For this study, we subsampled the dataset to 26,185 cells to enable faster *scTRAM* experimentation and reduce training and benchmarking time for incremental training experiment discussed in Figure 4. The subsampling was performed using a stratified approach based on the cell type annotation. Specifically, the method retains all cells from underrepresented cell types while capping the number of cells from any overrepresented group to a maximum of 1000. This is done to preserve diversity while preventing dominant cell types from skewing the dataset. A fixed random seed was used to ensure reproducibility during the selection of cells from overrepresented categories.

##### Number of Features

The original gene expression matrix included 29,052 genes. Following standard preprocessing protocols, we identified 8,192 highly variable genes (HVGs) using the full dataset prior to subsampling (219,731 cells), in accordance with best practices for single-cell transcriptomic analysis. (Heumos et al., 2023).

##### Sourc

Raw single-cell RNA-seq data are publicly available from ArrayExpress under accession number E-MTAB-10551. Additional annotations and visualizations are accessible via the CellxGene platform, as described in (Biology et al., 2023).

#### C.1.3. La-Manno dataset

The *La Manno* dataset, named after the first author of the original study, offers a comprehensive single-cell transcriptomic analysis of neurogenesis within the developing mouse hippocampus. Focusing on the dentate gyrus region, the study captures cellular dynamics at two critical postnatal stages—day 0 (P0) and day 5 (P5). This dataset delineates the complex developmental trajectories of neural progenitor cells as they differentiate into various cell types, including astrocytes, oligodendrocyte precursor cells (OPCs), granule neurons, and pyramidal neurons. The high-resolution data facilitate the exploration of lineage branching and maturation processes during early postnatal brain development (La Manno et al., 2018).

##### Number of Samples

A total of 18,213 single cells were included in the version of the dataset used for this study.

##### Number of Features

The dataset originally profiled 27,998 genes. After preprocessing, 5,000 highly variable genes (HVGs) were retained, following best practices in the field (Heumos et al., 2023).

*Source*: The dataset is accessible through the scVelo Python package and can be loaded using the function`dentategyrus lamanno` under `scvelo.datasets`. Additional information and documentation are available at scVelo Dentate Gyrus La Manno Dataset Documentation (Bergen et al., 2020).

#### C.1.4. rivatsan dataset

The *Srivatsan* dataset, named after the first author of the original study, generated using the sci-Plex platform which leverages nuclear hashing for multiplexing, captures single-cell transcriptional responses to a wide array of chemical treatments. This dataset spans three human cancer cell lines subjected to 188 different small molecules, allowing for detailed investigation of dosage-dependent effects and differential drug sensitivities. In a single large-scale screening experiment, nearly 650,000 single-cell transcriptomes were collected from around 5000 unique conditions. The dataset reveals diverse cellular responses to drug treatments, patterns shared among chemically related compounds, and subtle distinctions within specific drug categories, notably among histone deacetylase (HDAC) inhibitors (Srivatsan et al., 2020).

##### Number of Samples

A total of 14,811 single cells were included in the version of the dataset used for this study.

##### Number of Features

Gene expression was measured across 4,999 genes.

##### Source

The raw and processed data can be accessed through the NCBI Gene Expression Omnibus (GEO) under accession number GSE139944. The version used here is a preprocessed and subsetted form, consistent with the approach taken in the CPA framework (Lotfollahi et al., 2023), and was made available by the CPA authors. Our team did not apply additional filtering or modification to the dataset.

### C.2. Models

All experiments that use the *Suo* dataset were run with exactly the same configurations as the original TarDis paper; the full settings are available in the TarDis public repository. In addition, we relied on a pre-trained model object kindly provided by the TarDis authors, which streamlined our downstream analyses and the benchmarking of the *scTRAM* suite. For all other experiments based on the *Garcia-Alonso, La-Manno*, and *Srivatsan* datasets, we principally employed each method’s default parameters; the full configuration files are archived in our public repository to ensure complete reproducibility.

#### C.2.1. scVI

scVI employs a deep variational autoencoder architecture that models single-cell gene expression through a zeroinflated negative binomial (ZINB) likelihood with learned dispersion parameters and dropout probabilities (Lopez et al., 2018). The generative model assumes that observed UMI counts *x*_*n,g*_ for cell *n* and gene *g* arise from a ZINB distribution parameterized by rate *λ*_*n,g*_, dispersion *θ*_*g*_, and dropout probability *π*_*n,g*_; these parameters are computed via neural networks that decode a low-dimensional latent representation *z*_*n*_ ∼ *N* (0, *I*) drawn from a standard Gaussian prior. The inference network approximates the posterior *q*(*z*_*n*_ |*x*_*n*_) through a recognition model that encodes observed expression profiles into distributional parameters (*µ*_*n*_, *σ*_*n*_) of a diagonal Gaussian, enabling efficient amortized variational inference via the reparameterization trick. During training, the evidence lower bound (ELBO) balances reconstruction accuracy—measured by the ZINB loglikelihood—against KL divergence between the approximate posterior and prior, regularizing the latent space toward a unit Gaussian while maintaining sufficient expressivity to capture biological variation. The model’s capacity to handle zero-inflation and technical noise through explicit dropout modeling makes it particularly suitable for sparse single-cell count data, while the continuous latent space naturally accommodates smooth biological transitions and manifold structures underlying cellular differentiation processes.

#### C.2..2. scANVI

scANVI extends the scVI framework by incorporating semisupervised learning to leverage partially available cell-type annotations during representation learning (Xu et al., 2021). The generative model augments the scVI likelihood with a categorical distribution over cell types, where the type probability *p*(*c*_*n*_ = *k*| *z*_*n*_) is computed through a classification head that maps latent representations to type probabilities via a softmax-activated neural network. For annotated cells, the model enforces consistency between predicted and observed cell types through a cross-entropy loss term, while unlabeled cells contribute only to the standard scVI reconstruction objective. The joint training procedure optimizes a composite ELBO that includes the original scVI terms plus the supervised classification loss, weighted by the annotation frequency to balance supervised and unsupervised components. This semi-supervised architecture enables the model to learn latent representations that simultaneously preserve transcriptional similarity and respect known cell-type boundaries, often yielding embeddings with enhanced biological interpretability compared to purely unsupervised approaches. The incorporation of type information during training tends to produce more structured latent spaces where cells of the same type cluster together while maintaining smooth transitions between related types, based on our preliminary results, making scANVI particularly effective for scenarios where trajectory fidelity and cell-type preservation are both priorities.

#### C.2.3. inVAE

inVAE is a deep generative model that decomposes latent space into two principal components: an invariant subspace, **z**_inv_, capturing condition-independent biological signals, and a spurious subspace, **z**_spur_, which encodes nuisance variation that differs across user-defined conditions (Aliee et al., 2024). A single decoder reconstructs observations (e.g., gene counts) from the concatenation (**z**_inv_, **z**_spur_, *ℓ*), where *ℓ* commonly denotes a library-size or scaling factor. The training objective augments the standard VAE ELBO with three key regularizers: *(i)* A Hilbert–Schmidt Independence Criterion (HSIC) term that penalizes residual mutual information between **z**_inv_ and the conditioning covariates, ensuring **z**_inv_ remains invariant, *(ii)* a total-correlation penalty designed to factorize the joint posterior, thereby discouraging undesired dependencies among latent dimensions and improving latent disentanglement, *(iii)* a cosine alignment term that brings together **z**_inv_ embeddings from matched conditions, reinforcing the notion of a shared manifold across experimental batches or perturbations. Through this structured regularization, inVAE establishes a ‘bijection’ from the invariant latent space to cross-condition manifolds, rendering **z**_inv_ stable against perturbations in **z**_spur_. Empirically, the approach demonstrates strong transferability across diverse experimental settings by preserving only biologically conserved information in **z**_inv_ while isolating condition-specific or nuisance variations in **z**_spur_.

#### C.2.4.TarDis

TarDis is a novel deep generative model built upon a variational autoencoder (VAE) framework, specifically engineered for the targeted disentanglement of multiple covariates in complex single-cell genomics datasets (Inecik et al., 2024). The model constructs a sophisticated latent representation *z*_*n*_ = (*z*_*n*0_, [*z*_*nk*_]*k* ∈*J*_*k*_) where each targeted covariate *k* receives its own dedicated latent subspace *z*_*nk*_, while *z*_*n*0_ captures residual biological variation independent of the targeted covariates. The model employs a composite loss function that combines standard VAE objectives (reconstruction loss using negative binomial distribution for count data, and KL divergence regularization) with targeted auxiliary losses ℒ_*C*_ that enforce disentanglement through self-supervised contrastive learning. The core innovation lies in its covariate-specific loss components that employ a self-supervised contrastive learning strategy: for each covariate, the model generates positive and negative data point pairs based on covariate similarity, then applies four distinct loss terms that simultaneously pull together representations sharing the same covariate value in the reserved latent space *znk* while pushing apart different covariate values, and conversely ensures the unreserved space *zn*0 remains distant from all covariate conditions. This architectural design enables TarDis to handle both categorical covariates (such as disease status, tissue type) and continuous covariates (such as age, drug dosage) within a unified framework, with continuous variables receiving distance-weighted negative pair losses that preserve their natural ordering and create interpretable gradient representations in the latent space. TarDis addresses fundamental challenges in singlecell genomics where gene expression patterns emerge from overlapping biological processes and technical artifacts, requiring precise separation of invariant biological signals from spurious correlations. Its unique capability to generate ordered latent representations for continuous covariates enables previously unfeasible hypothesis-driven analyses, such as isolating organ-specific developmental gene expression patterns while maintaining batch correction, or examining dose-dependent drug responses independent of patient characteristics.

#### C.2.5. Harmony

Harmony employs an iterative correction algorithm that removes batch effects from existing embeddings through geometric transformations rather than learning representations de novo (Korsunsky et al., 2019b). The method begins with a pre-computed embedding (typically from principal component analysis) and applies successive corrections that align corresponding cell populations across different batches or datasets. The algorithm models batch effects as smooth deformations of the embedding space and estimates these transformations through kernel-based smoothing operations that preserve local neighborhood structure while correcting global alignment. Harmony iteratively updates cell coordinates by computing weighted averages within local neighborhoods, where weights are determined by similarity measures that account for both geometric proximity and batch identity. The correction procedure continues until convergence, yielding embeddings where batch-specific effects are minimized while biological variation is preserved. The method’s reliance on geometric correction rather than generative modeling makes it computationally efficient and broadly applicable to diverse types of pre-computed embeddings, though its performance depends critically on the quality of the initial representation and the assumption that batch effects manifest as smooth, correctable distortions.

#### C.2.6. PCA

PCA (Principal Component Analysis) computes lowdimensional embeddings through eigendecomposition of the covariance matrix, identifying orthogonal directions of maximum variance in the high-dimensional expression space (Abdi & Williams, 2010). The method begins by centering the data matrix and computing its sample covariance, then extracts the top eigenvectors corresponding to the largest eigenvalues to form the embedding basis. Each principal component represents a linear combination of genes that captures a distinct mode of variation across cells, with components ordered by their explained variance contribution. The embedding coordinates are obtained by projecting the original high-dimensional data onto the principal component subspace, yielding a low-dimensional representation that optimally preserves the global variance structure of the dataset. While PCA provides a computationally efficient and theoretically grounded dimensionality reduction approach, its linear nature limits its capacity to capture complex nonlinear relationships that characterize many biological processes. The method’s assumption of Gaussian-distributed data and its sensitivity to outliers can further impact performance on single-cell datasets, which typically exhibit high sparsity, technical noise, and complex distributional properties that violate standard PCA assumptions.

### C.3. Code Availability

All code supporting this work is publicly available at GitHub: https://github.com/theislab/sctram.

### C.4. Compute Resources and System Configuration

We developed and evaluated *scTRAM* using two distinct computational configurations. The primary system consisted of high-performance hardware specifically configured for efficient parallel processing of large-scale single-cell datasets, with detailed specifications provided below. Notably, while our MacBook Pro systems with M1 processors proved sufficient for development, testing, and smaller-scale analyses—demonstrating that *scTRAM*’s core functionality can be executed on standard consumer hardware—we utilized the high-performance system primarily to parallelize experiments and accelerate large-scale benchmarking. The *scTRAM* package was developed within mamba environments (Gu & Dao, 2023) as specified in our GitHub repository, with environment specification files that ensure computational reproducibility across platforms and prevent package inconsistencies that could affect analytical outcomes. The mamba environment is provided for clearer reproducibility, allowing users to create identical software configurations regardless of their underlying operating system. The package is currently under active development and will be released through both pip and conda-forge distribution channels to facilitate broader accessibility and integration into existing single-cell analysis workflows.

Our primary computational infrastructure employed for *scTRAM*’s development and evaluation consisted of highperformance hardware specifically configured for efficient parallel processing of large-scale single-cell data matrices. Our computational nodes were equipped with dual Intel Xeon Gold 6230 processors operating at a base frequency of 2.1 GHz with boost capabilities up to 3.9 GHz, providing 20 cores per processor for robust parallel execution of computational tasks. System memory consisted of 256 GB DDR4 RAM per computational node, enabling efficient handling of large sparse matrices without excessive memory swapping or I/O bottlenecks that would otherwise compromise computational efficiency. For processing extensive datasets, we allocated 64 GB of memory, while smaller datasets with reduced dimensionality or sparsity were processed with a 16 GB memory allocation, which optimized resource utilization while maintaining computational throughput. All computational tasks were orchestrated through an internal SLURM-based compute cluster configured for dynamic resource allocation based on job priority and current system load (Yoo et al., 2003). This orchestration allowed parallel execution of *scTRAM*’s component algorithms—such as graph construction, diffusion map computation, and metric evaluation—across multiple CPU cores. The specialized infrastructure facilitated efficient parallel execution of benchmarking experiments across multiple dataset configurations, metric combinations, and representation models without introducing computational constraints that might have biased our comparative analyses. The computational resources described herein proved sufficient for all experimental evaluations reported in this work; no additional computational infrastructure was necessary for reproducing the presented results or extending the analysis to datasets of comparable scale and complexity.

## D. Supplementary Methods

### D.1. Trajectory Decomposition Algorithms

#### D.1.1. Alternative 1

The trajectory decomposition algorithm partitioned the reference lineage graph *G*_ref_ into interpretable subgraphs capturing critical developmental transitions through a multistage procedure. First, all branch points—nodes with outdegree ≥ 2—were identified; for each such node *v*, the maximal subtree rooted at *v* was extracted by computing its descendant set *D* (*v*) and forming the induced subgraph *G*_ref_[*D* ∪ (*v*) {*v*}]. To qualify for inclusion, subtrees were required to contain at least *τ* nodes (default *τ* = 3) and exhibit a single root node, ensuring structural coherence as a connected trajectory segment. Concurrently, linear paths emanating from each branch point successor were traced iteratively: starting at each immediate child *u* of *v*, nodes were appended sequentially until encountering a node with out-degree ≠ 1 or a leaf, forming a path *P* = (*v, u*, …, *w*). Paths shorter than *τ* nodes were discarded; others were retained as candidate segments. To capture the dominant developmental axis, the longest path in *G*_ref_ was computed via dynamic programming on the directed acyclic graph, with ties broken arbitrarily. This path, representing the maximal linear progression through the lineage hierarchy, was included provided its length met the *τ* threshold. Finally, to handle root nodes not associated with branch points, the algorithm extracted subtrees originating from each root *r* with in-degree 0, again requiring size ≥ *τ* and single-rootedness. All surviving subgraphs were assigned unique identifiers through appending incremental indices to their trajectory metadata. The union of these procedures yielded a comprehensive set of trajectory segments spanning bifurcating subtrees, linear differentiation paths, and the principal lineage backbone, enabling multi-scale fidelity assessment across *G*_ref_’s topological hierarchy.

#### D.1.2. Alternative 2 (adopted in this study)

The trajectory decomposition algorithm was designed to partition a directed acyclic graph (DAG) *G*= (*V*, ℰ) with exactly one root node into a collection of connected subgraphs that collectively preserved hierarchical, linear, and branching trajectory structures while satisfying a minimum size constraint *k*_min_. The algorithm proceeded through four complementary decomposition strategies applied sequentially to *G*, each generating candidate subgraphs that were validated against structural criteria before inclusion. First, hierarchical functional modules were identified by iterating over all branch points (nodes with out-degree ≥ 2) and the root; for each such node *v*, the induced subgraph over *v* and its descendants was extracted, ensuring coverage of all downstream trajectories originating from key bifurcation events. Second, the longest path *P*_max_ in *G*was computed via dynamic programming, then decomposed into all contiguous subsequences of length ≥ *k*_min_, including overlapping windows to capture local linear progression patterns. Third, branch-to-leaf trajectories were enumerated by identifying all leaf nodes (out-degree 0) and extracting the shortest path from the root to each leaf, followed by analogous windowed decomposition to handle polytomous branching. Fourth, stage-wise decomposition was performed by tracing linear chains from each branch point’s immediate successors until subsequent bifurcations, prepending the branch point to each chain to anchor substructures at biologically meaningful decision points. Candidate subgraphs were retained only if they satisfied three validity conditions: *(i)* the subgraph contained at least *k*_min_ nodes; *(ii)* it was weakly connected; *(iii)*it possessed exactly one root node (in-degree 0 within the subgraph). Overlapping regions between adjacent subgraphs were intentionally preserved to ensure redundant coverage of transition zones between trajectory segments. The union of valid subgraphs from all four strategies formed the final decomposition, with each subgraph annotated to record its constituent nodes and edges. This multi-resolution approach guaranteed that both global hierarchy (via descendant modules and longest paths) and local transitions (via branch-anchored chains and leaf-oriented paths) were represented, enabling granular performance analysis across all scales of *G*’s topology.

### D.2. Descriptions for scIB Metrics

- **Bio-conservation**
  - Cell Type Average Silhouette Width
  - Local Inverse Simpson’s Index (cLISI)
  - Normalized Mutual Information
  - Adjusted Rand Index
  - Isolated Label F1 Score
- **Batch Correction**
  - Graph Connectivity
  - Batch Average Silhouette Width
  - k-nearest-neighbor Batch Effect Test
  - Principal Component Regression
  - Local Inverse Simpson’s Index (iLISI)
- **Other**
  - **Aggregate Scores**
  - Average Silhouette Width
  - Mutual Information
  - Rand Index

#### D.2.1. Average Silhouette Width

The average silhouette width (ASW) (Rousseeuw, 1987b) quantified how cohesively cells clustered by transcriptional profile while remaining separable from neighboring clusters, addressing whether biological identity was preserved after integration. For each cell **x**_*n*_, the silhouette coefficient s(**x**_*n*_) was computed as 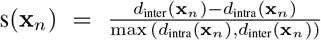 where *d*_intra_(**x**_*n*_) represented the mean Euclidean distance to other cells in the same cluster, and *d*_inter_(**x**_*n*_) denoted the minimum mean distance to cells in any other cluster. The global ASW was obtained by averaging s(**x**_*n*_) over all *N*_*C*_ cells: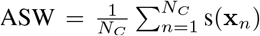. Scores spanned [*−*1, 1], with 1 indicating perfect cluster separation, 0 ambiguous boundaries, and -1 severe misclassification; computational complexity scaled as 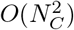 due to pairwise distance calculations. The metric provided an intuitive geometric assessment of cluster fidelity and batch mixing (Rousseeuw, 1987a), but its sensitivity to outlier cells and quadratic runtime limited utility in ultra-large datasets.

#### D.2.2. Cell Type Average Silhouette Width

The cell type average silhouette width (celltypeASW) (Luecken et al., 2022) quantifies how well a low-dimensional embedding preserves discrete cell type annotations by measuring clustering fidelity. For each cell *i*, the silhouette coefficient *s*(*i*) ∈ [−1, 1] is computed as (*b*(*i*) −*a*(*i*))*/* max {*a*(*i*), *b*(*i*)}, where *a*(*i*) denotes the mean intra-cluster distance to other cells in *i*’s type, and *b*(*i*) the mean distance to cells in the nearest neighboring cluster. The metric aggregates these values across all cells and applies an affine transformation to rescale the mean silhouette width into [0, 1]: celltypeASW = (ASW_*c*_ +1)*/*2, where ASW_*c*_ is the mean silhouette coefficient over cell type labels. Values near 1 indicate compact, well-separated cell type clusters; values near 0 suggest overlapping or poorly resolved types. The computation scales quadratically with cell count *n* due to pairwise distance calculations. A principal strength lies in its interpretability as a normalized measure of cluster cohesion and separation; however, the metric assumes cell types form convex, isolated groups and may underestimate biological continuity or hierarchical differentiation.

#### D.2.3. Batch Average Silhouette Width

The batch average silhouette width (Batch ASW) quantifies the degree to which batch effects persist in an integrated dataset, answering whether technical variations across batches have been sufficiently mitigated to prevent obscuring biological signal. The metric leverages the silhouette coefficient, which measures the compactness of each batch’s embedding relative to other batches; specifically, for each cell **x**_*n*_ with batch label *j*, the silhouette coefficient *s*_batch_(**x**_*n*_) compares the average distance to cells in the same batch against those in the nearest other batch. To compute Batch ASW, a normalized score is derived per batch label *j* as 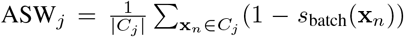, where *C*_*j*_ denotes cells in batch *j*, followed by averaging across all batches: 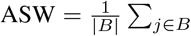 batchASW_*j*_ (Luecken et al., 2022). The score ranges between 0 and 1, with lower values indicating better batch mixing; a score near 0 implies minimal batch effect, while higher values reflect persistent technical variation (Haghverdi et al., 2018). The computational complexity is dominated by the pairwise distance calculations for silhouette coefficients, requiring *O*(*n*^2^) operations for *n* cells. A principal strength lies in its interpretability, combining intra-batch cohesion and inter-batch separation into a single metric; however, its reliance on predefined batch labels and sensitivity to cluster granularity can conflate biological heterogeneity with technical artifacts. Additionally, the quadratic scaling limits applicability to very large datasets.

#### D.2.4. DIsolated Label F1 Score

The Isolated Label F1 Score evaluates how effectively an integration method preserves the identity of rare cell types across batches, addressing whether biologically distinct but infrequent cell populations remain separable after integration. Precision quantifies the fraction of correctly identified cells within a predicted cluster relative to all cells assigned to it, defined as Precision = TP*/*(TP + FP), while recall measures the proportion of true positives captured among all actual members of a cell type, given by Recall = TP*/*(TP +FN). The F1 score harmonizes these as 2 × (Precision × Recall)*/*(Precision + Recall), providing a balanced measure of clustering accuracy. For isolated labels—cell identities present in few batches—the score computes class-wise F1 by optimizing cluster assignments to maximize per-label accuracy, thereby assessing preservation of rare cell types in imbalanced single-cell datasets (Sokolova & Lapalme, 2009; Luecken et al., 2022). While the canonical scIB implementation derives F1 via cluster labels, our analysis instead computed the isolated label average silhouette width (ASW) from the `scib-metrics` package (Gayoso, 2025), which evaluates separation quality analogously but with lower computational complexity. Scores range from 0 to 1, where higher values indicate better retention of rare cell identities; the complexity for ASW-based computation scales linearly with the number of cells and labels, *O*(*nt*). A principal strength lies in handling class imbalance inherent to single-cell genomics, directly targeting integration efficacy for rare populations. However, performance depends critically on the accuracy of prior annotations, and the substitution of ASW for cluster-based F1, though faster, conflates separation quality with cluster purity.

#### D.2.5. Mutual Information

Mutual information (MI) quantified the reduction in uncertainty about the inferred trajectory given knowledge of the reference trajectory, thereby measuring their shared information (Duncan, 1970; Kraskov et al., 2004). Formally, for random variables 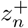 (inferred trajectory components) and 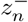 (reference trajectory components), MI was computed as:

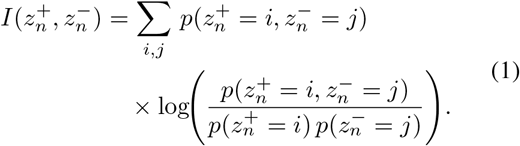

where 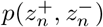 denotes their joint probability distribution, and 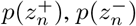 their marginals. MI is non-negative and symmetric; a value of 0 indicates statistical independence between trajectories, while higher values reflect increasing dependency. Estimation employed k-nearest-neighbor entropy approximation (Kraskov et al., 2004), with computational complexity dominated by pairwise distance calculations (*O*(*n*^2^) for *n* cells). A key advantage lies in MI’s sensitivity to non-linear associations beyond linear correlation metrics; however, accurate estimation requires careful hyperparameter selection for neighborhood size and becomes unreliable in high-dimensional spaces due to the curse of dimensionality.

#### D.2.6. Normalized Mutual Information

Normalized mutual information (NMI) quantified the preservation of cell-type label information after data integration, addressing whether inferred cell clusters correspond to known biological types. The metric was computed as the mutual information (MI) between the cluster assignments before 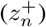 and after 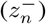 integration, normalized by the geometric mean of their entropies to mitigate biases from dataset size and cluster entropy variability:

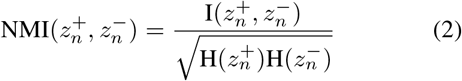

where 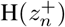 and 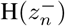 denote the Shannon entropies of each clustering. NMI ranges from 0 (independent clusterings) to 1 (perfect correspondence), with higher values indicating greater biological consistency; this bounded scale enabled direct comparisons across integration methods and datasets. The metric provided an intuitive assessment of label preservation but exhibited diminishing sensitivity as clusterings approached perfection, struggling to distinguish nearly optimal from ideal solutions (Vinh et al., 2010). While NMI’s normalization addressed critical scaling concerns inherent to raw MI, its reliance on predefined celltype labels limited applicability to fully annotated datasets (Luecken et al., 2022).

#### D.2.7. Rand Index

The Rand index (RI) quantified the agreement between the inferred cell-type clustering and a ground-truth reference partitioning, answering whether pairs of cells preserved their co-clustering relationships across both partitions. Its computation formalized clustering similarity through pairwise comparisons: true positives (TP) counted cell pairs co-clustered in both partitions; true negatives (TN) counted pairs separated in both; false positives (FP) and false negatives (FN) accounted for discrepancies. RI was computed as the ratio of concordant pairs to all possible pairs, expressed as 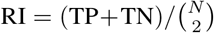, where *N* denoted the total number of cells. RI spanned [0, 1], with higher values indicating stronger clustering concordance; however, its n ïve calculation incurred *O*(*N* ^2^) complexity due to exhaustive pairwise comparisons. While RI provided an intuitive assessment of global clustering fidelity, its interpretation required caution in unbalanced cluster distributions, as it exhibited bias toward larger clusters and lacked adjustment for chance agreement. Its principal strength lay in invariance to label permutations and cluster numbering; conversely, its sensitivity to granularity mismatches and quadratic scaling limited utility in large-scale single-cell atlases.

#### D.2.8. Adjusted Rand Index

The Adjusted Rand Index (ARI) quantified the statistical significance of cluster label agreement between two partitions, adjusting for chance overlap; it addressed the biological question of whether cell type clusters inferred from integrated data matched ground-truth annotations better than random expectation. The ARI was computed by first enumerating all pairs of cells and categorizing them into four groups: pairs clustered together in both partitions (*a*), only the first (*b*), only the second (*c*), or neither (*d*). Letting *n*_*ij*_ denote the count of cells shared between cluster *i* in the inferred partition and cluster *j* in the reference, the index was expressed as:

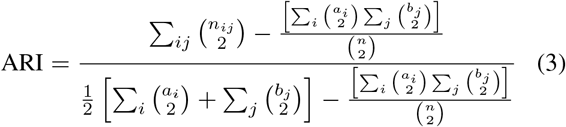

where *a*_*i*_ and *b*_*j*_ represented cluster sizes in each partition, and 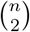 computed the total cell pairs. This formulation normalized the Rand Index (RI) by its expected value under hypergeometric distribution assumptions (Hubert & Arabie, 1985; Luecken et al., 2022). The ARI ranged from 1 to −1, where 1 indicated perfect concordance, 0 matched random expectation, and negative values implied anti-correlation; computational complexity scaled as *O*(*n*^2^) due to pairwise comparisons. A principal strength lay in its correction for chance agreement, making it robust for comparing partitions with imbalanced cluster sizes; however, interpretation required caution when ground-truth labels were incomplete or ambiguous, as the metric assumed label noise to be negligible and penalized all deviations equally regardless of biological plausibility.

#### D.2.9. k-nearest neighbor Batch Effect Test

The k-nearest neighbor batch effect test (kBET) evaluates whether batch labels are homogenously distributed across local neighborhoods in a high-dimensional embedding, thereby quantifying the presence of residual batch effects after integration. To compute kBET (Bü ttner et al., 2018; Luecken et al., 2022), a k-nearest-neighbor (kNN) graph is constructed in a principal component analysis (PCA)-reduced space using Euclidean distances. For each cell *n*, the algorithm identifies its *k* nearest neighbors and computes the observed batch proportion vector **p**_*n*_ ∈ ℝ^|*B*|^, where |*B*| is the number of batches. Under the null hypothesis of batch homogeneity, the expected proportion *q*_*j*_ for batch *j* equals its global prevalence. A chi-square test compares **p**_*n*_ to **q** via the statistic

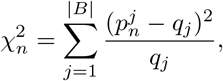

yielding a p-value *p*_*n*_ per cell. The final kBET score aggregates the fraction of cells where *p*_*n*_ exceeds a significance threshold *α* (typically 0.05):

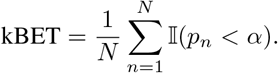

Scores range from 0 to 1, where 0 indicates perfect batch mixing (all p-values ≥ *α*) and 1 signifies severe batch effects (all p-values *< α*). Computational complexity is dominated by kNN graph construction, which is *O*(*N* ^2^) in the worst case but *O*(*N* log *N*) with approximate methods. A key strength is kBET’s statistical rigor, as it formally tests the null hypothesis of batch independence while accounting for dataset-wide batch proportions. However, performance depends on the choice of *k*, which must balance locality preservation and statistical power; small *k* increases variance, while large *k* dilutes local signal. Additionally, sparse batches may violate chi-square assumptions, necessitating permutation tests. The implementation is available at github.com/theislab/kBET.

#### D.2.10. Graph Connectivity

The graph connectivity metric evaluates whether cells sharing a biological identity form a coherently connected neighborhood in the k-nearest neighbor (kNN) graph constructed from the integrated data. For each cell identity class *c* ∈ *C*, a subset kNN graph *G*(*N*_*c*_, *E*_*c*_) was constructed using only cells annotated with *c*. The score was computed as the mean ratio of the largest connected component (LCC) size to the total number of nodes in each class-specific subgraph (Luecken et al., 2022), formally expressed as:

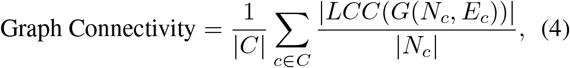

where |*LCC*(*G*(*N*_*c*_, *E*_*c*_)) | denotes the node count of the largest connected component for class *c* and |*N*_*c*_| is the total number of cells with identity *c*. The metric ranges from 0 to 1, where 1 indicates perfect intra-class connectivity (all cells of the same type form a single connected component) and 0 signifies complete fragmentation; values closer to 1 are desirable. Computational complexity is dominated by kNN graph construction, which scales as *O* (*N* ^2^) in the number of cells *N*, rendering the metric costly for large datasets. A principal strength of this metric is its direct assessment of topological integrity for cell-type neighborhoods, critical for graph-based downstream analyses such as clustering or trajectory inference. However, its reliance on pairwise distances for kNN graph construction introduces scalability limitations, particularly when applied to datasets with *N >* 10^5^ cells.

#### D.2.11. Principal Component Regression

The principal component regression (PCR) score assessed the extent to which batch effects explain variance in the integrated data, addressing whether technical variability persists across principal components. The metric decomposes the total variance attributable to batch effects by computing, for each principal component (PC), the product of its explained variance and the coefficient of determination (*R*^2^) from a linear regression of the batch variable onto that PC (Luecken et al., 2022). Formally, given a data matrix *C* and batch variable *B*, the batch-associated variance Var(*C*|*B*) is defined as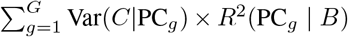, where Var(*C*|PC_*g*_) is the variance explained by the *g*-th PC and *R*^2^(PC_*g*_ |*B*) quantifies the linear dependence between the PC and *B*. PCR values range between 0 and 1, with higher scores indicating greater residual batch effects; optimal integration minimizes this score. Computational complexity is dominated by eigendecomposition during PCA, scaling as *O*(min(*N, e*)^3^) for *N* cells and *e* features, followed by *O*(*GN*) operations for *G* linear regressions. A key strength lies in its interpretability as a weighted variance decomposition, enabling direct comparison of batch effects across integration methods. However, PCR assumes linear relationships between batch variables and latent spaces, limiting its sensitivity to non-linear batch effects that manifest in higher-order interactions or manifold structures.

#### D.2.12. Local Inverse Simpson’s Index (cLISI/iLISI)

The cell-type Local Inverse Simpson’s Index (cLISI) evaluates whether distinct biological identities are preserved as separable neighborhoods in integrated data, answering if local cell-type homogeneity is maintained; the integration Local Inverse Simpson’s Index (iLISI) quantifies the mixing of batches within local neighborhoods, assessing if technical confounders are mitigated. For each cell, a k-nearest-neighbor (kNN) graph was constructed, and the inverse Simpson’s index was computed over the categorical distribution (cell types for cLISI, batches for iLISI) within each neighborhood. Let *N*_*C*_ and *N*_*B*_ denote the number of cell types and batches, respectively. For cLISI, the diversity score 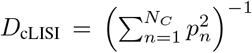 was computed, where *p*_*n*_ is the proportion of the *n*-th cell type; analogously, iLISI used 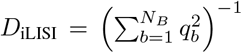, where *q*_b_ is the proportion of the *b*-th batch. Scores were rescaled from the raw range [1, *N*] (where *N* is *N*_*C*_ for cLISI and *N*_*B*_ for iLISI) to [0, 1] via affine transformation, with 0 indicating minimal separation (cLISI) or mixing (iLISI) and 1 denoting optimal preservation or integration. Computationally, both metrics incur *O*(*Nk*) complexity for neighborhood retrieval and diversity calculation across *N* cells, assuming *k* neighbors per cell. High cLISI scores (near 1) reflect neighborhoods dominated by a single cell type, confirming bio conservation; high iLISI scores (near 1) indicate neighborhoods with balanced batch representation, signifying technical integration. The inverse scaling between cLISI and iLISI necessitates joint optimization to balance biological fidelity and batch correction. A principal strength lies in their unified graph-based framework, enabling direct comparison across integration methods (Korsunsky et al., 2019a; Luecken et al., 2022); however, performance is sensitive to the choice of *k*, assumes local neighborhoods are representative of global structure, and may conflate biological and technical variation when batches are confounded with cell types.

#### D.2.13. Aggregate Scores

The `scIB` framework synthesizes the individual quality metrics described above into two domain-specific subscores— one for *bio conservation* and one for *batch correction*—and finally into a single *overall integration score*. This composite view enables rapid, quantitative ranking of integration methods without manually weighing every metric (Luecken et al., 2022).

Step 1: Normalizing each metric to [0, 1]: Because the raw metrics differ in scale, range, and directionality (i.e., some metrics improve as their values increase, while others improve as values decrease), each metric *m*_*k*_ is first rescaled to a common unit interval [0, 1] so that higher values consistently indicate better performance. This rescaled version is denoted 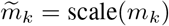, where the scaling function is typically a min–max or quantile transformation chosen to preserve monotonicity in performance. For example, metrics that naturally increase with better integration—such as ARI, NMI, celltypeASW, or cLISI—are directly mapped linearly to [0, 1]. Conversely, metrics that decrease with better integration—such as batchASW, kBET, or PCR—are first inverted (e.g., 1 *− m*_*k*_) before applying the same normalization. This ensures that all normalized metrics 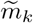 can be meaningfully aggregated without introducing bias from inconsistent scaling or directionality.

Step 2: Computing domain-specific subscores: Each normalized metric is assigned to one of two categories: bio conservation or batch correction. The set of biological metrics includes celltypeASW, cLISI, ARI, NMI, and isolatedlabel-F1, while the batch metrics include batchASW, iLISI, kBET, PCR, and graph-connectivity. Letting 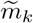 denote the normalized value of metric *k*, the bio-conservation ological set, and the batch correction score as the average over the batch set. Formally, this yields a bio score is computed as the average of all 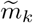 in the biscore of 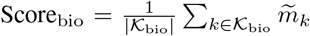 and a batch score of 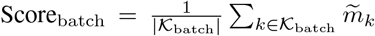, where *K*_bio_ and *K*_batch_ index the relevant metric sets. Both scores range from 0 to 1, with higher values reflecting better bio conservation or more effective batch mixing, respectively.

Step 3: Deriving the overall integration score: By default, `scIB` computes the overall integration score as the arithmetic mean of the bio conservation and batch correction subscores, i.e., Score_overall_ = (Score_bio_ + Score_batch_)*/*2, which yields a value between 0 and 1. A value near 1 indicates that the integration method has successfully preserved biological structure while removing technical batch effects. Conversely, a value near 0 suggests that the method failed in at least one of these dimensions. Although equal weighting is used by default, this balance can be adjusted in domain-specific contexts—for example, giving more weight to Score_bio_ in lineage-tracing studies, or to Score_batch_ in highly heterogeneous multi-batch datasets.

Implementation and practical caveats: The normalization of individual metrics within `scIB` requires careful consideration to ensure comparability and interpretability across diverse scales and behaviors. A common approach is min– max scaling, which maps metric values to the unit interval [0, 1] based on global extrema; however, this method can be unstable if outlier methods produce extreme values that distort the scale. To address this, quantile-based or robust scaling transformations are often preferred, as they are less sensitive to outliers and heavy-tailed distributions. Beyond normalization, the method of aggregating scores also affects interpretability. While direct averaging preserves the magnitude of each normalized metric *m*, it can be skewed by high-variance metrics or irregular distributions. An alternative is to convert each *m*_*k*_ to a percentile rank before averaging, yielding a rank-based aggregate score that is less susceptible to extreme values and enhances robustness. A further consideration arises from the reliance of certain biological metrics—such as ARI, NMI, and celltypeASW—on ground-truth cell-type annotations. Incomplete, noisy, or inconsistent labels can artificially inflate or deflate Score_bio_, thereby undermining the validity of comparisons across integration methods. Finally, a core challenge in evaluating data integration lies in the inherent trade-off between preserving biological structure and removing technical artifacts. Methods that prioritize batch removal may inadvertently obscure meaningful biological variation, resulting in a high Score_batch_ but a low Score_bio_; conversely, conservative approaches that preserve subtle cell-state differences may under-correct for batch effects. The overall integration score in `scIB` exposes this tension explicitly, encouraging users to interpret composite performance in light of domain-specific goals and dataset characteristics. In summary, the composite scoring scheme in `scIB` provides an interpretable, single-number summary while retaining the granularity of domain-specific subscores. Users can therefore *(i)* rank competing integration pipelines quickly, *(ii)* inspect whether deficiencies stem from biological distortion or residual batch effects, and *(iii)* tune parameters or weights according to study-specific priorities.

### D.3. Metric Descriptions for Trajectory Evaluation

#### Group 1

1. Concordance Index
2. Wasserstein Distance for Pseudotime
3. Dynamic Time Warping Distance
4. R-Squared with Spline
5. R-Squared
6. Mean Absolute Error
7. Mean Squared Error
8. Pearson Correlation
9. Spearman Correlation
10. Kendall’s *τ* Correlation
11. Normalized Mutual Information
12. Mutual Information (KDE)
13. CDF Cramér–von Mises
14. Kolmogorov–Smirnov Statistic
15. 15. Geary’s *C* for Pseudotime
16. Moran’s *I* for Pseudotime

#### Group 2

1. Graph Edit Distance
2. Frobenius
3. Accuracy
4. *L*_1_ Norm
5. Spectral Distance
6. Hamming Distance
7. Jaccard Similarity
8. Precision
9. Recall
10. F1 Score
11. Mantel Correlation
12. Average Shortest Path Difference
13. Laplacian Spectral EMD
14. Permutation-Marginalized SSIM
15. Clustering Coefficient Difference
16. Persistence Diagram Distance

#### Group 3

1. Branch Silhouette Score
2. Normalized Mean Curvature Score
3. Moran’s *I* for Embedding
4. Geary’s *C* for Embedding
5. Embedding Distance Correlation
6. Sammon’s Stress
7. Graph-Based Trustworthiness
8. Neighborhood Preservation Score
9. Directionality Preservation
10. Wasserstein Distance for Embedding
11. Directed Trajectory Validation Score

#### D.3.1. Concordance Index

The Concordance Index (CI) evaluates the probability that the inferred pseudotime ordering of cell states matches the reference trajectory’s temporal progression (Longato et al., 2020; Steck et al., 2007). It operates on pseudotime vectors (*p*_ref_, *p*_inf_), quantifying pairwise order agreement through normalized counts of concordant and tied pairs. For each admissible pair of cells with distinct reference pseudotimes, the product of pseudotime differences in both vectors was computed; concordant pairs (positive product) contributed fully to the score, while ties in the inferred pseudotime contributed partially. The CI equals the ratio of such contributions to all admissible pairs, equivalent to the area under the ROC curve for pairwise comparisons. Scores range from 0 (perfect discordance) to 1 (perfect concordance), with 0.5 indicating random agreement; higher values denote better preservation of the reference ordering. The computation requires *O*(*n*^2^) time due to pairwise comparisons, which may be prohibitive for large *n*. A principal strength is its non-parametric nature and global assessment of ordering fidelity; however, exclusion of tied reference pairs and sensitivity to inferred ties may limit interpretability when pseudotimes are insufficiently resolved.

#### D.3.2. Wasserstein Distance for Pseudotime

The Wasserstein distance quantifies the minimal transport cost required to align the inferred pseudotime distribution with the reference distribution, addressing whether the inferred temporal progression preserves the global statistical properties—such as central tendency and dispersion—of cellular developmental timing (Panaretos & Zemel, 2019; Piccoli & Rossi, 2016). The metric operates on pseudotime vectors (*p*_ref_, *p*_inf_), computed as the first Wasserstein distance between their empirical distributions: 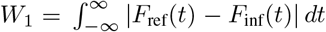, where *F*_ref_ and *F*_inf_ are cumula-tive distribution functions. This measures the minimal work needed to morph one distribution into the other, integrating over all quantile mismatches. Scores are non-negative, with 0 indicating identical distributions and higher values reflecting increasing distributional divergence; magnitude scales with the pseudotime axis. Computational complexity is *O*(*n* log *n*) due to sorting for quantile alignment. The metric’s principal strength lies in its sensitivity to both location shifts (e.g., systematic temporal delays) and dispersion differences (e.g., compressed or expanded pseudotime scales); however, it assumes comparable distribution supports and may obscure local temporal inversions, as it aggregates discrepancies across the entire pseudotime domain.

#### D.3.3 Dynamic Time Warping Distance

The Dynamic Time Warping (DTW) distance quantifies the minimum cumulative alignment cost between reference and inferred pseudotime sequences under temporal warping, addressing whether the inferred trajectory preserves temporal progression patterns despite local distortions in timing or velocity (Berndt & Clifford, 1994; Zhao & Itti, 2018). The metric operates on min-max normalized pseudotime vectors (*p*_ref_, *p*_inf_), computed by finding the optimal warping path that minimizes the sum of Euclidean distances between aligned elements, normalized by the path length to ensure scale invariance. Let *P* denote the set of all admissible warping paths; the DTW distance is 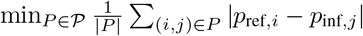, where |*P* | is the path length. Scores are non-negative, with lower values indicating better alignment; a value of 0 corresponds to identical sequences, while higher values reflect increasing temporal distortion. The FastDTW algorithm (Wu & Keogh, 2020) reduces computational complexity to *O*(*n*) via a multilevel approach with a search radius heuristic, contrasting with the *O*(*n*^2^) complexity of exact DTW. The metric’s principal strength is its invariance to nonlinear temporal warping, making it suitable for evaluating trajectories with variable progression rates; however, sensitivity to pseudotime noise and dependence on the radius parameter may obscure biologically meaningful local variations or over-smooth subtle temporal shifts.

#### D.3.4. R-squared with Spline

The R-squared with Spline metric evaluates the proportion of variance in the reference pseudotime vector explainable by a flexible nonlinear function of the inferred pseudotime, addressing whether the inferred trajectory captures systematic—potentially nonlinear—temporal relationships between cell states. The metric operates on pseudotime vectors (*p*_ref_, *p*_inf_), first sorting both vectors by the inferred pseudotime to enforce monotonicity, then fitting a univariate spline of degree *k* to model *p*_ref_ as a smoothed function of *p*_inf_ (Ratner, 2017; Hastie, 2017). The coefficient of determination was computed as 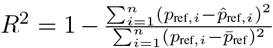, where 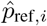 denotes spline-predicted values and 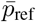 the mean reference pseudotime. Scores range (, 1], with 1 indicating perfect explainability via the spline, 0 implying no improvement over the mean reference, and negative values indicating worse fit; higher values reflect stronger nonlinear correspondence. Computational complexity is *O*(*n* log *n*) for sorting and *O*(*n*) for spline fitting, assuming efficient implementations. The metric’s principal strength is its capacity to detect nonlinear monotonic associations beyond linear correlation, leveraging spline flexibility; however, results depend critically on the spline’s degree and smoothing hyperparameters, while sorting by inferred pseudotime assumes temporal coherence, potentially misrepresenting trajectories with local ordering inversions or non-monotonic relationships.

#### D.3.5. R-squared

The R-squared coefficient quantifies the proportion of variance in the reference pseudotime vector explained by the inferred pseudotime progression, addressing whether temporal distances between cell states in the inferred trajectory systematically account for variations observed in the ground truth (Ratner, 2017). The metric operates on pseudotime vectors (*p* _ref_, *p*_inf_), computed as 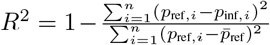 where 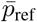 is the mean reference pseudotime, representing the ratio of unexplained variance to total variance. Scores range from to 1, where 1 indicates perfect linear explainability, 0 implies no linear predictive power beyond the mean reference value, and negative values denote worse performance than a constant model; higher values reflect stronger linear correspondence. Computational complexity is *O*(*n*) from mean and variance calculations. The metric’s strength lies in its interpretability as a normalized measure of linear association strength, directly aligned with regression analysis conventions; however, it assumes linearity and homoscedasticity, rendering it insensitive to nonlinear relationships, while negative scores complicate interpretation in contexts where inferred trajectories are expected to be structurally congruent with the reference.

#### D.3.6. Mean Absolute Error

The Mean Absolute Error (MAE) measures the average magnitude discrepancy between reference and inferred pseudotime vectors, addressing whether the inferred temporal progression preserves the absolute distances of cell states from the root relative to the ground truth (Hodson, 2022; Qi et al., 2020). The metric operates on pseudotime vectors (*p*_ref_, *p*_inf_), first normalized to [0, 1] via minmax scaling to ensure scale invariance, then computed as 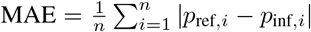, representing the expected absolute deviation across cells. Scores range in [0, 1], with 0 indicating perfect alignment of scaled pseudotimes and higher values reflecting linearly increasing average error magnitude; unlike squared error metrics, MAE weights all deviations proportionally. Computational complexity is *O*(*n*) from element-wise differences and summation. Its principal strength lies in robustness to outliers and interpretability as a scale-invariant measure of global temporal distortion, suitable for quantifying systematic biases; however, linear error weighting may underestimate the impact of localized severe inaccuracies, while normalization precludes assessment of absolute pseudotime scaling fidelity.

#### D.3.7. Mean Squared Error

The Mean Squared Error (MSE) quantifies the average squared deviation between reference and inferred pseudo-time vectors, evaluating how accurately the inferred temporal progression preserves the absolute magnitude differences of cell states relative to the reference trajectory (Hodson, 2022). The metric operates on pseudotime vectors (*p*_ref_, *p*_inf_), first normalized to [0, 1] via min-max scaling to ensure scale invariance, then computed as 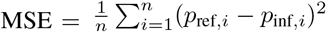, representing the expected squared error over all cells. Scores are non-negative, with 0 indicating perfect alignment in scaled pseudotimes and higher values reflecting increasing average error magnitude; the squaring penalizes large deviations disproportionately. Computational complexity is *O*(*n*) from element-wise operations and summation. Its principal strength lies in providing an interpretable, scale-invariant measure of global temporal distortion magnitude, sensitive to both systematic biases and localized inaccuracies; however, sensitivity to outliers and quadratic error weighting may overemphasize rare but severe discrepancies, while normalization precludes direct interpretation of absolute pseudotime differences.

#### D.3.8. Pearson Correlation

The Pearson correlation coefficient quantifies the degree of linear correspondence between reference and inferred pseudotime orderings, addressing whether progression along the inferred trajectory preserves relative temporal distances between cell states proportionally (Essam et al., 2022; van den Heuvel & Zhan, 2022). It operates on pseudotime vectors (*p*_ref_, *p*_inf_), computed as the covariance of the paired vectors normalized by the product of their standard deviations: 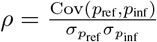. This measures linear association through the ratio of their joint variability to individual variabilities, with perfect linearity yielding |*ρ*| = 1 and complete independence yielding *ρ* = 0. Scores range in [−1, 1], where 1 indicates identical linear ordering, -1 denotes perfect inverse correlation, and magnitudes reflect effect sizes; values near 0 suggest negligible linear relationship, though nonlinear associations may persist. Computational complexity is *O*(*n*) from means, variances, and covariance calculations. Its principal strength lies in interpretability as a normalized measure of proportional temporal agreement, sensitive to both ranking and scale alignment; however, sensitivity to outliers and restriction to linear relationships limit utility when pseudotime distortions involve nonlinear monotonic transformations or localized perturbations.

### D.3.9. Spearman Correlation

The Spearman rank correlation coefficient assesses whether the inferred pseudotime progression preserves the monotonic ordering of cell states relative to the reference trajectory, answering whether temporal relationships between cells are consistently ranked irrespective of absolute temporal distances (Essam et al., 2022; van den Heuvel & Zhan, 2022). The metric operates on pseudotime vectors (*p*_ref_, *p*_inf_), computed by replacing each pseudotime value with its rank within the respective vector and then calculating the Pearson correlation between the rank-transformed 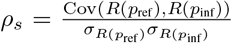, where *R*(*·*) denotes rank assignment. This non-parametric approach measures the strength and direction of monotonic associations, invariant to nonlinear scaling. Scores range in [−1, 1], with 1 indicating perfect rank agreement, -1 denoting perfect inversion, and 0 implying no monotonic relationship; values near extremes suggest robust preservation or reversal of temporal hierarchy. Computational complexity is *O*(*n* log *n*) due to ranking, dominated by sorting operations. The metric’s principal strength lies in robustness to non-Gaussian noise and outliers, capturing nonlinear but order-preserving pseudotime distortions; however, it discards information about relative pseudotime magnitudes between cells and may misrepresent trajectories with frequent tied ranks, as tied values are averaged during ranking, potentially diluting sensitivity to local order perturbations.

#### D.3.10. Kendall’s Tau Correlation

Kendall’s tau correlation coefficient evaluates the agreement in pairwise ordering between reference and inferred pseudotime vectors, addressing whether the inferred trajectory preserves the relative temporal hierarchy of cell states as defined by their concordant or discordant progression orders (Essam et al., 2022; van den Heuvel & Zhan, 2022). The metric operates on pseudotime vectors (*p*_ref_, *p*_inf_), computed as the normalized difference between the number of concordant and discordant cell pairs: 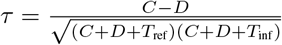 where *C* and *D* denote concordant (same order in both vectors) and discordant (opposite order) pairs, while *T*_ref_ and *T*_inf_ account for ties in each vector. This non-parametric measure quantifies the probability that the inferred pseudotime ordering aligns with the reference, invariant to monotonic transformations. Scores range in [− 1, 1], with 1 indicating perfect ordinal agreement, -1 complete inversion, and 0 statistical independence; magnitudes reflect the strength of association, where |*τ*| *>* 0.5 suggests strong concordance. Computational complexity is *O*(*n*^2^) due to pairwise comparisons, limiting scalability for large *n*. Its principal strength lies in robustness to non-linearities and distributional assumptions, providing a distribution-free assessment of rank correspondence; however, sensitivity diminishes in the presence of frequent tied ranks, common in discretized or coarsely resolved pseudotime estimates, and computational cost becomes prohibitive for datasets exceeding 10^4^ cells.

#### D.3.11. Normalized Mutual Information

The Normalized Mutual Information (NMI) quantifies the statistical dependence between reference and inferred pseudotime distributions, addressing whether the inferred trajectory captures non-linear and non-monotonic patterns in cellular progression timing beyond rank or linear correlations (Kvålseth, 2017; Belghazi et al., 2018; Duncan, 1970; Kraskov et al., 2004; Vinh et al., 2010). The metric operates on *z*-score normalized pseudotime vectors (*p*_ref_, *p*_inf_), computed by adaptively discretizing both vectors into bins determined by the Freedman-Diaconis rule—optimizing for data spread while handling zero-variance cases (Freedman & Diaconis, 1981)—then calculating the mutual information between discretized distributions normalized by their average entropy: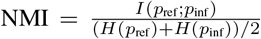, where *I* denotes mutual information and *H* entropy. Scores range in [0, 1], with 1 indicating perfect dependency (including inverted or nonlinearly transformed orderings) and 0 statistical independence; intermediate values reflect the strength of distributional alignment. Computational complexity is *O*(*n* log *n*) from sorting and adaptive binning. The metric’s principal strength lies in invariance to monotonic transformations and sensitivity to arbitrary distributional correspondences; however, dependence on discretization granularity and insensitivity to directional relationships limit its utility for assessing temporal coherence, as identical scores may arise from progression-preserving versus scrambled orderings with equivalent dependency strength.

#### D.3.12. Mutual Information KDE

The Mutual Information (MI) KDE metric quantifies the statistical dependence between reference and inferred pseudotime vectors, addressing whether cellular progression timing in the inferred trajectory preserves non-linear, non-monotonic associations with the ground-truth temporal ordering. The metric operates on *z*-score standardized pseudotime vectors (*p*_ref_, *p*_inf_), computing their mutual information via kernel density estimation (KDE) with covariance regularization and data whitening (Weglarczyk, 2018; Zandieh et al., 2023; Tong et al., 2025). After standardization, joint and marginal probability densities were estimated using Gaussian KDEs with bandwidths optimized via the Scott rule (Scott, 1979), stabilized through Cholesky decomposition of the regularized covariance matrix and whitening to prevent numerical instability. The MI was calculated as MI(*p*_ref_, *p*_inf_) = E; [log(*p*_*XY*_ (*x, y*)*/*(*p*_*X*_ (*x*)*p*_*Y*_ (*y*))], where *p*_*XY*_ and *p*_*X*_, *p*_*Y*_ denote joint and marginal densities estimated from whitened data, adjusted by the Jacobian determinant of the whitening transform (Duncan, 1970; Liao & He, 2021; Kraskov et al., 2004). The score ranges from 0 (statistical independence) to ∞, with higher values indicating stronger dependence; interpretation relies on relative comparisons due to the unbounded range. Computational complexity is *O*(*n*^2^) from pairwise KDE evaluations, limiting scalability to very large datasets. A principal strength lies in non-parametric capture of arbitrary dependencies, including non-linear and non-functional associations undetectable by correlation-based metrics; however, estimator bias increases for small sample sizes (*n <* 100), and results depend on bandwidth selection, requiring standardized inputs to ensure comparability across systems.

#### D.3.13. CDF Cramér-von Mises

The Cramér-von Mises (CvM) statistic evaluates whether the inferred pseudotime distribution globally aligns with the reference, answering whether cellular progression timing is captured without systematic bias across the entire trajectory. It operates on normalized pseudotime vectors (*p*_ref_, *p*_inf_), first min-max scaling each to [0, 1] to mitigate scale discrepancies, then computing the integrated squared difference between their empirical cumulative distribution functions (ECDFs) via the two-sample Cramér-von Mises test (Genest et al., 2006; 2007). Let *F*_ref_ and *F*_inf_ denote the ECDFs; the CvM statistic aggregates (*F*_inf_(*x*) − *F*_ref_(*x*))^2^ over all observed *x*, weighted by the pooled sample distribution. The result is a non-negative score where 0 indicates identical ECDFs and higher values reflect increasing divergence; its magnitude depends on both distributional shape and spread. Computational complexity is *O*(*n* log *n*) due to ECDF sorting, where *n* is the number of cells. A principal strength is its non-parametric, holistic sensitivity to any deviation in cumulative distribution, making it robust to non-monotonic or nonlinear temporal distortions; however, the statistic conflates biologically meaningful temporal shifts with stochastic noise and requires sufficient cell counts to reliably estimate ECDFs, potentially overpenalizing minor discrepancies in small datasets.

#### D.3.14. Kolmogorov-Smirnov Statistic

The Kolmogorov-Smirnov (KS) statistic evaluates the maximum discrepancy between the empirical cumulative distribution functions (CDFs) of reference and inferred pseudotime vectors, addressing whether the inferred trajectory recapitulates the global temporal distribution of cell states in *G*_ref_ (Lopes et al., 2007; Steinskog et al., 2007; Drezner et al., 2010; Razali et al., 2011). The metric operates on min-max normalized pseudotime vectors (*p*_ref_, *p*_inf_) ∈ [0, 1]^*n*^ × [0, 1]^*n*^, where normalization ensures scale invariance. The KS statistic was computed as the supremum of absolute differences between the CDFs of *p*_ref_ and *p*_inf_: *D* = sup_*x∈*[0,1]_ |*F*_ref_(*x*) − *F*_inf_(*x*) |, where *F*_ref_ and *F*_inf_ denote the empirical CDFs. The score ranges in [0, 1], with 0 indicating identical CDFs and higher values reflecting increasing divergence; values approaching 1 signify maximal distributional mismatch. Computational complexity is *O*(*n* log *n*) due to sorting operations for CDF estimation. A principal strength is the metric’s non-parametric nature, requiring no assumptions about distributional form while being sensitive to point-wise differences; however, sensitivity is typically greatest near the median rather than distribution tails, and small sample sizes may reduce reliability, as finite-sample CDF estimates become less precise.

#### D.3.15. Geary’s C for Pseudotime

Geary’s C statistic evaluates whether cells proximal in the reference trajectory graph exhibit coherent pseudotimes in the inferred trajectory, addressing whether local temporal coherence is preserved relative to the reference topology (Anselin, 1995; DeTomaso & Yosef, 2021; DeTomaso et al., 2019; de Jong et al., 1984). The metric operates on the reference adjacency matrix *A*_ref_ and inferred pseudotime vector *p*_inf_, first standardizing *p*_inf_ to zero mean and unit variance to isolate spatial patterns. A cell-cell adjacency matrix was derived from *A*_ref_ by propagating inter-cell-type connections to individual cells, then normalized to row-stochasticity. Geary’s C was computed as 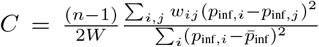, where *w*_*ij*_ denotes adjacency weights, *W* their global sum, and 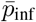 the mean pseudotime. Scores range [0, 2], where values below 1 indicate positive spatial autocorrelation (neighboring cells have similar pseudotimes), 1 implies randomness, and values above 1 suggest negative autocorrelation (neighbors diverge). Computational complexity is dominated by sparse matrix operations, scaling linearly with the number of non-zero edges in *A*_ref_. A principal strength is sensitivity to local pseudotime discrepancies, capturing fragmented or inverted progression along reference branches; however, dependence on the spatial weights matrix necessitates careful normalization, and the statistic conflates topological inaccuracies with pseudotime noise, potentially overpenalizing biologically plausible temporal heterogeneity within connected regions.

#### D.3.16. Moran’s I for Pseudotime

The Moran’s I metric evaluates whether cells adjacent in the reference trajectory exhibit spatial autocorrelation in their inferred pseudotime, testing if locally connected cell states share similar progression timing (de Jong et al., 1984; Tiefelsdorf & Boots, 1995; Van den Berge et al., 2020). The score operates on the reference cell-type adjacency matrix *A*_ref_ and inferred pseudotime vector *p*_inf_, first expanding *A*_ref_ into a cell-cell adjacency matrix by connecting cells sharing adjacent types, then computing Moran’s I as 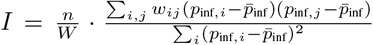, where *n* is the cell count, *w*_*ij*_ are adjacency-derived weights, *W* their global sum, and 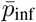 the mean pseudotime. The statistic ranges [*™*1, 1], with *I* ≈ −1 indicating strong positive autocorrelation (adjacent cells share similar pseudotimes), *I* ≈−1 dispersion, and *I* ≈ 0 spatial randomness; higher values denote better coherence with the reference topology. Computed in *O*(*n*) time via sparse linear algebra, the metric’s principal strength is its direct assessment of pseudotemporal consistency with the reference graph’s local structure. However, it disregards edge directionality and depends critically on the reference adjacency’s construction, potentially conflating biological proximity with graph abstraction artifacts.

#### D.3.17. Graph Edit Distance

The Graph Edit Distance (GED) quantifies the structural dissimilarity between the reference and inferred trajectory graphs by computing the minimum number of edge additions or deletions required to transform one graph into the other (Gao et al., 2010; Blumenthal et al., 2020; Abu-Aisheh et al., 2015). The metric operates on the binary adjacency matrices (*A*_ref_, *A*_inf_), where *A*_inf_ is obtained by thresholding the inferred inter-type connectivity matrix. After excluding self-loops, edge sets for both graphs were extracted and their symmetric difference computed, with GED defined as the cardinality of this set, equivalent to the total count of edges present in exactly one graph. The resulting score is a non-negative integer, where 0 indicates isomorphic graphs and higher values reflect increasingly divergent topologies; computational complexity is *O*(*t*^2^) in the number of cell types *t*, due to pairwise edge comparisons. A principal strength lies in its direct interpretation as the minimal edit cost for graph alignment, providing an unambiguous measure of structural fidelity; however, the metric’s quadratic scaling renders it impractical for large *t*, and its sensitivity to thresholding artifacts or minor edge discrepancies may overemphasize topological noise in sparse graphs.

#### D.3.18. Frobenius

The Frobenius metric quantifies the global structural dissimilarity between the reference and inferred cell-type transition graphs, addressing whether the inferred adjacency matrix *W*_inf_ preserves the topological connectivity of the reference graph *A*_ref_. The metric operates on the adjacency matrices (*A*_ref_, *W*_inf_) ∈ {0, 1} ^*t*×*t*^ ×ℝ^*t*×*t*^, computed as the Frobenius norm (Bö ttcher & Wenzel, 2008; Peng et al., 2016) of their element-wise difference matrix: 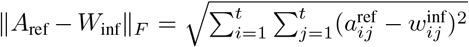. This norm measures the Euclidean distance between vectorized adjacency matrices, providing a holistic assessment of edge agreement across all *t* cell types. The metric yields a non-negative scalar, where 0 indicates identical adjacency structures; higher values reflect increasing discrepancies in edge presence, with upper bounds determined by the number of discordant edges. Computational complexity is *O*(*t*^2^) due to full matrix traversal, remaining tractable for typical cell-type counts *t*. A principal strength lies in its sensitivity to all structural deviations, comprehensively capturing both spurious and missing edges; however, the metric does not localize discrepancies or prioritize biologically critical connections, treating all edge mismatches equally regardless of their position or biological relevance in the trajectory.

#### D.3.19. Accuracy

The Accuracy metric quantifies the global structural congruence between the reference and inferred cell-type adjacency graphs, addressing the question of how precisely both present and absent edges are recovered in the inferred topology (Ratner, 2017; Huffaker et al., 2010). The score operates on the reference binary adjacency matrix *A*_ref_ and the inferred weighted adjacency matrix *W*_inf_, where the latter was binarized via a user-specified threshold *τ* to yield *A*_inf_. Accuracy was computed as the proportion of matching elements between *A*_ref_ and *A*_inf_, equivalent to the ratio of true positives (correctly inferred edges) and true negatives (correctly inferred non-edges) to the total number of possible edges. Scores span [0, 1], with 1 indicating perfect agreement between reference and inferred graphs, 0 complete discordance, and intermediate values reflecting the fraction of correctly recovered edges and non-edges; higher scores denote superior fidelity. The computation requires *O*(*t*^2^) time for *t* cell types, reflecting element-wise comparison of *t* × *t* matrices. A principal strength lies in the metric’s conceptual simplicity and balanced consideration of both edge presence and absence; however, performance is sensitive to threshold selection and may be inflated in sparse graphs where true negatives dominate, while its inability to distinguish between false positive and false negative errors limits granular insight into directional discrepancies in edge inference.

#### D.3.20. L1 Norm

The L1 Norm quantifies the total magnitude of edge discrepancies between the reference and inferred adjacency matrices, addressing how much the inferred graph’s connectivity deviates from the ground truth in terms of edge presence and weight (Wang et al., 2011; Miller et al., 2010). The metric operates on the pair (*A*_ref_, *W*_inf_), where *A*_ref_ is the reference adjacency matrix and *W*_inf_ denotes the inferred weighted adjacency matrix. The score was computed as the *L*_1_ norm of the element-wise difference between the two matrices, ∥*A*_ref_ −*W*_inf_ ∥_1_ = ∑_*i,j*_ |*A*_ref,*ij*_ −*W*_inf,*ij*_ |, summing absolute deviations across all edges. The score is non-negative, with 0 indicating identical adjacency matrices and higher values reflecting cumulative discrepancies; the computational complexity is *O*(*t*^2^) in the number of cell types *t* due to full matrix traversal. A principal strength is the metric’s simplicity and direct interpretability as a total error measure, sensitive to both the number and magnitude of edge-wise differences; however, it uniformly weights discrepancies irrespective of topological context, potentially obscuring the structural impact of specific edge changes and lacking spatial localization within the graph.

#### D.3.21. Spectral Distance

The Spectral Distance assesses topological dissimilarity between the reference and inferred cell-type adjacency graphs by comparing their spectral properties, addressing whether the inferred graph preserves global structural characteristics such as connectivity and expansion (Jovanović & Stanić, 2012; Konukoglu et al., 2012). The metric operates on adjacency matrices (*A*_ref_, *W*_inf_), computing eigenvalues for each and sorting them to account for permutation invariance; the spectral distance was then calculated as the Euclidean norm of the difference between these sorted eigenvalues, *d* = ∥*λ*(*A*_ref_) −*λ*(*W*_inf_) ∥ _2_, where *λ*(·) denotes the sorted eigenvalue spectrum. The score is non-negative, with *d* = 0 indicating identical spectra (isomorphic graphs under the same basis) and higher values reflecting increasing topological divergence; practical upper bounds depend on matrix dimensions and edge densities. Computational complexity is dominated by eigendecomposition at *O*(*t*^3^) for *t* cell types, making it efficient for small *t* but prohibitive for large graphs. A principal strength is sensitivity to global properties like spectral gaps and connected components, which are invariant to local perturbations; however, the metric cannot discern local edge discrepancies and may yield identical scores for non-isomorphic cospectral graphs, which share eigenvalues despite differing topologies.

#### D.3.22. Hamming Distance

The Hamming Distance quantifies the number of edge discrepancies between the inferred and reference cell-type adjacency matrices, answering how topologically faithful the inferred graph is in terms of binary edge presence (Ganagi & Ramane, 2016; Ehounou et al., 2020). The metric operates on the reference adjacency matrix *A*_ref_ and the thresholded inferred adjacency matrix *A*_inf_, computed by binarizing the inferred inter-type connectivity matrix *W*_inf_ using a userspecified threshold. The score was calculated as the elementwise sum of mismatched entries between *A*_ref_ and the binarized *A*_inf_, equivalent to the cardinality of the symmetric difference between their edge sets: ∑_*i,j*_ |*A*_ref,*ij*_ *− A*_inf,*ij*_|. Scores range from 0 (identical graphs) to *t*^2^ (maximally dissimilar graphs), where *t* is the number of cell types; higher values indicate greater structural divergence. The computational complexity is *O*(*t*^2^) due to exhaustive pairwise comparison. A principal strength is its interpretability as a direct count of edge additions or deletions required to reconcile the graphs; however, the metric disregards edge weights, topological context, and hierarchical importance of specific edges, rendering it insensitive to whether discrepancies occur in critical transitions or peripheral connections. Additionally, dependence on the binarization threshold may inflate scores if the inferred graph’s continuous connectivity structure is inadequately discretized.

#### D.3.23. Jaccard Similarity

The Jaccard Similarity assesses the overlap between the edge sets of the reference and inferred cell-type graphs, answering how topologically similar the inferred trajectory is to the ground truth (Sathre et al., 2022; Besta et al., 2020). It operates on the reference adjacency matrix *A*_ref_ and the inferred weighted adjacency matrix *W*_inf_, with the latter binarized via a user-defined threshold *τ* to yield *A*_inf_. The score was computed as the ratio |ℰ_ref_ ∩ ℰ_inf_| */*|ℰ_ref_ ∪ ℰ_inf_ |, where ℰ_ref_ and ℰ_inf_ denote the edge sets of *A*_ref_ and thresholded *A*_inf_, respectively. Values range from 0 (no common edges) to 1 (identical edge sets), with higher scores indicating stronger topological agreement; the worst-case time complexity is *O*(*t*^2^) for *t* cell types, proportional to the number of edges in practice. A principal strength is its intuitive interpretation as the fraction of shared edges, providing a direct measure of structural fidelity; limitations include sensitivity to the threshold *τ*, disregard for edge weights and directionality, and potential misestimation when edge multiplicities or graded connectivity inform trajectory validity.

#### D.3.24. Precision

The Precision metric evaluates the proportion of inferred edges that are present in the reference trajectory, addressing whether the inferred graph avoids spurious transitions not supported by the ground truth (James et al., 2013; Sokolova & Lapalme, 2009; Powers, 2020; Besta et al., 2020). The score operates on the reference adjacency matrix *A*_ref_ and the inferred weighted adjacency matrix *W*_inf_, which was thresholded at a user-specified value *τ* to yield a binary adjacency matrix *A*_inf_ = п (*W*_inf_ ≥*τ*). Precision was computed as the ratio of true positives (edges present in both *A*_ref_ and *A*_inf_) to predicted positives (all edges in *A*_inf_), formalized as precision = ∥*A*_ref_ ∘*A*_inf_∥ _1_ */* ∥*A*_inf_∥ _1_, where ∘ denotes element-wise multiplication and ∥·∥_1_ the L1 norm. Scores range in [0, 1], with 1 indicating all inferred edges are correct and 0 indicating none are present in the reference; higher values denote better specificity in edge prediction. The computation requires *O*(*t*^2^) time for *t* cell types due to pairwise matrix comparisons. A principal strength is its direct assessment of edge prediction reliability, critical for avoiding false-positive transitions in downstream analyses; however, precision does not penalize false negatives and can be artificially inflated in sparse graphs by conservatively predicting few edges, while threshold sensitivity necessitates careful parameterization to balance specificity and recall.

#### D.3.25. Recall

The recall metric evaluates the proportion of true edges in the reference trajectory graph *G*_ref_ that are successfully recovered by the inferred adjacency structure, addressing whether critical state transitions in the biological process are preserved (James et al., 2013; Sokolova & Lapalme, 2009; Powers, 2020; Besta et al., 2020). The score operates on the reference adjacency matrix *A*_ref_ and the inferred adjacency matrix *W*_inf_ ∈ ℝ^*t*×*t*^, which contains continuous edge weights. To compute recall, *W*_inf_ was first binarized via a user-defined threshold *τ*, yielding *A*_inf_ = I(*W*_inf_ *≥ τ*); true positives were counted as ∑_*i,j*_ п (*A*_ref,*ij*_ = 1 ∧ *A*_inf,*ij*_ = 1), while actual positives corresponded to ∑_*i,j*_ *A*_ref,*ij*_. The re-call score was then calculated as the ratio of true positives to actual positives, equivalent to TP*/*(TP + FN) in binary classification. Scores range from 0 to 1, where 1 indicates all reference edges were recovered and 0 signifies complete failure to detect true transitions; higher values denote superior edge recovery. With *t* cell types, the computation requires *O*(*t*^2^) time due to pairwise comparisons across the adjacency matrices. A principal strength is its direct interpretability for assessing structural fidelity, as missing edges directly imply overlooked biological transitions; however, the metric disregards false positives entirely and is sensitive to the choice of *τ*, which may inflate scores if overly permissive or underestimate performance if too stringent.

#### D.3.26.F1 Score

The F1 Score evaluates the accuracy of edge prediction in the inferred adjacency matrix relative to the reference graph, addressing whether the inferred trajectory recovers true cell-type transitions while balancing false positives and false negatives (James et al., 2013; Sokolova & Lapalme, 2009; Powers, 2020; Besta et al., 2020). The metric operates on the binary adjacency matrices (*A*_ref_, *A*_inf_), where the inferred matrix *A*_inf_ was obtained by thresholding a weighted inter-type connectivity matrix. To compute the score, true positives (edges present in both matrices), predicted positives (edges in *A*_inf_), and actual positives (edges in *A*_ref_) were enumerated; precision (fraction of predicted edges that are true) and recall (fraction of true edges recovered) were calculated as precision = TP*/*predicted positives and recall = TP*/*actual positives, respectively. The F1 Score was derived as their harmonic mean: F1 = 2· precision · recall*/*(precision + recall), yielding a value in [0, 1] where higher scores indicate better-balanced edge recovery. The metric’s computational complexity is *O*(*t*^2^) for *t* cell types, dominated by pairwise comparisons across the adjacency matrices. A principal strength is its synthesis of precision and recall into a unified measure, penalizing extremes of over- or under-prediction; however, reliance on a user-defined threshold for binarization and exclusion of true negatives—which may dominate sparse graphs—limit interpretability in scenarios with class imbalance or ambiguous edge weights.

#### D.3.27. Mantel Correlation

The Mantel correlation evaluates whether the inferred cell-type-level graph preserves the global connectivity structure of the reference trajectory by statistically assessing the correlation between their respective shortest-path distance matrices (Legendre & Fortin, 2010; Legendre et al., 2015; Borcard & Legendre, 2012). The metric operates on the reference adjacency matrix *A*_ref_ and the inferred weighted adjacency matrix *W*_inf_. Each adjacency matrix was converted to a graph, and all-pairs shortest path distances were computed using the Floyd-Warshall algorithm (Hougardy, 2010) to derive distance matrices *D*_ref_ and *D*_inf_; the Mantel test then calculated the Spearman rank correlation between the vectorized upper triangles of these matrices, with significance assessed via permutation testing. The resulting correlation coefficient served as the similarity score. The coefficient ranges in [−1, 1], where 1 indicates perfect structural concordance, -1 perfect inverse correlation, and 0 no association; higher values denote greater preservation of the reference’s global connectivity. The Floyd-Warshall step dominates complexity at *O*(*t*^3^) per graph for *t* cell types, making the metric computationally intensive for large *t*. A principal strength is the incorporation of global topological information through shortest-path distances, capturing higher-order structural relationships beyond direct adjacency comparisons; however, the metric requires fully connected graphs and assumes distance matrices meaningfully represent the underlying biological trajectory, potentially failing when graphs contain disconnected components or when shortest paths do not reflect biological progression.

#### D.3.28. Average Shortest Path Difference

The Average Shortest Path Difference evaluates whether the inferred graph preserves the global navigability and efficiency of information flow inherent to the reference trajectory’s connectivity structure (Gonzalez-Escribano et al., 2022; Goldberg & Harrelson, 2005; Mao & Zhang, 2013). The metric operates on adjacency matrices (*A*_ref_, *Z*_inf_), where *Z*_inf_ ∈{*A*_inf_, *W*_inf_} represents the inferred graph’s binary or weighted adjacency structure. For each graph, the average shortest path length—computed via the Floyd-Warshall algorithm (Hougardy, 2010) with *O*(*t*^3^) complexity for *t* cell types—was derived by averaging geodesic distances between all connected node pairs, considering edge weights if present; the absolute difference between these averages quantified structural divergence. Scores are non-negative, with 0 indicating identical global navigability and higher values reflecting increasing discrepancy in connectivity efficiency; the metric is particularly sensitive to topological perturbations that alter critical paths. A principal strength lies in its holistic assessment of graph-wide transport efficiency, capturing deviations in overall connectivity that local edge-wise comparisons may miss; however, the requirement for fully connected graphs, neglect of individual path distributions, and quadratic sensitivity to minor edge modifications that induce disproportionate changes in average path length limit its utility for fragmented or noisy trajectories.

#### D.3.29. Laplacian Spectral EMD

The Laplacian Spectral Earth Mover’s Distance (EMD) quantifies global structural dissimilarity between the reference cell-type adjacency graph and the inferred trajectory’s connectivity by comparing their spectral properties (Dodonova et al., 2016; Gambhir et al., 2025; Aouchiche & Hansen, 2013; Nath & Paul, 2014). This metric evaluates whether the inferred graph preserves the reference’s topological invariants—such as connectivity patterns and community structure—encoded in the eigenvalues of the graph Laplacian. The inputs were the reference adjacency matrix *A*_ref_ ∈ {0, 1} ^*t*×*t*^ and the inferred weighted adjacency matrix *W*_inf_ ∈ ℝ ^*t*×*t*^. For each graph, the graph Laplacian was computed as *L* = *D* − *A*, where *D* is the diagonal degree matrix; the eigenvalues of *L* were extracted, normalized by their sum (equivalent to the trace of *L*), and compared via the Wasserstein distance (refer to D.3.2) to measure the minimal cost of transforming one normalized eigenvalue distribution into the other. The resulting score is non-negative, with 0 indicating identical spectra (isomorphic graphs) and higher values reflecting increasing topological divergence; computational complexity is *O*(*t*^3^) due to eigendecomposition, scaling cubically with the number of cell types *t*. A principal strength is the metric’s sensitivity to global structural properties—including connected components, community organization, and spectral gaps—while remaining agnostic to local edge-wise discrepancies; however, reliance on undirected graphs and prohibitive eigendecomposition costs for large *t* limit applicability to small or densely connected cell-type graphs, and normalization failures may occur if either graph is disconnected or empty.

#### D3.30. Permutation-Marginalized SSIM

The Permutation-Marginalized Structural Similarity Index (SSIM) assesses whether the inferred weighted adjacency matrix *W*_inf_ shares consistent local structural patterns with the reference *A*_ref_ across all possible node orderings, addressing whether their spatial organizations are perceptually similar irrespective of arbitrary cell-type indexing (Brunet et al., 2011; Bakurov et al., 2022). The metric operates on adjacency matrices (*A*_ref_, *W*_inf_), treating them as intensity images where pixel values encode edge weights. To marginalize over label permutations, the SSIM—which compares luminance, contrast, and structure via local windowed statistics—was computed over multiple permutations of rows and columns, either exhaustively (for small *t*) or through Monte Carlo sampling (for large *t*), then averaged. The final score 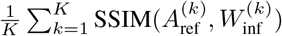 reflects the expected similarity under random node alignments, where 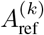 and 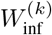 denote the *k*-th permutation. Scores range in [−1, 1], with 1 indicating identical structural patterns, -1 inverse correlation, and 0 no similarity; higher values denote permutation-invariant consistency. Complexity is *O*(*Kt*^2^) for *t* cell types and *K* permutations, manageable for typical *t* but scaling quadratically with *t*. A principal strength is label invariance, critical for graph comparisons where node order carries no semantic meaning; however, reliance on local image-like features may overlook global topological discrepancies, and computational cost grows non-trivially with *t* when exhaustive permutation is required.

#### D.3.31. Clustering Coefficient Difference

The Clustering Coefficient Difference evaluates whether the inferred cell-type adjacency matrix preserves the local clustering structure of the reference graph, specifically addressing the preservation of tightly-knit communities (triangles). It operates on the reference and inferred adjacency matrices (*A*_ref_, *W*_inf_). The score was computed as the absolute difference between the average clustering coefficients of the two graphs, with each coefficient calculated as the mean of node-wise clustering coefficients that quantify the density of triangles incident to each node; edge weights were incorporated to account for connection strength (Watts & Strogatz, 1998; Opsahl & Panzarasa, 2009; Tantardini et al., 2019). The score ranges from 0 (identical local clustering) to 1 (maximal dissimilarity), with higher values indicating divergence in local connectivity patterns; computational complexity is *O*(*t*^3^) for dense graphs but scales with the number of edges in sparse topologies. A principal strength is its sensitivity to local structural perturbations affecting community cohesiveness, such as spurious or missing triangles; however, the metric does not capture global topological discrepancies and may be inflated by edge noise that selectively impacts triangle formation without altering broader connectivity.

#### D.3.32. GDV Similarity

The GDV Similarity metric evaluates the preservation of local connectivity patterns between the reference and inferred cell-type adjacency graphs by comparing their distributions of small connected subgraphs (graphlets). The score answers whether the inferred graph recapitulates the prevalence and configuration of 3-node and 4-node motifs—such as paths, cycles, stars, and triangles—that encode critical biological information about branching, differentiation, and cellular neighborhood organization (Pržulj, 2007; Yaveroğlu et al., 2014; Milenković & Pržulj, 2008). The metric operates on symmetric adjacency matrices (*A*_ref_, *A*_inf_), where *A*_inf_ is obtained by thresholding the inferred inter-type connectivity matrix. For each graph, Graphlet Degree Vectors (GDVs) were computed by enumerating all 3-node and 4-node connected subgraphs, classifying them into 8 predefined types, and counting each node’s participation; this yielded, for each node, an 8-dimensional vector encoding its local topology. Jensen-Shannon Divergence (JSD) was then calculated between the two graphs’ node-wise distributions of counts for each graphlet type, aggregated across types via averaging, and transformed into a similarity score as 1− JSD. The score ranges from 0 (maximal divergence in local structure) to 1 (identical GDV distributions), with higher values indicating better preservation of motif architecture. Computational complexity is dominated by GDV enumeration, scaling as *O*(*t*^4^) for *t* cell types, rendering the metric suitable for moderate-sized graphs but challenging at scale. A principal strength is its sensitivity to local topological patterns that global metrics overlook, providing granular insight into motif conservation; however, reliance on exhaustive subgraph enumeration limits scalability, and the focus on small graphlets may underrepresent higher-order connectivity patterns critical for complex trajectories.

#### D.3.33. Weisfeiler-Lehman Distance

The Weisfeiler-Lehman Distance evaluates whether the inferred adjacency matrix preserves the hierarchical multiscale connectivity patterns of the reference graph by quantifying their structural isomorphism through iterative neighborhood aggregation (Shervashidze et al., 2011; Kriege et al., 2020; Morris et al., 2019). The metric operates on binarized adjacency matrices (*A*_ref_, *A*_inf_), thresholded to discard weak edges and optionally stripped of self-loops. To compute the distance, both matrices were converted into undirected graphs, followed by synchronized Weisfeiler-Lehman (WL) label refinement over *h* iterations—a procedure that augments node labels with compressed multiset representations of their *h*-hop neighborhoods. The WL subtree kernel aggregated counts of matching label sequences across iterations, measuring similarity as the normalized inner product 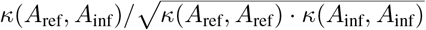, where *κ* denotes the subtree kernel; the final distance was 1 − *κ*_norm_. Scores range in [0, 1], with 0 indicating structural identity and higher values reflecting divergence; complexity is *O*(*ht*^2^) for *t* cell types due to iterative neighborhood aggregation. A principal strength is sensitivity to hierarchical topological patterns invariant to node permutations; however, the distance depends on the binarization threshold and iteration depth, disregards edge weights, and does not localize discrepancies to substructures.

#### D.3.34. GIN GNN Similarity

This metric assesses whether the inferred cell-type adjacency matrix *A*_inf_ preserves the multi-scale topological properties of the reference graph *A*_ref_, addressing structural fidelity through graph neural embeddings. The inputs (*A*_ref_, *A*_inf_) were converted to node feature matrices via 13 topological descriptors—including eigenvector centrality, weighted PageRank (Andersen et al., 2007), and structural core numbers—then processed through a fixed-weight Graph Isomorphism Network (GIN) to generate node embeddings (Xu et al., 2018; Kim & Ye, 2020; Chen et al., 2019; Bai et al., 2019). Node features were aggregated into graph-level embeddings through stabilized moment encoding (mean, variance, skewness, kurtosis), interaction terms (element-wise products, normalized differences), and positional encodings of sorted embeddings; the final similarity was computed as the cosine similarity between graph embeddings: 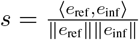, where *e*_ref_, *e*_inf_ denote the aggregated embeddings. Scores range [−1, 1], with 1 indicating identical structural profiles and -1 anticorrelation; higher values reflect stronger multi-scale topological alignment. Computational complexity is dominated by *O*(*t*^3^) eigenvector centrality computations but mitigated by fixed GIN inference. The metric’s principal strength lies in its theoretically grounded use of GIN’s Weisfeiler-Lehman isomorphism expressivity and multi-feature aggregation, enabling sensitivity to both local connectivity patterns and global graph topology; however, reliance on ad-hoc feature engineering and threshold-sensitive binarization may conflate structural differences with feature scaling artifacts or obscure continuous edge weight information.

#### D.3.35. Maximum Common Subgraph Distance

The Maximum Common Subgraph (MCS) distance evaluates the structural similarity between the inferred and reference cell-type adjacency matrices, addressing whether the inferred trajectory graph captures the essential connectivity patterns of the ground-truth topology (Bunke & Shearer, 1998; Sanfeliu & Fu, 1983; Riesen & Bunke, 2009; Gao et al., 2010). The metric operates on the binary adjacency matrices (*A*_ref_, *A*_inf_), where the inferred matrix was binarized via a user-defined threshold. The core computation converts both matrices into undirected graphs and calculates their Graph Edit Distance (GED) under a cost model that allows free node substitutions (permitting graph isomorphism checks), prohibits node insertions/deletions via prohibitive costs, and assigns unit costs to edge insertions or deletions. This configuration reduces the GED to |ℰ_ref_| + |ℰ_inf_| −2 |ℰ_MCS_|, where ℰ_MCS_ denotes the edge set of the maximum common subgraph, thereby measuring the minimum number of edge operations needed to align the graphs. The MCS distance ranges from 0 (isomorphic graphs) upwards, with higher values indicating greater structural divergence; its computational complexity is NP-hard due to the combinatorial nature of subgraph isomorphism, though optimized heuristics enable tractable computation for graphs with *t* ≤ 60 cell types. A principal strength is its invariance to node permutations and satisfaction of metric properties (non-negativity, identity, symmetry, triangle inequality), ensuring robust topological comparison; however, reliance on exact GED computation limits scalability to large graphs, and potential non-convergence may yield NaN values, necessitating fallback strategies in practice.

#### D.3.36. Random Walk Kernel Distance

The Random Walk Kernel Distance quantifies the dissimilarity between the global connectivity structures of reference and inferred cell-type adjacency matrices, answering whether the inferred graph preserves the multiscale transition patterns encoded in the reference topology (Li et al., 2012; Kang et al., 2012; Kashima et al., 2003). The metric operates on symmetric adjacency matrices (*A*_ref_, *W*_inf_) ∈ {0, 1} ^*t*×*t*^ ×ℝ^*t*×*t*^, computing a normalized random walk kernel that compares walk counts across all path lengths up to a truncation limit *k*_max_. For decay factor *λ*, the kernel 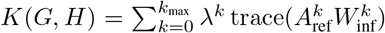 aggregates weighted traces of aligned matrix powers, capturing the agreement in walk structures; the normalized kernel 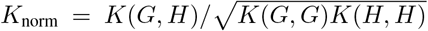 ensures scale invariance via Cauchy-Schwarz bounding (Borgwardt & Kriegel, 2005). The final distance 1−*K*_norm_ ranges in [0, 1], where 0 indicates identical walk structures and 1 maximal divergence; higher values denote increasing topological distortion. To ensure convergence, *λ* is constrained below 1*/*(*ρ*(*A*_ref_)*ρ*(*W*_inf_)), where *ρ* denotes spectral radius, with *k*_max_ automatically adjusted to avoid numerical insignificance. Computational complexity is *O*(*k*_max_*t*^3^) due to *k*_max_ matrix multiplications, limiting scalability for large *t*. The metric’s principal strength lies in its holistic integration of local and global connectivity via exponentially decaying walk contributions, providing robustness to localized discrepancies; however, sensitivity to *λ* and truncation effects may bias comparisons when long-range dependencies dominate biological transitions.

#### D.3.37. Persistence Diagram Distance

The Persistence Diagram Distance evaluates the topological fidelity of the inferred graph by quantifying discrepancies in multiscale topological features—specifically connected components (0-dimensional homology) and cycles (1-dimensional homology)—between the reference and inferred cell-type adjacency matrices (Cohen-Steiner et al., 2005; Kerber et al., 2017). The metric operates on (*A*_ref_, *W*_inf_), where *W*_inf_ is the inferred weighted adjacency matrix. For each matrix, edge weights were normalized to [0, 1], converted to distances via reciprocal transformation with regularization to avoid division by zero, and augmented by shortest-path distances to ensure global connectivity. A Vietoris-Rips filtration was applied to the resulting distance matrix (Bauer, 2021), constructing a simplicial complex that tracks the birth and death of topological features across scales; persistence diagrams for dimensions 0 (connected components) and 1 (cycles) were extracted, with infinite persistence features approximated by finite values scaled to the maximal finite death time. The 1-Wasserstein distance (Carriere et al., 2017) between corresponding diagrams of reference and inferred graphs was computed for each homology dimension and summed, yielding a composite topological divergence score. The score is non-negative, with lower values indicating closer topological correspondence; computational complexity is dominated by the *O*(*t*^3^) shortest-path calculation for *t* cell types and *O*(*t*^2^) persistence diagram comparisons. A principal strength lies in jointly capturing discrepancies in component structure and cyclic patterns, enabled by the stability of Wasserstein distance under perturbations; limitations include sensitivity to distance matrix regularization heuristics, exclusion of higher-dimensional topological features (e.g., voids), and prohibitive computational cost for large *t*.

#### D.3.38. Branch Silhouette Score

The Branch Silhouette Score evaluates whether cells from distinct trajectory branches form well-separated clusters in the inferred embedding, addressing the biological question of how effectively the representation preserves bifurcation events and lineage segregation (refer to D.2.1 for ASW description). The metric operates on the reference adjacency matrix *A*_ref_ and the full embedding *Y*, leveraging cell-type annotations to assign cells to terminal branches derived from *A*_ref_’s hierarchical structure. Each cell was mapped to the most terminal branch (leaf) reachable via the shortest path from the root in *A*_ref_, prioritizing biologically distal branches to maximize cluster distinctiveness. The silhouette score was computed as 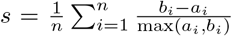, where *a*_*i*_ denotes the mean intra-branch distance and *b*_*i*_ the mean nearest-inter-branch distance for cell *i* in *Y*, using a user-specified distance metric. Scores range [−1, 1], with higher values indicating better branch separation; a score of ™1 implies perfect clustering, 1 complete overlap, and 0 indifferentiable branches. The computational complexity is *O*(*n*^2^) due to pairwise distance calculations, which may limit scalability for large *n*. A principal strength is the direct alignment with biological trajectory topology, ensuring clusters correspond to ground-truth branches; however, the metric assumes *A*_ref_ is a directed tree and requires sufficient cell-type annotations, failing to capture continuum-like differentiation or subtle substructure within branches.

#### D.3.39. Normalized Mean Curvature Score

The Normalized Mean Curvature Score (NMCS) quantifies the geometric complexity of developmental trajectories in the embedding space relative to the reference graph structure, answering whether inferred cell-state transitions exhibit biologically plausible curvature patterns. The metric operates on the reference adjacency matrix *A*_ref_ and cell embedding *Y*, where *A*_ref_ encodes permitted transitions between *t* cell types and *Y* provides lowdimensional coordinates for all cells. Developmental paths were extracted as source-to-sink routes in *A*_ref_, then represented in *Y* by cubic splines fitted to cell-type centroids along each path. For each spline *C*(*s*) parameterized by arc length *s*, the instantaneous curvature *κ* was computed via the generalized *n*-dimensional formula 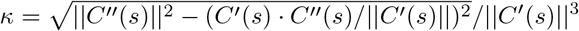, capturing local deviation from linear progression; path-level scores were normalized by *π* and aggregated across all paths using the median to yield the final score. NMCS ranges in [0, 1], where 0 indicates perfectly linear trajectories and 1 corresponds to maximal curvature complexity; higher values denote intricate branching patterns or abrupt fate decisions. The computational complexity is dominated by spline fitting (*O*(*n*) per path) and path enumeration (*O*(*t*^2^) for dense graphs). A principal strength lies in explicit modeling of branching topology through multiple developmental pathways, providing sensitivity to both global trajectory geometry and local transition smoothness; however, reliance on cubic splines may underestimate curvature in highly discontinuous embeddings, and path extraction becomes computationally intensive for graphs with exponential growth in source-sink routes.

#### D.3.40. Moran’s I for Embedding

Moran’s I evaluates whether the inferred embedding *Y* preserves the spatial autocorrelation structure dictated by the reference trajectory’s topology, addressing the biological question of how well local similarity patterns in the embedding align with the connectivity defined by *G*_ref_ (For a detailed derivation and the original citations, refer to Section D.3.16; the present formulation here is analogous apart from the alternative weighting scheme). The metric operates on (*A*_ref_, *Y*), where *A*_ref_ is the reference adjacency matrix and *Y* is the full cell embedding. To compute Moran’s I, the reference cell-type adjacency *A*_ref_ was first expanded into a cell-cell adjacency matrix *W* ∈ ℝ^*n*×*n*^ by connecting all cells of adjacent types, then row-normalized to form a spatial weights matrix. For each dimension of *Y*, Moran’s I was calculated as 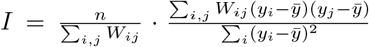 where *n* is the number of cells, *y*_*i*_ is the embedding value of cell *i*, and 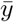 the dimension-wise mean; the final score averaged I across all embedding dimensions. Scores range approximately in [−1, 1], with 1 indicating perfect positive spatial autocorrelation (embedding similarity aligns with reference connectivity), -1 perfect dispersion, and 0 no spatial structure; higher values denote better preservation of the reference topology. The complexity is *O*(*n*) per embedding dimension due to sparse matrix-vector operations, making it linear in cell count for fixed dimensionality. A principal strength is its sensitivity to global and local embedding structure through the spatial weights matrix, providing a holistic measure of topological fidelity; however, the metric assumes cell-type adjacency perfectly reflects expected cellcell relationships, potentially overpenalizing embeddings that capture substructure within annotated types or exhibit biologically valid deviations from the reference graph.

#### D.3.41. Geary’s C for EMBEDDING

The Geary’s C Embedding metric quantifies the extent to which local dissimilarities in the full embedding *Y* align with non-adjacent cell-type relationships in the reference graph *G*_ref_, addressing whether biologically distant cell states exhibit systematically larger embedding-space deviations (For a detailed derivation and the original citations, refer to Section D.3.15; the present formulation here is analogous apart from the alternative weighting scheme). The metric operates on (*A*_ref_, *Y*), where *A*_ref_ is the reference adjacency matrix and *Y* the *N* -cell embedding. For each cell type in *G*_ref_, a cell-cell adjacency matrix was constructed by propagating *A*_ref_ to all cells via label-wise pooling, creating a sparse weight matrix *W* ∈ ℝ^*N* ×*N*^ where 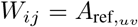 if cell *i* belongs to type *u* and cell *j* to type *v*. Geary’s C was computed per embedding dimension as 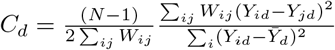 for dimension, then averaged across dimensions. Scores range [0, 2], with values *<* 1 indicating positive spatial autocorrelation (local homogeneity), 1 randomness, and *>* 1 anti-correlation (excessive local heterogeneity); lower scores denote better preservation of reference-driven dissimilarity patterns. Complexity is *O*(| ℰ|+ *Ne*^2^) for|ℰ |reference edges and *e* dimensions, dominated by pairwise difference computations. A principal strength is sensitivity to local variance anomalies complementary to global autocorrelation measures like Moran’s I; however, dependence on label-wise adjacency propagation may obscure cell-specific trajectory deviations, while normalization assumptions limit interpretability for embeddings with non-stationary variance structures.

#### D.3.42. Embedding Distance Correlation

This metric evaluates whether the relative distances between cell types in a low-dimensional embedding space mirror their topological relationships in the reference trajectory graph, addressing how well the embedding preserves the, graph’s structural geometry (Székely et al., 2007; Mantel, 1967; Moon et al., 2019). The computation operates on the reference graph adjacency matrix *A*_ref_ and the full embedding matrix *Y*, leveraging cell-type labels to compute centroid positions in the embedding space. For each pair of cell types, the shortest path distance in the undirected version of *A*_ref_ was computed via breadth-first search, while their embedding distance was derived as the Euclidean distance between corresponding centroids. Spearman’s rank correlation coefficient *ρ* was then calculated between these two sets of pairwise distances, excluding pairs disconnected in the reference graph. The resultant *ρ* ranges in [−1, 1], where 1 indicates perfect monotonic correspondence between graph and embedding distances, 0 implies no association, and -1 denotes perfect inverse correlation (biologically implausible); higher values signify stronger structural preservation. With *t* cell types, complexity is *O*(*t*^2^) from pairwise comparisons and all-pairs shortest paths, remaining tractable for typical *t* values. A principal strength is the metric’s invariance to scaling transformations in either space, focusing purely on relative distance ordering; however, reliance on distinct cell-type centroids assumes homogeneous populations and may fail when within-type variance dominates between-type separation, while quadratic scaling limits utility for extremely large type sets.

#### D.3.43. Sammon’s Stress

Sammon’s Stress evaluates how well the low-dimensional embedding *Y* preserves the global and local topological structure of the reference trajectory graph *G* _ref_, with emphasis on accurate representation of proximate cell-type relationships critical for biological transitions (Sammon, 2006; Lee & Verleysen, 2008; Ghojogh et al., 2020). The metric operates on the reference adjacency matrix *A*_ref_ and the full embedding *Y*, computing pairwise shortest-path distances between cell types in *G*_ref_ and Euclidean or cosine distances between their centroids in *Y*. To ensure scale invariance, an optimal scaling factor *s*^∗^ was analytically derived to minimize the stress function 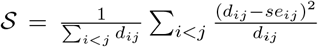 where *d*_*ij*_ and *e*_*ij*_ denote graph and embedding distances, respectively; *s*^∗^ is given by 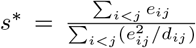. The resulting stress *S* is non-negative, with lower values (theoretically bounded below by 0) indicating stronger structural preservation; values below 0.1 typically denote high fidelity. Computational complexity is dominated by all-pairs shortest paths (*O*(*t*^3^) for *t* cell types) and centroid distance calculations (*O*(*t*^2^*e*)), though sparse graphs reduce this to *O*(*t*^2^) with connected components. A principal strength is the optimal scaling factor, which eliminates sensitivity to arbitrary embedding scales while emphasizing local relationships through inverse-distance weighting; however, reliance on cell-type centroids may obscure within-type heterogeneity, and exclusion of disconnected node pairs limits applicability to fully connected reference trajectories.

#### D.3.44. Graph-Based Trustworthiness

The Graph-Based Trustworthiness metric evaluates the preservation of local neighborhoods between the cell type hierarchy defined by the reference graph and the lowdimensional embedding, addressing whether biologically proximate cell types in *G*_ref_ remain neighboring in the learned manifold (Venna & Kaski, 2001; Jiang et al., 2016; Ge et al., 2023). The score operates on the reference adjacency matrix *A*_ref_ and the full embedding *Y*, computing for each cell the permissible cross-type neighbors based on graph distances and penalizing those that appear prematurely in the embedding’s nearest neighbors. For a cell *i* of type *v*, the allowed rank threshold for cross-type neighbors was determined by *t*_*i*_ = max(*k* ™*m*_*i*_, 0), where *m*_*i*_ counts same-type neighbors; penalties accrued when cross-type neighbors of *i* in *Y* exceeded *t*_*i*_ in graph-based rank, computed via shortest-path distances and cumulative type counts. The total penalty was normalized by the theoretical maximum—accounting for cell type frequencies and graph connectivity—yielding a score *T* (*k*) = 1 ™ (∑ penalties)*/*(∑ max penalties). Scores range in [0, 1], with 1 indicating perfect neighborhood preservation and 0 maximal violation; higher values denote better alignment. Preprocessing graph distances requires *O*(*t*^2^) time, while neighbor validation scales as *O*(*nk*) for *n* cells and *k* neighbors. The metric’s principal strength lies in leveraging explicit cell type hierarchies to assess embedding fidelity, ensuring robustness to varying type distributions; however, reliance on shortest-path distances may overlook nuanced biological relationships, and preprocessing becomes prohibitive for large *t*.

#### D.3.45. Neighborhood Preservation Score

The Neighborhood Preservation Score (NPS) evaluates how well an embedding preserves transitions between cell types defined by the reference biological graph, addressing whether local neighborhoods in the embedding space correspond to permissible state transitions (Lee & Verleysen, 2008; Heiser & Lau, 2020; Martins et al., 2015). The metric operates on the reference adjacency matrix *A*_ref_ and the inferred embedding *Y*, leveraging the K-nearest neighbor (KNN) graph derived from *Y* to assess neighborhood composition. For each cell, neighbors in the KNN graph were examined to identify those with cell types adjacent to the cell’s type in *A*_ref_; cells with a proportion of different-type neighbors exceeding a threshold *τ* were retained, and the fraction of their different-type neighbors corresponding to allowed transitions in *A*_ref_ was computed. The final score was obtained by averaging these fractions across qualifying cells. NPS ranges from 0 to 1, where 1 indicates perfect preservation of reference transitions in embedding neighborhoods, and 0 denotes no alignment; higher values reflect better biological fidelity. The computation scales linearly with the number of cells *n, O*(*n*), assuming a fixed KNN size. A principal strength is its focus on biologically critical cross-type transitions, enhancing robustness to homogeneous neighborhoods through thresholding; however, the score depends on the choice of *τ* and may underrepresent transitions in cells with sparse cross-type interactions.

#### D.3.46. Directionality Preservation

The Directionality Preservation metric evaluates whether local directional relationships in the inferred embedding *Y* align with transitions encoded in the reference graph *G*_ref_, addressing whether biological trajectories manifest as geometrically coherent progressions in the embedding space. The metric operates on (*A*_ref_, *Y*), where *A*_ref_ is the reference adjacency matrix and *Y* the full embedding. For each edge (*u* → *v*) in *G*_ref_, the centroids of nodes *u* and *v* were computed, defining a graph direction vector Δ_*uv*_ = centroid_*v*_ ™ centroid_*u*_. Cells proximal to the line segment between centroids were identified, expanded via their k-nearest neighbors, and subjected to local PCA; the cosine similarity between the first principal component (PC1) and Δ_*uv*_ was calculated, measuring directional alignment. The final score averaged these similarities across all edges. Values range in [−1, 1], with 1 indicating perfect alignment, 0 no systematic correlation, and -1 anti-correlation; higher scores reflect stronger preservation of reference directions. Complexity scales as *O*(|ℰ | (*ke* + *e*^3^)) for|ℰ|edges, *k* cells per edge, and *e* embedding dimensions. A principal strength is the integration of local manifold structure through PCA, enabling detection of directional signals beyond pairwise distances; however, reliance on linear PCA components and sensitivity to neighbor-graph parameters may obscure nonlinear directional relationships or overemphasize spurious local correlations.

#### D.3.47. Wasserstein Distance for Embedding

The Wasserstein Distance (WD) evaluates the alignment between the distribution of cell-type transitions prescribed by the biological graph and the empirical transition patterns observed in the embedding’s neighborhood structure. This metric addresses whether the embedding preserves the permissible state transitions defined by the reference trajectory while respecting their topological proximity (Huizing et al., 2022; Schiebinger et al., 2019; Peyré et al., 2019; Panaretos & Zemel, 2019; Piccoli & Rossi, 2016; Carriere et al., 2017). The computation operates on the reference adjacency matrix *A*_ref_ and the embedding *Y*, leveraging the latter’s k-nearest-neighbor (kNN) graph to model local transition likelihoods. For each cell type *u*, two probability distributions were constructed: *p*_*u*_ (reference transitions), derived from *A*_ref_’s outgoing edges from *u*, and *q*_*u*_ (embedding transitions), estimated from the kNN cell-type composition among neighbors of cells annotated as *u*. The Wasserstein distance between *p*_*u*_ and *q*_*u*_ was computed using a cost matrix *C* ∈ ℝ ^*t*×*t*^, where *C*_*ij*_ encoded the shortest-path distance between cell types *i* and *j* in *A*_ref_; unreachable pairs incurred a penalty of *t* + 1. The final score averaged these distances across all cell types. WD ranges from 0 (perfect alignment) to unbounded positive values, with lower scores indicating better structural concordance. Computational complexity is *O*(*n* + *t*^4^), dominated by the *O*(*t*^3^) earth mover’s distance computation per cell type and *O*(*n*) neighbor aggregation. A key strength is the integration of global graph topology via shortest-path costs, ensuring biologically meaningful penalization of transitions violating trajectory connectivity. However, WD assumes kNN neighborhoods directly reflect transition probabilities and may over-penalize disconnected graph components, potentially conflating biological rarity with structural incompatibility.

#### D.3.48. Directed Trajectory Validation Score

The Directed Trajectory Validation Score (DTVS) evaluates whether the local neighborhood structure in a cell embedding *Y* aligns with the branching topology of a reference trajectory *A*_ref_, addressing how well the embedding preserves biologically defined developmental bifurcations (Jeon et al., 2023; Thrun et al., 2023; Liu et al., 2019; Rieck & Leitte, 2015; Cape et al., 2024). The metric operates on (*A*_ref_, *Y*) by first identifying branch points in *A*_ref_ and their descendant branches, then comparing spectral clusters derived from *Y* ‘s k-nearest-neighbor graph against branch memberships predicted through neighbor voting. For each branch point, cells were assigned to branches via majority voting of their neighbors’ cell-type labels in *Y*, while spectral clustering with *Y* ‘s connectivities matrix generated topology-driven clusters; the weighted F1-score between these assignments quantified local-to-global consistency, with harmonic penalties applied for undetected branches. Scores range in [0, 1], where 1 indicates perfect alignment between embedding neighborhoods and reference branches, and 0 denotes complete discordance; computational complexity is dominated by spectral clustering at *O*(*n*^3^) per branch point. The principal strength lies in jointly evaluating local neighbor agreements and global manifold structure through multi-scale validation; however, dependence on accurate cell-type annotations and sensitivity to the spectral clustering hyperparameters may obscure topological fidelity when label noise exists or branch sizes are imbalanced.

## Footnotes

1 *’Not all those who wander are lost*.*’ — J. R. R. Tolkien* Even embeddings that appear visually disordered may still honor the intrinsic trajectory manifold, highlighting the need for quantitative metrics to tell the difference.

2 Refer to Supplementary D.1 for comprehensive description of the employed decomposition approach.

3 Refer to Figure S7 for complete graph annotations and Figure S8 for UMAP projections of the full hematopoietic manifold.

4 For a detailed discussion of the theoretical assumptions and known limitations, refer to Supplementary B. Experimental insights—including dataset descriptions, model trajectory details, and a link to the *scTRAM* **codebase**—are provided in Supplementary C.

5 A post-hoc audit revealed that the early-myeloid lineage in the *Suo* dataset was misannotated; as none of the analyses or inferences reported here depend on that segment, we have left the original labels intact to preserve exact reproducibility and will issue corrected annotations in a subsequent revision.

## References

Abdi, H. and Williams, L. J. Principal component analysis. Wiley interdisciplinary reviews: computational statistics, 2(4):433–459, 2010.

Abu-Aisheh, Z., Raveaux, R., Ramel, J.-Y., and Martineau, P. An exact graph edit distance algorithm for solving pattern recognition problems. In 4th International Conference on Pattern Recognition Applications and Methods 2015, 2015.

Aliee, H., Kapl, F., Pham, D., Cakir, B., Jimba, T., Cranley, J., Teichmann, S. A., Meyer, K. B., Vento-Tormo, R., and Theis, F. J. invae: Conditionally invariant representation learning for generating multivariate single-cell reference maps. bioRxiv, pp. 2024–12, 2024.

Andersen, R., Chung, F., and Lang, K. Using pagerank to locally partition a graph. Internet Mathematics, 4(1): 35–64, 2007.

Anselin, L. Local indicators of spatial association—lisa. Geographical analysis, 27(2):93–115, 1995.

Aouchiche, M. and Hansen, P. Two laplacians for the distance matrix of a graph. Linear algebra and its applications, 439(1):21–33, 2013.

Bai, Y., Ding, H., Bian, S., Chen, T., Sun, Y., and Wang, W. Simgnn: A neural network approach to fast graph similarity computation. In Proceedings of the twelfth ACM international conference on web search and data mining, pp. 384–392, 2019.

Bakurov, I., Buzzelli, M., Schettini, R., Castelli, M., and Vanneschi, L. Structural similarity index (ssim) revisited: A data-driven approach. Expert Systems with Applications, 189:116087, 2022.

Bauer, U. Ripser: efficient computation of vietoris–rips persistence barcodes. Journal of Applied and Computational Topology, 5(3):391–423, 2021.

Belghazi, M. I., Baratin, A., Rajeshwar, S., Ozair, S., Bengio, Y., Courville, A., and Hjelm, D. Mutual information neural estimation. In International conference on machine learning, pp. 531–540. PMLR, 2018.

Bergen, V., Lange, M., Peidli, S., Wolf, F. A., and Theis, F. J. Generalizing rna velocity to transient cell states through dynamical modeling. Nature biotechnology, 38 (12):1408–1414, 2020.

Berndt, D. J. and Clifford, J. Using dynamic time warping to find patterns in time series. In Proceedings of the 3rd international conference on knowledge discovery and data mining, pp. 359–370, 1994.

Besta, M., Kanakagiri, R., Mustafa, H., Karasikov, M., Rätsch, G., Hoefler, T., and Solomonik, E. Communication-efficient jaccard similarity for high-performance distributed genome comparisons. In 2020 IEEE International Parallel and Distributed Processing Symposium (IPDPS), pp. 1122–1132. IEEE, 2020.

Biology, C. S.-C., Abdulla, S., Aevermann, B., Assis, P., Badajoz, S., Bell, S. M., Bezzi, E., Cakir, B., Chaffer, J., Chambers, S., et al. Cz cellxgene discover: A single-cell data platform for scalable exploration, analysis and modeling of aggregated data. bioRxiv, pp. 2023–10, 2023.

Blumenthal, D. B., Boria, N., Gamper, J., Bougleux, S., and Brun, L. Comparing heuristics for graph edit distance computation. The VLDB journal, 29(1):419–458, 2020.

Borcard, D. and Legendre, P. Is the mantel correlogram powerful enough to be useful in ecological analysis? a simulation study. Ecology, 93(6):1473–1481, 2012.

Borgwardt, K. M. and Kriegel, H.-P. Shortest-path kernels on graphs. In Fifth IEEE international conference on data mining (ICDM’05), pp. 8–pp. IEEE, 2005.

Böttcher, A. and Wenzel, D. The frobenius norm and the commutator. Linear algebra and its applications, 429 (8-9):1864–1885, 2008.

Brunet, D., Vrscay, E. R., and Wang, Z. On the mathematical properties of the structural similarity index. IEEE Transactions on Image Processing, 21(4):1488–1499, 2011.

Bunke, H. and Shearer, K. A graph distance metric based on the maximal common subgraph. Pattern recognition letters, 19(3-4):255–259, 1998.

Büttner, M., Miao, Z., Wolf, F. A., Teichmann, S. A., and Theis, F. J. A test metric for assessing single-cell rna-seq batch correction. Nature Methods, 16(1):43–49, December 2018. ISSN 1548-7105.

Cape, J., Yu, X., and Liao, J. Z. Robust spectral clustering with rank statistics. Journal of Machine Learning Research, 25(398):1–81, 2024.

Carriere, M., Cuturi, M., and Oudot, S. Sliced wasserstein kernel for persistence diagrams. In International conference on machine learning, pp. 664–673. PMLR, 2017.

Chari, T. and Pachter, L. The specious art of single-cell genomics. PLOS Computational Biology, 19(8):e1011288, 2023.

Chen, C., Peng, R., Ying, L., and Tong, H. Network connectivity optimization: Fundamental limits and effective algorithms. In Proceedings of the 24th ACM SIGKDD International Conference on Knowledge Discovery & Data Mining, pp. 1167–1176, 2018.

Chen, Z., Villar, S., Chen, L., and Bruna, J. On the equivalence between graph isomorphism testing and function approximation with gnns. Advances in neural information processing systems, 32, 2019.

Cohen-Steiner, D., Edelsbrunner, H., and Harer, J. Stability of persistence diagrams. In Proceedings of the twentyfirst annual symposium on Computational geometry, pp. 263–271, 2005.

Cui, H., Tejada-Lapuerta, A., Brbić, M., Saez-Rodriguez, J., Cristea, S., Goodarzi, H., Lotfollahi, M., Theis, F. J., and Wang, B. Towards multimodal foundation models in molecular cell biology. Nature, 640(8059):623–633, 2025.

de Jong, P., Sprenger, C., and van Veen, F. On extreme values of moran’s i and geary’s c. Geographical Analysis, 16(1):17–24, 1984.

DeTomaso, D. and Yosef, N. Hotspot identifies informative gene modules across modalities of single-cell genomics. Cell systems, 12(5):446–456, 2021.

DeTomaso, D., Jones, M. G., Subramaniam, M., Ashuach, T., Ye, C. J., and Yosef, N. Functional interpretation of single cell similarity maps. Nature communications, 10 (1):4376, 2019.

Dodonova, Y., Belyaev, M., Tkachev, A., Petrov, D., and Zhukov, L. Kernel classification of connectomes based on earth mover’s distance between graph spectra. arXiv preprint 1611.08812, 2016.

Drezner, Z., Turel, O., and Zerom, D. A modified kolmogorov–smirnov test for normality. Communications in Statistics—Simulation and Computation®, 39(4): 693–704, 2010.

Duncan, T. E. On the calculation of mutual information. SIAM Journal on Applied Mathematics, 19(1):215–220, 1970.

Ehounou, W., Barth, D., De Moissac, A., Watel, D., and Weisser, M.-A. Minimizing the hamming distance between a graph and a line-graph to discover the topology of an electrical network. Journal of Graph Algorithms and Applications, 24(3):133–153, 2020.

Essam, F., El, H., and Ali, S. R. H. A comparison of the pearson, spearman rank and kendall tau correlation coefficients using quantitative variables. Asian J. Probab. Stat, pp. 36–48, 2022.

Freedman, D. A. and Diaconis, P. On the histogram as a density estimator: L 2 theory. Zeitschrift für Wahrscheinlichkeitstheorie und Verwandte Gebiete, 57(4):453–476, 1981.

Gambhir, R., Larkoski, A. J., and Thaler, J. Specter: efficient evaluation of the spectral emd. Journal of High Energy Physics, 2024(12):1–48, 2025.

Ganagi, A. B. and Ramane, H. S. Hamming distance between the strings generated by adjacency matrix of a graph and their sum. Algebra and discrete mathematics, 22(1), 2016.

Gao, X., Xiao, B., Tao, D., and Li, X. A survey of graph edit distance. Pattern Analysis and applications, 13:113–129, 2010.

Garcia-Alonso, L., Lorenzi, V., Mazzeo, C. I., Alves-Lopes, J. P., Roberts, K., Sancho-Serra, C., Engelbert, J., Marečková, M., Gruhn, W. H., Botting, R. A., et al. Single-cell roadmap of human gonadal development. Nature, 607(7919):540–547, 2022.

Gayoso, A. scib-metrics: Accelerated, Pythononly, single-cell integration benchmarking metrics, 2025. URL https://github.com/YosefLab/scib-metrics. xAccessed 2025-05-19.

Ge, Y., Ma, J., Zhang, L., Li, X., and Lu, H. Trustworthinessaware knowledge graph representation for recommendation. Knowledge-Based Systems, 278:110865, 2023.

Genest, C., Quessy, J.-F., and Rémillard, B. Local efficiency of a cramér–von mises test of independence. Journal of Multivariate Analysis, 97(1):274–294, 2006.

Genest, C., Quessy, J.-F., and Rémillard, B. Asymptotic local efficiency of cramér–von mises tests for multivariate independence. arXiv preprint 2503.04393, 2007.

Ghojogh, B., Ghodsi, A., Karray, F., and Crowley, M. Multidimensional scaling, sammon mapping, and isomap: Tutorial and survey. arXiv preprint 2009.08136, 2020.

Goldberg, A. V. and Harrelson, C. Computing the shortest path: A search meets graph theory. In SODA, volume 5, pp. 156–165, 2005.

Gonzalez-Escribano, A., Llanos, D. R., and Ortega-Arranz, H. The shortest-path problem: Analysis and comparison of methods. Springer Nature, 2022.

Gu, A. and Dao, T. Mamba: Linear-time sequence modeling with selective state spaces. arXiv preprint 2312.00752, 2023.

Haghverdi, L., Buettner, F., and Theis, F. J. Diffusion maps for high-dimensional single-cell analysis of differentiation data. Bioinformatics, 31(18):2989–2998, 2015.

Haghverdi, L., Lun, A. T. L., Morgan, M. D., and Marioni, J. C. Batch effects in single-cell rna-sequencing data are corrected by matching mutual nearest neighbors. Nature Biotechnology, 36(5):421–427, April 2018. ISSN 1546-1696. doi: 10.1038/nbt.4091.

Hastie, T. J. Generalized additive models. Statistical models in S, pp. 249–307, 2017.

Heiser, C. N. and Lau, K. S. A quantitative framework for evaluating single-cell data structure preservation by dimensionality reduction techniques. Cell reports, 31(5), 2020.

Heumos, L., Schaar, A. C., Lance, C., Litinetskaya, A., Drost, F., Zappia, L., Lücken, M. D., Strobl, D. C., Henao, J., Curion, F., et al. Best practices for single-cell analysis across modalities. Nature Reviews Genetics, 24(8):550– 572, 2023.

Hodson, T. O. Root mean square error (rmse) or mean absolute error (mae): When to use them or not. Geoscientific Model Development Discussions, 2022:1–10, 2022.

Hougardy, S. The floyd–warshall algorithm on graphs with negative cycles. Information Processing Letters, 110 (8-9):279–281, 2010.

Hu, Y., Wan, S., Luo, Y., Li, Y., Wu, T., Deng, W., Jiang, C., Jiang, S., Zhang, Y., Liu, N., et al. Benchmarking algorithms for single-cell multi-omics prediction and integration. Nature Methods, 21(11):2182–2194, 2024.

Hubert, L. and Arabie, P. Comparing partitions. Journal of Classification, 2(1):193–218, December 1985. ISSN 1432-1343.

Huffaker, B., Dhamdhere, A., Fomenkov, M., and Claffy, K. Toward topology dualism: improving the accuracy of as annotations for routers. In International Conference on Passive and Active Network Measurement, pp. 101–110. Springer, 2010.

Huizing, G.-J., Peyré, G., and Cantini, L. Optimal transport improves cell–cell similarity inference in single-cell omics data. Bioinformatics, 38(8):2169–2177, 2022.

Inecik, K. and Theis, F. scare: Attribution regularization for single cell representation learning. bioRxiv, pp. 2023–07, 2023.

Inecik, K., Kara, A., Rose, A., Haniffa, M., and Theis, F. J. Tardis: Achieving robust and structured disentanglement of multiple covariates. bioRxiv, 2024.

James, G., Witten, D., Hastie, T., Tibshirani, R., et al. An introduction to statistical learning, volume 112. Springer, 2013.

Jeon, H., Kuo, Y.-H., Aupetit, M., Ma, K.-L., and Seo, J. Classes are not clusters: Improving label-based evaluation of dimensionality reduction. IEEE Transactions on Visualization and Computer Graphics, 30(1):781–791, 2023.

Jiang, W., Wang, G., Bhuiyan, M. Z. A., and Wu, J. Understanding graph-based trust evaluation in online social networks: Methodologies and challenges. Acm Computing Surveys (Csur), 49(1):1–35, 2016.

Jovanović, I. and Stanić, Z. Spectral distances of graphs. Linear algebra and its applications, 436(5):1425–1435, 2012.

Kang, U., Tong, H., and Sun, J. Fast random walk graph kernel. In Proceedings of the 2012 SIAM international conference on data mining, pp. 828–838. SIAM, 2012.

Kashima, H., Tsuda, K., and Inokuchi, A. Marginalized kernels between labeled graphs. In Proceedings of the 20th international conference on machine learning (ICML-03), pp. 321–328, 2003.

Kerber, M., Morozov, D., and Nigmetov, A. Geometry helps to compare persistence diagrams, 2017.

Kim, B.-H. and Ye, J. C. Understanding graph isomorphism network for rs-fmri functional connectivity analysis. Frontiers in neuroscience, 14:630, 2020.

Klein, D., Fleck, J. S., Bobrovskiy, D., Zimmermann, L., Becker, S., Palma, A., Dony, L., Tejada-Lapuerta, A., Huguet, G., Lin, H.-C., et al. Cellflow enables generative single-cell phenotype modeling with flow matching. bioRxiv, pp. 2025–04, 2025.

Konukoglu, E., Glocker, B., Criminisi, A., and Pohl, K. M. Wesd–weighted spectral distance for measuring shape dissimilarity. IEEE transactions on pattern analysis and machine intelligence, 35(9):2284–2297, 2012.

Korsunsky, I., Millard, N., Fan, J., Slowikowski, K., Zhang, F., Wei, K., Baglaenko, Y., Brenner, M., Loh, P.-r., and Raychaudhuri, S. Fast, sensitive and accurate integration of single-cell data with harmony. Nature Methods, 16 (12):1289–1296, November 2019a. ISSN 1548-7105.

Korsunsky, I., Millard, N., Fan, J., Slowikowski, K., Zhang, F., Wei, K., Baglaenko, Y., Brenner, M., Loh, P.-r., and Raychaudhuri, S. Fast, sensitive and accurate integration of single-cell data with harmony. Nature methods, 16(12): 1289–1296, 2019b.

Kraskov, A., Stögbauer, H., and Grassberger, P. Estimating mutual information. Physical review E, 69(6):066138, 2004.

Kriege, N. M., Johansson, F. D., and Morris, C. A survey on graph kernels. Applied Network Science, 5:1–42, 2020.

Kvålseth, T. O. On normalized mutual information: measure derivations and properties. Entropy, 19(11):631, 2017.

La Manno, G., Soldatov, R., Zeisel, A., Braun, E., Hochgerner, H., Petukhov, V., Lidschreiber, K., Kastriti, M. E., Lönnerberg, P., Furlan, A., et al. Rna velocity of single cells. Nature, 560(7719):494–498, 2018.

Lee, J. and Verleysen, M. Quality assessment of nonlinear dimensionality reduction based on k-ary neighborhoods. New Challenges for Feature Selection in Data Mining and Knowledge Discovery, pp. 21–35, 2008.

Legendre, P. and Fortin, M.-J. Comparison of the mantel test and alternative approaches for detecting complex multivariate relationships in the spatial analysis of genetic data. Molecular ecology resources, 10(5):831–844, 2010.

Legendre, P., Fortin, M.-J., and Borcard, D. Should the mantel test be used in spatial analysis? Methods in Ecology and Evolution, 6(11):1239–1247, 2015.

Li, X., Su, X., and Wang, M. Social network-based recommendation: a graph random walk kernel approach. In Proceedings of the 12th ACM/IEEE-CS joint conference on Digital Libraries, pp. 409–410, 2012.

Liao, H. and He, J. Jacobian determinant of normalizing flows. arXiv preprint 2102.06539, 2021.

Liu, X., Zhuang, C., Murata, T., Kim, K.-S., and Kertkeidkachorn, N. How much topological structure is preserved by graph embeddings? Computer Science and Information Systems, 16:11–11, 2019.

Longato, E., Vettoretti, M., and Di Camillo, B. A practical perspective on the concordance index for the evaluation and selection of prognostic time-to-event models. Journal of biomedical informatics, 108:103496, 2020.

Lopes, R. H., Reid, I., and Hobson, P. R. The twodimensional kolmogorov-smirnov test. arXiv preprint 2503.04393, 2007.

Lopez, R., Regier, J., Cole, M. B., Jordan, M. I., and Yosef, N. Deep generative modeling for single-cell transcriptomics. Nature methods, 15(12):1053–1058, 2018.

Lotfollahi, M., Klimovskaia Susmelj, A., De Donno, C., Hetzel, L., Ji, Y., Ibarra, I. L., Srivatsan, S. R., Naghipourfar, M., Daza, R. M., Martin, B., et al. Predicting cellular responses to complex perturbations in high-throughput screens. Molecular systems biology, 19(6):e11517, 2023.

Luecken, M. D., Büttner, M., Chaichoompu, K., Danese, A., Interlandi, M., Müller, M. F., Strobl, D. C., Zappia, L., Dugas, M., Colomé-Tatché, M., et al. Benchmarking atlas-level data integration in single-cell genomics. Nature methods, 19(1):41–50, 2022.

Mantel, N. The detection of disease clustering and a generalized regression approach. Cancer research, 27(2 Part 1): 209–220, 1967.

Mao, G. and Zhang, N. Analysis of average shortest-path length of scale-free network. Journal of Applied Mathematics, 2013(1):865643, 2013.

Martins, R. M., Minghim, R., Telea, A. C., et al. Explaining neighborhood preservation for multidimensional projections. In CGVC, pp. 7–14, 2015.

Milenković, T. and Pržulj, N. Uncovering biological network function via graphlet degree signatures. Cancer informatics, 6:CIN–S680, 2008.

Miller, B., Bliss, N., and Wolfe, P. Subgraph detection using eigenvector l1 norms. Advances in Neural Information Processing Systems, 23, 2010.

Moon, K. R., Van Dijk, D., Wang, Z., Gigante, S., Burkhardt, D. B., Chen, W. S., Yim, K., Elzen, A. v. d., Hirn, M. J., Coifman, R. R., et al. Visualizing structure and transitions in high-dimensional biological data. Nature biotechnology, 37(12):1482–1492, 2019.

Morris, C., Ritzert, M., Fey, M., Hamilton, W. L., Lenssen, J. E., Rattan, G., and Grohe, M. Weisfeiler and leman go neural: Higher-order graph neural networks. In Proceedings of the AAAI conference on artificial intelligence, volume 33, pp. 4602–4609, 2019.

Munkres, J. R. Topology. Prentice Hall, Upper Saddle River, NJ, 2nd edition, 2000. ISBN 9780131816299.

Najim, A. A. More faithfulness graph embedding. International Journal of Applied Mathematics Research, 4(2): 336, 2015.

Nath, M. and Paul, S. On the distance laplacian spectra of graphs. Linear Algebra and its Applications, 460:97–110, 2014.

Opsahl, T. and Panzarasa, P. Clustering in weighted networks. Social networks, 31(2):155–163, 2009.

Panaretos, V. M. and Zemel, Y. Statistical aspects of wasserstein distances. Annual review of statistics and its application, 6(1):405–431, 2019.

Peng, X., Lu, C., Yi, Z., and Tang, H. Connections between nuclear-norm and frobenius-norm-based representations. IEEE transactions on neural networks and learning systems, 29(1):218–224, 2016.

Peyré, G., Cuturi, M., et al. Computational optimal transport: With applications to data science. Foundations and Trends® in Machine Learning, 11(5-6):355–607, 2019.

Piccoli, B. and Rossi, F. On properties of the generalized wasserstein distance. Archive for Rational Mechanics and Analysis, 222:1339–1365, 2016.

Powers, D. M. Evaluation: from precision, recall and fmeasure to roc, informedness, markedness and correlation. arXiv preprint 2010.16061, 2020.

Pržulj, N. Biological network comparison using graphlet degree distribution. Bioinformatics, 23(2):e177–e183, 2007.

Qi, J., Du, J., Siniscalchi, S. M., Ma, X., and Lee, C.-H. On mean absolute error for deep neural network based vectorto-vector regression. IEEE Signal Processing Letters, 27: 1485–1489, 2020.

Qu, R., Cheng, X., Sefik, E., Stanley III, J. S., Landa, B., Strino, F., Platt, S., Garritano, J., Odell, I. D., Coifman, R., et al. Gene trajectory inference for single-cell data by optimal transport metrics. Nature biotechnology, 43(2): 258–268, 2025.

Qu, T. and Cai, Z. An improved isomap method for manifold learning. International Journal of Intelligent Computing and Cybernetics, 10(1):30–40, 2017.

Ratner, B. Statistical and machine-learning data mining:: Techniques for better predictive modeling and analysis of big data. Chapman and Hall/CRC, 2017.

Razali, N. M., Wah, Y. B., et al. Power comparisons of shapiro-wilk, kolmogorov-smirnov, lilliefors and anderson-darling tests. Journal of statistical modeling and analytics, 2(1):21–33, 2011.

Rieck, B. and Leitte, H. Persistent homology for the evaluation of dimensionality reduction schemes. Computer Graphics Forum, 34:431–440, 2015.

Riesen, K. and Bunke, H. Approximate graph edit distance computation by means of bipartite graph matching. Image and Vision computing, 27(7):950–959, 2009.

Rousseeuw, P. Rousseeuw, p.j.: Silhouettes: A graphical aid to the interpretation and validation of cluster analysis. comput. appl. math. 20, 53-65. Journal of Computational and Applied Mathematics, 20:53–65, 11 1987a. doi: 10.1016/0377-0427(87)90125-7.

Rousseeuw, P. J. Silhouettes: A graphical aid to the interpretation and validation of cluster analysis. Journal of Computational and Applied Mathematics, 20:53–65, November 1987b. ISSN 0377-0427.

Saelens, W., Cannoodt, R., Todorov, H., and Saeys, Y. A comparison of single-cell trajectory inference methods. Nature biotechnology, 37(5):547–554, 2019.

Sammon, J. W. A nonlinear mapping for data structure analysis. IEEE Transactions on computers, 100(5):401– 409, 2006.

Sanfeliu, A. and Fu, K.-S. A distance measure between attributed relational graphs for pattern recognition. IEEE transactions on systems, man, and cybernetics, pp. 353– 362, 1983.

Sathre, P., Gondhalekar, A., and Feng, W.-c. Edge-connected jaccard similarity for graph link prediction on fpga. In 2022 IEEE High Performance Extreme Computing Conference (HPEC), pp. 1–10. IEEE, 2022.

Schaar, A. C., Tejada-Lapuerta, A., Palla, G., Gutgesell, R., Halle, L., Minaeva, M., Vornholz, L., Dony, L., Drummer, F., Bahrami, M., et al. Nicheformer: a foundation model for single-cell and spatial omics. bioRxiv, pp. 2024–04, 2024.

Schiebinger, G., Shu, J., Tabaka, M., Cleary, B., Subramanian, V., Solomon, A., Gould, J., Liu, S., Lin, S., Berube, P., et al. Optimal-transport analysis of single-cell gene expression identifies developmental trajectories in reprogramming. Cell, 176(4):928–943, 2019.

Scott, D. W. On optimal and data-based histograms. Biometrika, 66(3):605–610, 1979.

Shervashidze, N., Schweitzer, P., Van Leeuwen, E. J., Mehlhorn, K., and Borgwardt, K. M. Weisfeiler-lehman graph kernels. Journal of Machine Learning Research, 12(9), 2011.

Sokolova, M. and Lapalme, G. A systematic analysis of performance measures for classification tasks. Information Processing; Management, 45(4):427–437, July 2009. ISSN 0306-4573. doi: 10.1016/j.ipm.2009.03.002.

Song, Y., Miao, Z., Brazma, A., and Papatheodorou, I. Benchmarking strategies for cross-species integration of single-cell rna sequencing data. Nature Communications, 14(1):6495, 2023.

Srivatsan, S. R., McFaline-Figueroa, J. L., Ramani, V., Saunders, L., Cao, J., Packer, J., Pliner, H. A., Jackson, D. L., Daza, R. M., Christiansen, L., et al. Massively multiplex chemical transcriptomics at single-cell resolution. Science, 367(6473):45–51, 2020.

Steck, H., Krishnapuram, B., Dehing-Oberije, C., Lambin, P., and Raykar, V. C. On ranking in survival analysis: Bounds on the concordance index. Advances in neural information processing systems, 20, 2007.

Steinskog, D. J., Tjøstheim, D. B., and Kvamstø, N. G. A cautionary note on the use of the kolmogorov–smirnov test for normality. Monthly Weather Review, 135(3):1151– 1157, 2007.

Suo, C., Dann, E., Goh, I., Jardine, L., Kleshchevnikov, V., Park, J.-E., Botting, R. A., Stephenson, E., Engelbert, J., Tuong, Z. K., et al. Mapping the developing human immune system across organs. Science, 376(6597): eabo0510, 2022.

Székely, G. J., Rizzo, M. L., and Bakirov, N. K. Measuring and testing dependence by correlation of distances. The Annals of Statistics, 35:2769–2794, 2007.

Tantardini, M., Ieva, F., Tajoli, L., and Piccardi, C. Comparing methods for comparing networks. Scientific reports, 9(1):17557, 2019.

Thrun, M. C., Märte, J., and Stier, Q. Analyzing quality measurements for dimensionality reduction. Machine Learning and Knowledge Extraction, 5(3):1076–1118, 2023.

Tiefelsdorf, M. and Boots, B. The exact distribution of moran’s i. Environment and Planning A, 27(6):985–999, 1995.

Tong, F., Lasocki, S., Orlecka-Sikora, B., et al. Nonparametric kernel density estimation of magnitude distribution for the analysis of seismic hazard posed by anthropogenic seismicity. arXiv preprint 2503.04393, 2025.

Van den Berge, K., Roux de Bézieux, H., Street, K., Saelens, W., Cannoodt, R., Saeys, Y., Dudoit, S., and Clement, L. Trajectory-based differential expression analysis for single-cell sequencing data. Nature communications, 11 (1):1201, 2020.

van den Heuvel, E. and Zhan, Z. Myths about linear and monotonic associations: Pearson’sr, spearman’s ρ, and kendall’s τ. The American Statistician, 76(1):44–52, 2022.

Vathy-Fogarassy, A. and Abonyi, J. Local and global mappings of topology representing networks. Information Sciences, 179(21):3791–3803, 2009.

Venna, J. and Kaski, S. Neighborhood preservation in nonlinear projection methods: An experimental study. In International conference on artificial neural networks, pp. 485–491. Springer, 2001.

Vinh, N., Epps, J., and Bailey, J. Information theoretic measures for clusterings comparison: Variants, properties, normalization and correction for chance. Journal of Machine Learning Research, 11:2837–2854, 10 2010.

Wang, H., Tang, Q., and Zheng, W. L1-norm-based common spatial patterns. IEEE Transactions on Biomedical Engineering, 59(3):653–662, 2011.

Watts, D. J. and Strogatz, S. H. Collective dynamics of ‘small-world’networks. nature, 393(6684):440–442, 1998.

Weglarczyk, S. Kernel density estimation and its application. In ITM web of conferences, volume 23, pp. 00037. EDP Sciences, 2018.

Weiler, P., Lange, M., Klein, M., Pe’er, D., and Theis, F. Cellrank 2: unified fate mapping in multiview single-cell data. Nature Methods, 21(7):1196–1205, 2024.

Wolf, F. A., Angerer, P., and Theis, F. J. Scanpy: largescale single-cell gene expression data analysis. Genome biology, 19:1–5, 2018.

Wolf, F. A., Hamey, F. K., Plass, M., Solana, J., Dahlin, J. S., Göttgens, B., Rajewsky, N., Simon, L., and Theis, F. J. Paga: graph abstraction reconciles clustering with trajectory inference through a topology preserving map of single cells. Genome biology, 20:1–9, 2019.

Wu, R. and Keogh, E. J. Fastdtw is approximate and generally slower than the algorithm it approximates. IEEE Transactions on Knowledge and Data Engineering, 34 (8):3779–3785, 2020.

Xu, C., Lopez, R., Mehlman, E., Regier, J., Jordan, M. I., and Yosef, N. Probabilistic harmonization and annotation of single-cell transcriptomics data with deep generative models. Molecular systems biology, 17(1):e9620, 2021.

Xu, K., Hu, W., Leskovec, J., and Jegelka, S. How powerful are graph neural networks? arXiv preprint 1810.00826, 2018.

Yaveroğlu, Ö.N., Malod-Dognin, N., Davis, D., Levnajic, Z., Janjic, V., Karapandza, R., Stojmirovic, A., and Pržulj, N. Revealing the hidden language of complex networks. Scientific reports, 4(1):4547, 2014.

Yoo, A. B., Jette, M. A., and Grondona, M. Slurm: Simple linux utility for resource management. In Workshop on job scheduling strategies for parallel processing, pp. 44– Springer, 2003.

Zandieh, A., Han, I., Daliri, M., and Karbasi, A. Kdeformer: Accelerating transformers via kernel density estimation. In International Conference on Machine Learning, pp. 40605–40623. PMLR, 2023.

Zhao, J. and Itti, L. shapedtw: Shape dynamic time warping. Pattern Recognition, 74:171–184, 2018.

